# Evolutionary assembly of cooperating cell types in an animal chemical defense system

**DOI:** 10.1101/2021.05.13.444042

**Authors:** Adrian Brückner, Jean M. Badroos, Robert W. Learsch, Mina Yousefelahiyeh, Sheila A. Kitchen, Joseph Parker

## Abstract

A long-standing challenge in biology is explaining how the functions of multicellular organs emerge from the underlying evolution of cell types. We deconstructed evolution of an organ novelty: a rove beetle gland that secretes a defensive cocktail. We show that gland function was pieced together via assembly of two cell types that manufacture distinct compounds. One cell type forms a chemical reservoir in the beetle’s abdomen and produces alkane and ester compounds. We demonstrate that this cell type is a hybrid of cuticle cells and ancient pheromone and adipocyte-like cells, and executes its function via a mosaic of enzymes sourced from each parental cell type. The second cell type synthesizes noxious benzoquinones using a chimeric pathway derived from conserved cellular energy and cuticle formation pathways. We present evidence that evolution of each cell type was shaped by coevolution between the two cell types: the benzoquinones produced by the second cell type dissolve in solvents produced by the first, yielding a potent secretion that confers adaptive value onto the gland as a whole. Our findings illustrate how cooperation between cell types can arise, generating new, organ-level behaviors.

## Introduction

Animal phylogeny has been shaped by the evolution of new organs with novel functional properties (Raff, 1996). Structures such as eyes, skeletons and brains were multicellular innovations that opened up new ecological frontiers and catalyzed major diversification (Erwin, 2021; Hunter, 1998). How the functions of complex organs originate has challenged biologists since Darwin (Darwin, 1859; Dawkins, 1986; Lynch, 2007; Oakley and Speiser, 2012; Shubin et al., 2009; West-Eberhard, 2003). Organs are composed of specialized cell types that perform dedicated roles within these structures (Schmidt-Rhaesa, 2007); yet, knowledge of how cell types are constructed at the molecular level during evolution, and how they cooperate to produce emergent, organ-level behaviors, remains fragmentary. At the gene regulatory level, new cell types are thought to arise from the evolution of unique, combinatorial transcription factor activities that specify novel cell identities (Arendt et al., 2016). However, a major problem is explaining how the downstream molecular components, which carry out the function of the differentiated cell, themselves originate. For a cell to execute its role, multiple gene products must be configured to work in concert. In a multicellular organ, this process must be coordinated across multiple cell types, fostering a division of labor that generates organ behavior (Kishi and Parker, 2021).

The molecular changes leading to new cell type- and organ-level functions are obscured by the deep antiquity of many animal organs (Griffith and Wagner, 2017; Shubin et al., 2009). However, insight may be gained from ‘evolutionary novelties’—lineage-specific structures that are absent in outgroups or serially homologous body parts (Erwin, 2015; Wagner and Lynch, 2010). Vertebrate eyes (Lamb et al., 2007), insect wings (Linz and Tomoyasu, 2018), beetle horns (Hu et al., 2019) and the mammalian placenta (Griffith and Wagner, 2017) are paradigms that connect genomic and developmental changes to the establishment of new phenotypes. Nevertheless, knowledge of how the functions of novel organs arise at the cellular level has to date been limited. One hurdle has been the challenge of measuring the transcriptomic composition of cell types— critical for tracing how different subfunctions within a cell evolved. The advent of single cell methods has now brought many questions regarding cell type evolution within reach (Marioni and Arendt, 2016). New tools permit decomposition of the transcriptome, enabling discovery of gene expression programs underlying cell identity, and quantitative assessment of relationships between cell types (Way and Greene, 2019). Here, we leverage these methods to trace the evolution of cooperating cell types in an animal organ.

Exocrine glands are archetypal organ novelties: unique, clade-specific glands have evolved convergently thousands of times in the Metazoa, transforming how animals interact via pheromonal communication and chemical defense, enabling specialized modes of prey capture and digestion, and aiding in substrate adhesion, desiccation avoidance, and antimicrobial protection (Brückner and Parker, 2020). Each origin of a novel gland involves the evolution of new secretory cell types that synthesize natural products (Torres and Schmidt, 2019). Venom glands of wasps, scorpions and cone snails, the explosive weaponry of bombardier beetles, and diverse mammalian scent glands are multicellular innovations composed of taxon-restricted cell types found nowhere else. For many such organs, including vertebrate salivary glands (Roussa, 2010), moth silk glands (Suzuki et al., 1990), cnidarian digestive glands (Babonis et al., 2019), and venom glands of many species (Surm and Moran, 2021), the final secretion is a cocktail produced by distinct secretory cell types. Biosynthetic synergism between cell types is pronounced in many small molecule-based chemical defense systems (Eisner et al., 2005a). Here, an inactive toxin and its activating enzyme may be produced by separate glands (Bourguignon et al., 2016; Eisner and Meinwald, 1966; Kuwahara et al., 2011). Alternatively, a soluble toxin and its solvent are secreted into a common reservoir, yielding a bioactive secretion (Francke and Dettner, 2005; Noirot et al., 1974). Such functional interdependence encapsulates how emergent, organ-level behaviors can arise from cooperation between cell types.

One animal clade is notable for pervasive glandular innovation. Rove beetles (Staphylinidae) are a radiation of 64,000 predominantly soil-dwelling predators (Betz et al., 2018). The beetles exhibit widespread evolution of abdominal glands that produce diverse compounds, including quinones, terpenes, alkaloids, iridoids, hydrocarbons and esters (Francke and Dettner, 2005). The unique, flexible body enables rove beetles to target secretions at other organisms, fostering interspecies relationships from chemical defense to symbiosis (Parker, 2016). The glands are often composite, multicellular organs that produce multi-compound secretions via distinct cell types (Dettner, 1993a). Although their chemistry and anatomy have been studied in many species (Francke and Dettner, 2005), how these organs function at the molecular level, and how they evolved in specific lineages, is unknown. Here, we examine the molecular architecture of one such gland, enabling us to infer how a novel organ and its constituent cell types evolved. We exploited the model species *Dalotia coriaria*, a member of the megadiverse Aleocharinae subfamily that possess a defensive ‘tergal gland’ on the dorsal abdomen (Steidle and Dettner, 1993). Using single cell transcriptomics of *Dalotia*’s abdominal segments, we uncovered novel pathways for defensive compounds in tergal gland cell types. By inferring transcriptomic and pathway relationships with more ancient insect cell types, we retrace how the tergal gland was functionally assembled during evolution. We present evidence that evolution followed an accessible path towards cell type cooperativity, building an organ capable of producing a multi-compound secretion that confers adaptive value.

## Results

### The tergal gland is an organ novelty

When provoked, *Dalotia* flexes its abdomen to release the contents of a large chemical defense gland—the tergal gland—positioned between abdominal segments A6 and A7 (Fig 1A, B) (Brand et al., 1973; Dettner, 1993b). The secretion is smeared directly onto the perceived threat and contains three benzoquinones (BQs) (Fig. 1D)—aromatics that are common arthropod defensive compounds (Blum, 1981; Eisner et al., 2005b; Pasteels and Grégoire, 1983; Wagner et al., 2020). BQs are topical irritants that bind TRPA1 channels (Ibarra and Blair, 2013), activating nociceptive pathways (Kang et al., 2010). The adaptive value of *Dalotia*’s secretion is demonstrated by experimentally depleting the contents of the gland, disarming the beetle and compromising its defense against predatory ants (Fig. 1C). The BQs may also have antimicrobial properties (Carcamo-Noriega et al., 2019; Dettner, 1993b; Lana et al., 2006).

**Figure 1.**
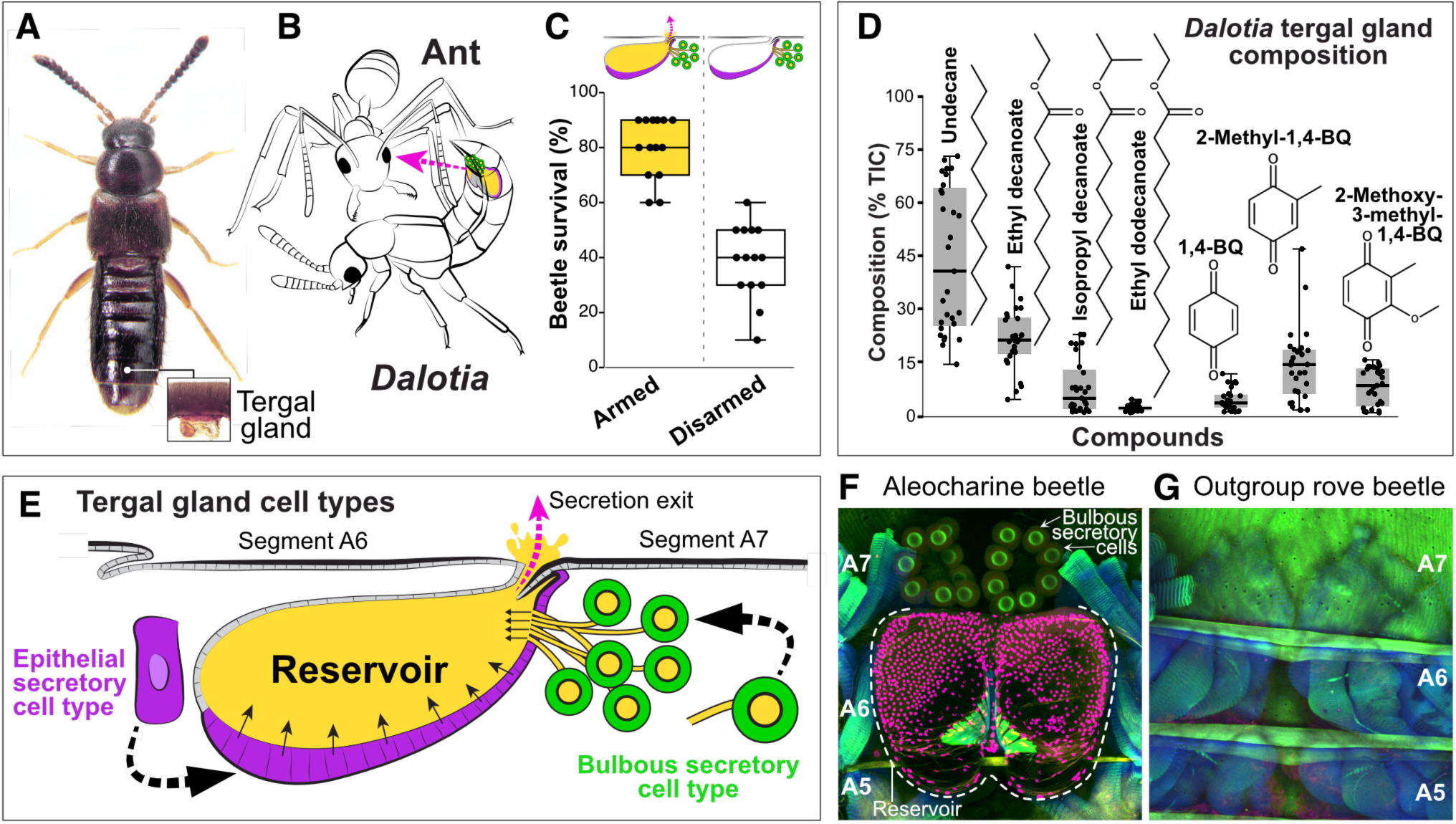
The tergal gland. **A:** *Dalotia coriaria* showing position of tergal gland. **B:** Cartoon of gland deployment in an ant encounter. **C:** Percentage *Dalotia* survival in arenas with predatory ants (*Liometopum occidentale*) after 48 h (armed versus disarmed tergal glands; Mann-Whitney U test: p< 0.001). **D:** Chemical composition of *Dalotia*’s tergal gland secretion (n = 29 beetles)**. E:** Cartoon cross section of abdominal showing tergal gland cell types: a secretory epithelium (magenta) invaginated between segments A6-A7 forming a chemical reservoir, and bulbous secretory cells (green) that connect to the reservoir via ducts. When provoked, the secretion is released via exit between segments. **F:** Confocal image of internal face of segments A5-A7 of aleocharine abdomen (magenta: Engrailed antibody labelling nuclei of reservoir cells; green: autofluorescence revealing bulbous secretory cells; blue: phalloidin-stained muscle). **G**: Confocal image of internal face of segments A5-A7 of outgroup rove beetle (Tachyporinae: *Coproporus*). No gland is present.

Although BQs are highly noxious, each compound in isolation is a solid. Consequently, the secretion also contains four less toxic compounds: large amounts of a C11 alkane, n-undecane, low amounts of two C10 esters, ethyl decanoate and isopropyl decanoate, and trace levels of a C12 ester, ethyl dodecanoate (Fig. 1D). These compounds may function as solvents, combining with BQs to create a potent secretion, as well as promoting spreading and penetration into cuticles (Blum, 1981; Dettner, 1984). The secretion is thus a cocktail, the composition of which may be explained by possible synergism between component parts. To produce this cocktail, the tergal gland consists of two anatomically distinct cell types: i) a secretory epithelium, composed of columnar cells that are continuous with the epidermis and form a large reservoir inside the body into which they secrete directly (Fig. 1E, F), and ii) large, bulbous secretory cells, 10–14 per animal, located posterior to the reservoir into which they feed via tubular ducts (Fig. 1E, F). The tergal gland, and these two cell types, are unique to aleocharines (Steidle and Dettner, 1993); they are absent in outgroup rove beetles, which cannot produce defensive compounds (Fig. 1G, Fig. S1). We sought to determine how these two cell types cooperate to synthesize *Dalotia*’s defensive secretion.

### The solvent cell type

*Dalotia*’s ability to produce defensive compounds is refractory to antibiotics (Fig. S2A-E). We therefore reasoned that biosynthesis relies not on symbiotic bacteria, as in some other chemically-defended rove beetles (Piel, 2002), but on enzymes encoded in the beetle’s genome and expressed within the gland. We created type-specific transcriptomes, recovering candidate enzymes expressed in each cell type (Fig. S3A, Fig. S4, Fig. S5). We focused initially on synthesis of the undecane and esters hypothesizing that their hydrocarbon chains derive from fatty acids (Stanley-Samuelson et al., 1988). Fatty acids are built from units of acetyl-CoA and malonyl-CoA produced by glycolysis (Wakil et al., 1983). Rearing *Dalotia* on D-glucose-^13^C_6_ food, we observed strong ^13^C incorporation into the undecane and esters (Fig. S6A), confirming that they originate *de novo* via the glycolytic pathway and fatty acid synthesis. In contrast, adding D_23_-dodecanoic acid to *Dalotia*’s food led to negligible deuterium incorporation (Fig. S6B). The compounds thus do not derive from consumed fatty acids.

In animals, the multistep reaction to produce fatty acids is catalyzed by a multidomain fatty acid synthase (FASN) (Smith, 1994; Smith et al., 2003). Animal FASNs are generally known to synthesize the commonest fatty acids, C16 and C18 in length (Smith et al., 2003). However, the gland compounds in *Dalotia* are shorter, C10-C12 in length. Notably, a predicted *FASN* was upregulated in the epithelial secretory cells comprising the reservoir (Fig. S4), but not in the bulbous secretory cells (Fig. S5). Strong expression in epithelial secretory cells was confirmed via *in situ* hybridization chain reaction (HCR) (Choi et al., 2018) (Fig. 2F). To deduce this *FASN*’s function, we temporally silenced its expression with systemic RNA interference (RNAi) (Parker et al., 2017; Tomoyasu and Denell, 2004) (Fig. S3B). Silencing causing near-total loss of undecane and all three esters without impacting the BQs (Fig. 2A, B, Fig. S7A). We name this enzyme ‘Master Fatty Acid Synthase’ (MFASN), due to its upstream role in synthesizing fatty acid-derived compounds in the tergal gland. We infer that the epithelial secretory cells (magenta cell type in Fig. 2J) are the source of these compounds. This cell type is hence named the ‘solvent cell type’.

**Figure 2:**
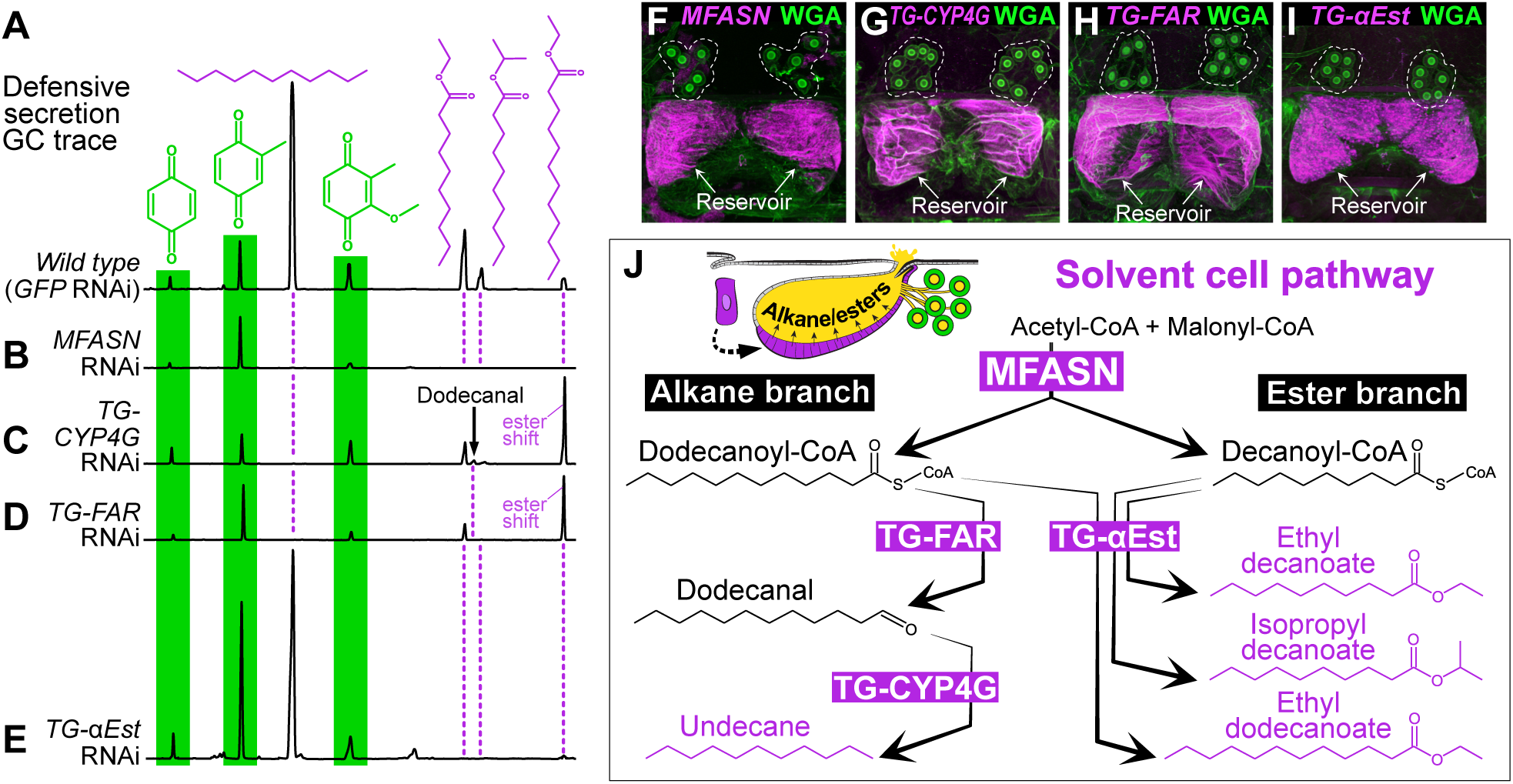
Identification of the solvent cell type and its biosynthetic pathway. **A–E:** Gas chromatograph traces of *Dalotia* tergal gland secretions **A:** wild type secretion showing peaks corresponding to each defensive compound. **B:** RNAi silencing of *MFASN* causes specific loss of the undecane and esters without affecting the BQs. **C:** Silencing *TG-CYP4G* causes loss of the undecane and the novel appearance of its dodecanal precursor. A strong increase in the C12 ester also occurs. **D:** Silencing *TG-FAR* causes loss of undecane without appearance of dodecanal. As with *TG-CYP4G* knockdown, the C12 ester level increases. **E:** Silencing *TG-αEst* causes selective of the three esters without affecting levels of undecane or the three BQ species. Statistical analyses related to the RNAi experiments (**B-E**) can be found in supplementary Table S1. **F–I:** Hybridization Chain Reaction (magenta) and Wheat Germ Agglutinin (green) labelling of tergal glands reveals expression of all solvent pathway enzymes in epithelial secretory cells comprising the reservoir: *MFASN* (**F**), *TG-CYP4G* (**G**)*, TG-FAR* (**H**)*, TG-αEst* (**I**). **J:** Diagram of the solvent pathway leading to synthesis of the undecane and esters in *Dalotia*’s tergal gland secretion.

**Table 1.**
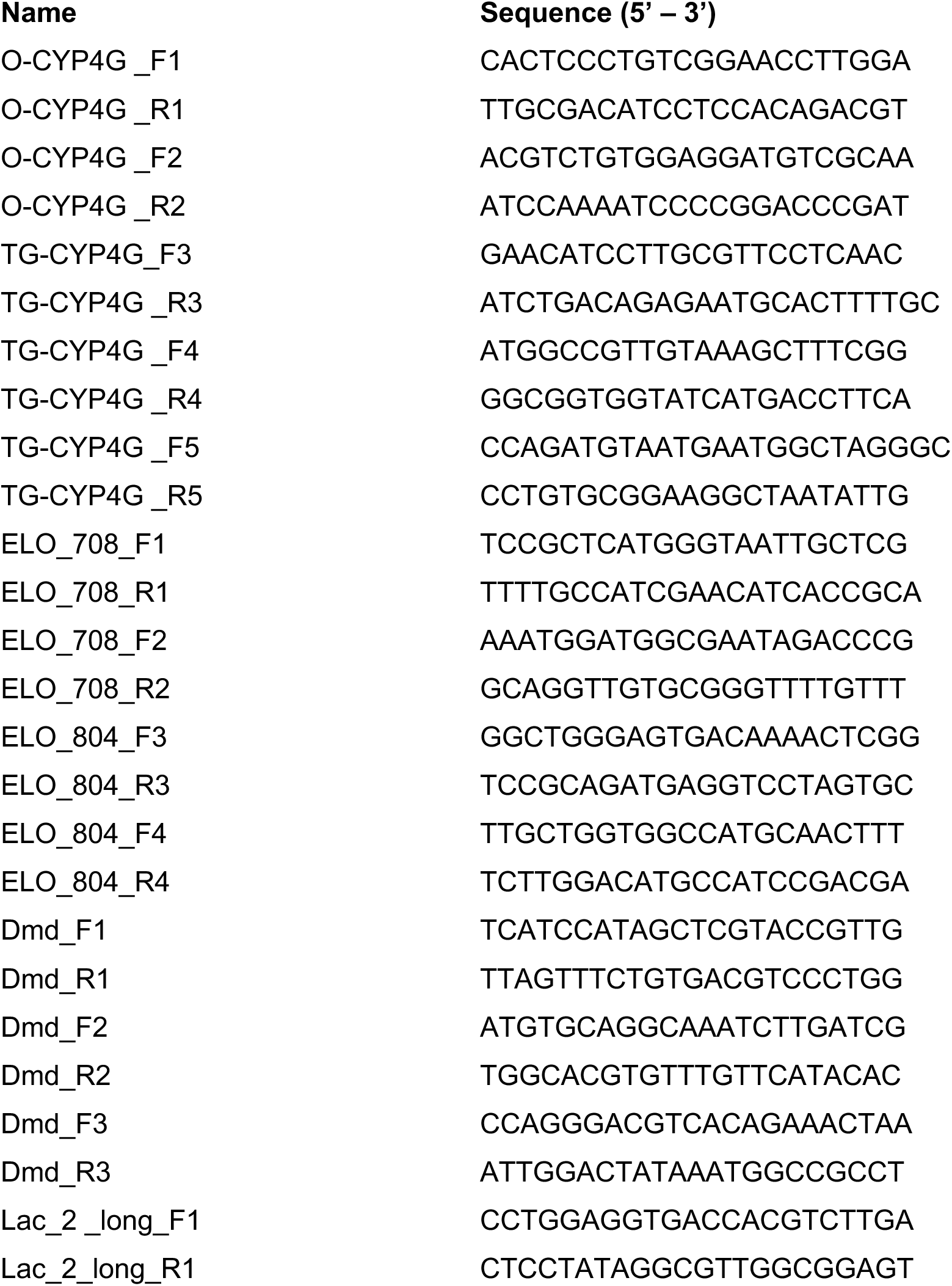
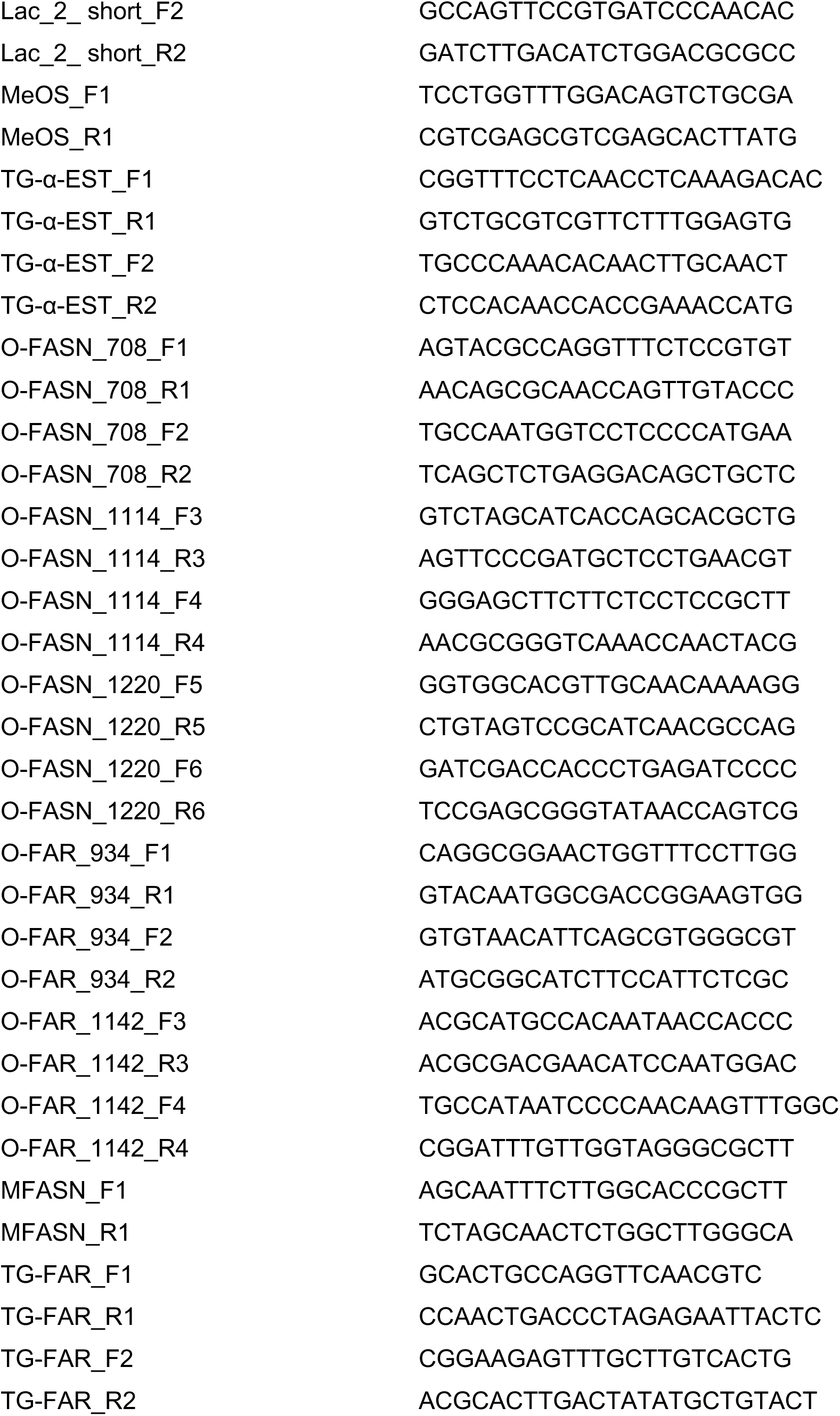
RNAi targets and their primer sequences.

Further analysis of the solvent cell transcriptome led to discovery of a complete fatty acid pathway downstream of MFASN. The maximally expressed transcript in solvent cells is a predicted cytochrome P450 (CYP) (Fig. S4), cell type-specific expression of which was confirmed by HCR (Fig. 2C). Phylogenetic analysis revealed that this CYP belongs to an insect-specific ‘CYP4G’ class (Fig. S7) that are known to decarbonylate aldehydes to create alkanes (Feyereisen, 2020; Qiu et al., 2012). Consistent with this function, silencing the tergal gland *CYP4G* (herein ‘*TG-CYP4G*’) caused selective loss of the undecane and the novel appearance of its C12 aldehyde precursor, dodecanal (Fig. 2C, Fig. S7B). We infer that TG-CYP4G functions downstream of MFASN as a terminal enzyme that decarbonylates dodecanal to make undecane. To produce the dodecanal intermediate, the activated C12 fatty acid (dodecanoyl-CoA) produced by MFASN must first be reduced to an aldehyde. Conspicuously, a predicted fatty acyl-CoA reductase (herein ‘*TG-FAR*’) was strongly expressed in solvent cells (Fig. S4, Fig. 2H). As with *TG-CYP4G*, silencing *TG-FAR* caused specific loss of undecane, but this time without the appearance of dodecanal in the secretion (Fig. 2D, Fig. S7C). We infer that solvent cells make undecane via an alkane pathway in which MFASN produces dodecanoic acid that is reduced by TG-FAR and decarbonylated by TG-CYP4G (Fig. 2J).

In addition to this alkane pathway, a second branch of the solvent cell fatty acid pathway yields the esters. Silencing *MFASN* caused loss of the three esters (Fig. 2B). Because two of these are C10 compounds (isopropyl and ethyl decanoate), MFASN must produce their C10 fatty acid precursor, decanoic acid, in addition to the dodecanoic acid precursor of undecane (Fig. 2L). Notably, we observed strong upregulation of a single α-esterase enzyme in solvent cells (Fig. S4). Silencing this enzyme (‘*TG-αEst*’) removed all three esters without affecting undecane (Fig. 2E, Fig. S7D). TG-αEst is thus the key enzyme in the ester branch of the solvent pathway. TG-αEst esterifies both C10-CoA and C12-CoA to yield the three esters (Fig. 2J).

### A cell atlas of *Dalotia* abdominal segments

The solvent cells create a volatile solution that we hypothesize dissolves the BQs. To investigate how this function evolved, we attempted to reconstruct the evolutionary sequence by which the solvent cells were pieced together. We applied single cell RNA sequencing (scRNAseq) using droplet microfluidics (10x Chromium, version 3) to entire adult abdominal segments, enabling us to determine the transcriptomic relationships of the solvent cells to other cell types within the *Dalotia* abdomen. We created two segmental atlases: ‘Segment 7’ comprising segment A7 plus the complete solvent cell reservoir (Fig. 3A); and ‘Segment 6’ encompassing segment A6 minus the solvent cell reservoir. Segment 6 is serially homologous but lacks the gland, hence approximating ancestral segment morphology (Fig. 3A, B). We integrated five 10x runs (Fig. S9A-C) and selected 3000 transcripts that capture transcriptional heterogeneity across cells for subsequent analyses. From the two abdominal atlases, we determined that 26 cell clusters provided biologically meaningful resolution for studying solvent cell relationships (Fig. 3C).

**Figure 3:**
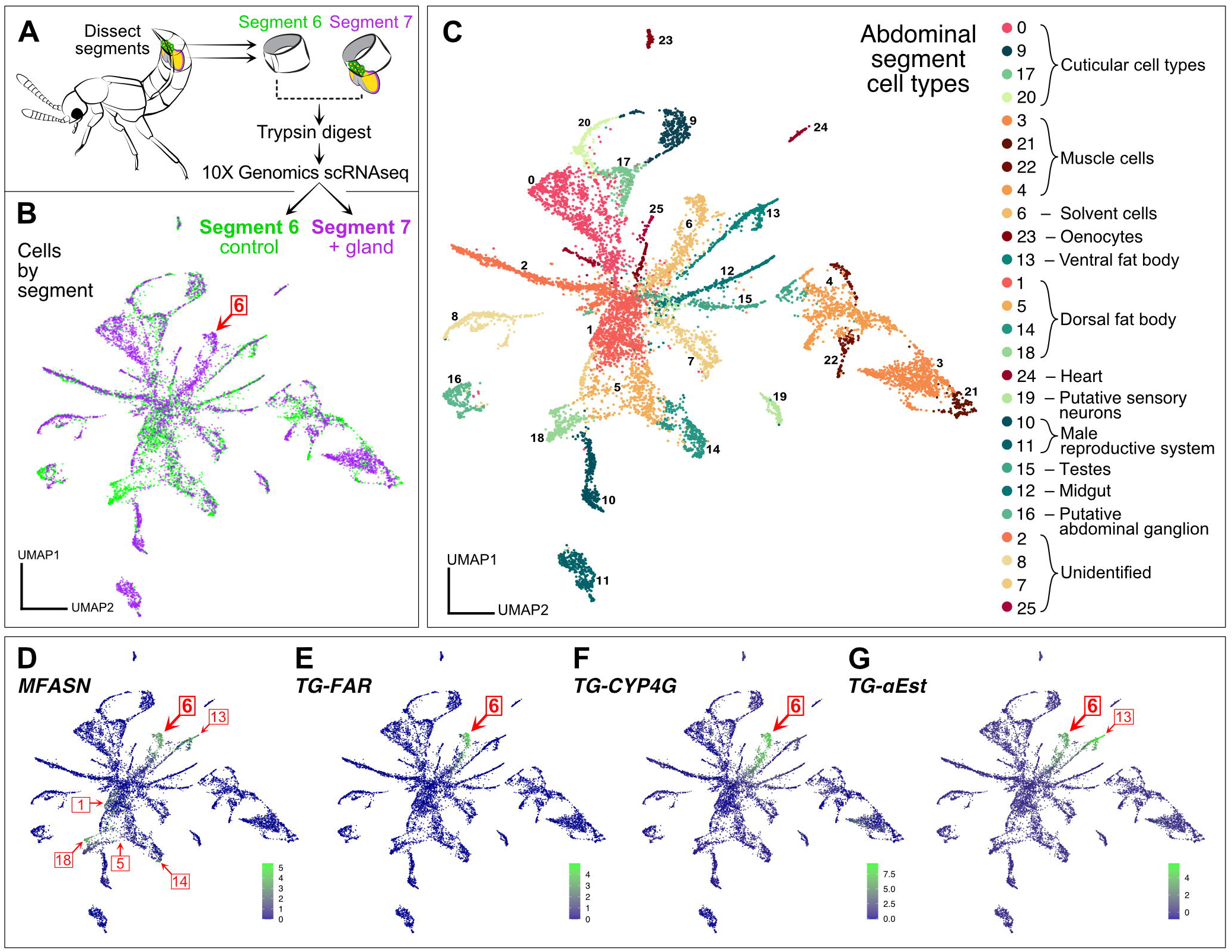
Single cell atlas of *Dalotia* abdominal segments. **A:** Scheme for 10x scRNAseq (for details see materials and methods). **B:** UMAP of cells separated by segment (green: Segment 6; magenta: Segment 7). Cell type 6—the solvent cells—is indicated, and composed exclusively of Segment 7 cells. **C:** UMAP of the 26 cell types and groupings into higher-order cell classes. **D–G:** Expression of solvent pathway enzymes: *MFASN* (**D**), *TG-FAR* (**E**), *TG-CYP4G* (**F**), *TG-αEst* (**G**). Numbers shown in the UMAPs correspond to cell types in C.

We annotated cell clusters and evaluated hierarchical relationships using each cell type’s mean expression of the 3000 gene set, producing a tree of transcriptomic similarity (Fig. S9D). We used this topology to evaluate whether the 26 cell types could be further grouped into higher-order classes via a random forest classifier (Breiman, 2001) (Fig. S9E). This classifier produced strong support for 14 higher groupings, referred to as ‘cell classes’ (Fig. 3C, Fig. S9D-F). Overall, 85% of all cells were correctly assigned to one cell class (Fig. S9E). The 26 cell types and 14 cell classes thus represent a robust classification of cell types in abdominal segments 6 and 7.

### Ester branch evolution via co-option of fat body enzymes

Within the abdominal cell atlas, a single cluster co-expressed the four solvent pathway enzymes (Fig. 3D-G; Fig. S10A–G). This cell type—cluster number 6—is thus the solvent cell type. This determination is further supported by cell type 6 being exclusively restricted to Segment 7 (Fig. 3B). To investigate solvent pathway evolution, we searched for features shared with other cell types. We observed that several cell types—1, 4, 5, 13 and 18—expressed *MFASN* to high levels, indicating fatty acid synthesis (Fig. 3D). We deduced that these cell types comprise *Dalotia*’s fat body—an ancient insect organ sharing many functions with vertebrate liver and adipose tissue (Arrese and Soulages, 2010). Fat body cells are sites of fat and glucose storage, as well as lipid biosynthesis, and are identifiable as lipid droplet-containing cells forming loose tissue that lines body segments and surrounds most organs. HCR-labelling revealed *MFASN* expression throughout *Dalotia*’s fat body (Fig. S11A, B). Phylogenetic analysis shows that *MFASN* is a single copy gene present in all beetles, as well as in *Drosophila* (Fig. 4L). *MFASN*’s deep conservation and broad expression across *Dalotia*’s fat body cell types imply an ancestral function in the insect fat body. We HCR-labelled the orthologous transcript in a distantly-related outgroup beetle, *Tribolium* (Tenebrionidae), revealing strong, fat body-specific expression (Fig. S11C). *MFASN*’s ancestral function was indeed in the fat body; production of C12 and C10 precursors thus evolved by co-opting *MFASN* into solvent cells.

**Figure 4:**
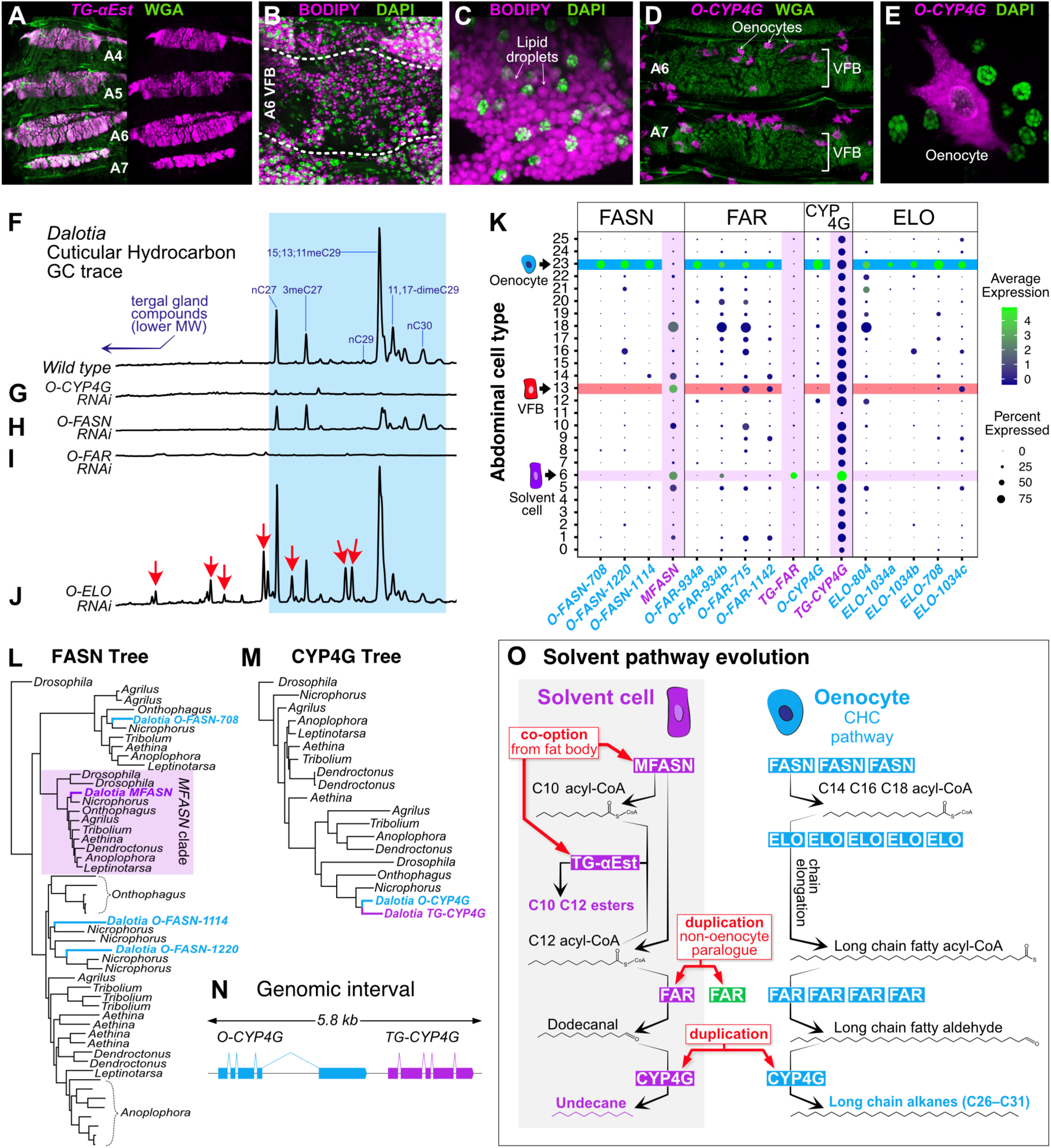
Solvent pathway evolution. **A:** Expression of *TG-αEst* (magenta: HCR) in segmental bands of VFB (green: Wheat Germ Agglutinin-stained membranes). **B**, **C**: BODIPY (magenta) labels lipid droplets in VFB cells (green: DAPI-stained nuclei). **D, E:** HCR of *O-CYP4G* (magenta) stains oenocytes in ventral abdomen. **D:** Intermingling of oenocytes and VFB (green: WGA). **E:** Single oenocyte with *O-CYP4G* expression (green: DAPI). **F**-**J**: GC traces of *Dalotia* CHCs. **F:** wild type with identities of major peaks indicated. **G:** Silencing *O-CYP4G* causes loss of CHCs. **H:** Silencing one of three oenocyte *O-FASN*s causes loss of most CHCs. **I:** Silencing one of three *O-FAR*s causes loss of CHCs. **J:** Silencing one of five *O-ELO*s shifts CHC profile to shorter chain lengths. **K**: Transcript expression dot plot of *FASN*, *FAR*, *CYP4G* and *ELO* gene family members across abdominal cell types. Dot size corresponds to the percentage of cells of each cell type expressing the transcript; color represents mean expression level. **L**-**M**: Maximum-likelihood trees of FASN (**L**) and CYP4G (**M**) families in genomes of *Dalotia*, other beetles and *Drosophila*. **N**: *Dalotia* genomic interval showing *TG-CYP4G* and *O-CYP4G* tandem copies. **O**: Model of solvent pathway assembly via co-option and duplication of VFB and oenocyte enzymes.

Adipocyte-like fat body cells can be highly heterogenous in insects (Haunerland and Shirk, 1995), and this is so in *Dalotia*. First, cell types 1, 5, 14 and 18—but not 13—cluster distinctly based on mean transcript expression (Fig. S9D), implying two different fat body classes. The four former cell types also express *Drosophila* fat body markers including *pumpless* and *apolipophorin*, whereas cell type 13 does not (Fig. S11D-F). Most strikingly, the two fat body classes differ in expression of a *Dalotia*-specific expansion of α*-*esterase enzymes (Fig. S12A-E). Early-branching members of this clade are expressed in cell types 5, 14 and 18 (Fig. S12B, C, E), whereas cell type 13 expresses a single, distal-branching copy—*TG-αEst*—shared exclusively with solvent cells (Fig. 3G, Fig. S12D). Using HCR of *TG-αEst*, we identified cell type 13 as subepidermal bands of fat body containing large lipid droplets (Fig. 4A-C). We refer to cell type 13 as ‘Ventral Fat Body (VFB)’, and other fat body types distributed more dorsally as ‘Dorsal Fat Body’ (DFB; consisting of cell types 1, 5, 14 and 18) (Fig. S9D, Fig. S11D). Given that the ancestral function of this α*-*esterase clade is likely in the fat body, and *TG-αEst* itself functions in the VFB specifically, we infer that the ester branch of the solvent pathway evolved by recruiting a fat body α*-*esterase to function downstream of MFASN (Fig. 4O).

### Alkane branch evolution via duplication of oenocyte enzymes

Many aleocharines are known to synthesize esters as the putative BQ solvent, and lack alkanes which may represent an evolutionarily more recent addition (Steidle and Dettner, 1993). Recruitment of MFASN and a fat body α*-*esterase may thus have been an early step in solvent pathway evolution, with the alkane branch added subsequently. We noted that one cell type— 23—expressed a paralogue of *TG-CYP4G* (Fig. 4K, Fig. S13A). CYP4Gs have been shown to function canonically in oenocytes—ancient pheromone-producing cells present in all insects (Makki et al., 2014). Oenocytes synthesize very long-chain (C25–C40) alkanes and alkenes known as cuticular hydrocarbons (CHCs), which are secreted onto the body and encode a chemical signature of species identity (Billeter et al., 2009). CHCs also form a waxy barrier that guards against desiccation (Koto et al., 2019; Qiu et al., 2012). In oenocytes, CYP4G performs terminal decarbonylation to yield the secreted CHCs (Feyereisen, 2020; Qiu et al., 2012). We HCR-labelled this *CYP4G*, revealing clusters of enlarged cells subepidermally in the ventral abdomen, often intermingled with VFB (Fig 4D, E). Silencing this locus led to near-total loss of *Dalotia*’s CHCs (Fig 4F, G; Figure S13E), confirming that these cells are the oenocytes. We name this enzyme *O-CYP4G* (*Oenocyte-CYP4G*).

Recent studies have delineated a conserved oenocyte CHC pathway (Blomquist and Ginzel, 2021). This pathway bears striking similarity to the alkane branch of the solvent pathway: oenocytes use FASNs to produce fatty acids that are reduced to aldehydes by FARs before decarbonylation by CYP4G (Blomquist and Ginzel, 2021; Holze et al., 2020)—modifications that mirror exactly the steps for undecane synthesis (Fig. 2J). Consistent with the deep conservation of the insect CHC pathway, we find that, in addition to *O-CYP4G*, cell type 23 expresses three *FASN* paralogues and four *FAR*s (Fig 4K, Fig S13B-C), As with *O-CYP4G*, silencing selected copies of these enzymes strongly diminished CHC production (Fig 4H, I, Fig. S13F, G). Hence, *Dalotia* expresses parallel alkane pathways in ancient oenocytes and novel solvent cells, and their close similarity implies common ancestry. Indeed, we find *TG-CYP4G* and *O-CYP4G* are sister duplicates (Fig. 4M), the two copies sitting tandemly in the genome (Fig. 4N). The ancestral CYP4G was likely an oenocyte enzyme that synthesized CHCs; duplication gave rise to oenocyte and solvent cell copies (Fig. 4O). Whether duplication of an *O-FAR* led to *TG-FAR* is less clear; *FAR*s undergo extensive gene birth-and-death, so their history is challenging to infer (Finet et al., 2019). *TG-FAR* is not an unambiguous sister paralogue of an *O-FAR*, so its role in alkane synthesis may have arisen convergently (Fig. S14). Regardless, the CHC pathway represented a pre-existing template: by recruiting FAR and CYP4G enzymes downstream of MFASN, alkanes were added to the defensive secretion (Fig. 4O).

Despite similar enzyme logic, the products of the two pathways are markedly different: the CHC pathway produces very-long-chain waxy hydrocarbons, while the solvent pathway makes medium-chain volatile liquids. Oenocytes make longer compounds via elongases of very-long-chain fatty acids (ELOs) (Blomquist and Ginzel, 2021; Holze et al., 2020) (Fig. 4O). We find *Dalotia* oenocytes express five ELOs (Fig. 4K, Fig. S13D), knockdown of which causes an altered profile with many shorter chain compounds (Fig. 4J, Fig. S13I), without reducing total CHC levels (Fig. S13H). In contrast, solvent cells express no ELOs (Fig. 4K, Fig. S13D). Hence, the solvent pathway evolved via selective FAR and CYP4G recruitment, without an ELO, enabling medium-chain biosynthesis (Fig. 4O).

### Solvent cell evolution via transcriptomic hybridization of ancient cell types

Solvent cells comprise part of the beetle’s cuticle, forming part of the intersegmental membrane joining abdominal segments A6 and A7. Developmentally, the cells derive from the epidermal posterior (P) compartment of segment A6, and express Engrailed (Fig. 1F, Fig. S15A, B), the P compartment selector transcription factor (Morata and Lawrence, 1975). Like the surrounding epidermis, solvent cells produce chitin, which forms an internal lining to the reservoir and is continuous with the rest of the exoskeleton (Steidle and Dettner, 1993). Despite their epidermal identity, solvent cells are enlarged, columnar secretory cells that manufacture alkane and ester compounds. To understand how solvent cells acquired this property, we studied their transcriptomic relationship to other cell types within the abdomen. We employed consensus non-negative matrix factorization (cNMF) (Kotliar et al., 2019), performing an unsupervised search for constellations of significantly co-expressed transcripts across all cells within the 10x data set, irrespective of their cell type. cNMF applies iterative NMF treatments on a transcripts by transcripts matrix to identify groups of significantly co-expressed genes (’gene expression programs’ or ‘GEPs’). GEPs discretize the transcriptome into building blocks that may be surrogates of cellular properties. A GEP used in one or a few cell types may confer aspects of cell identity; conversely, a GEP used by many cell types likely underlies a routine activity such as mitosis (Kotliar et al., 2019). Employing cNMF, we determined that 20 GEPs accurately capture a decomposed representation of the total transcriptome of segments 6 and 7 (Fig. S16A-D). We calculated the proportional contribution of each GEP to each individual cell type’s transcriptome, depicted as a usage map (Fig. 5A). Visualizing transcriptome composition in this way, some GEPs appear cell type-specific, such as GEP8 and GEP2—unique identity GEPs expressed in heart and midgut cells, respectively. Conversely, most cell types express GEPs 18 and 12, implying common cellular activities.

**Figure 5.**
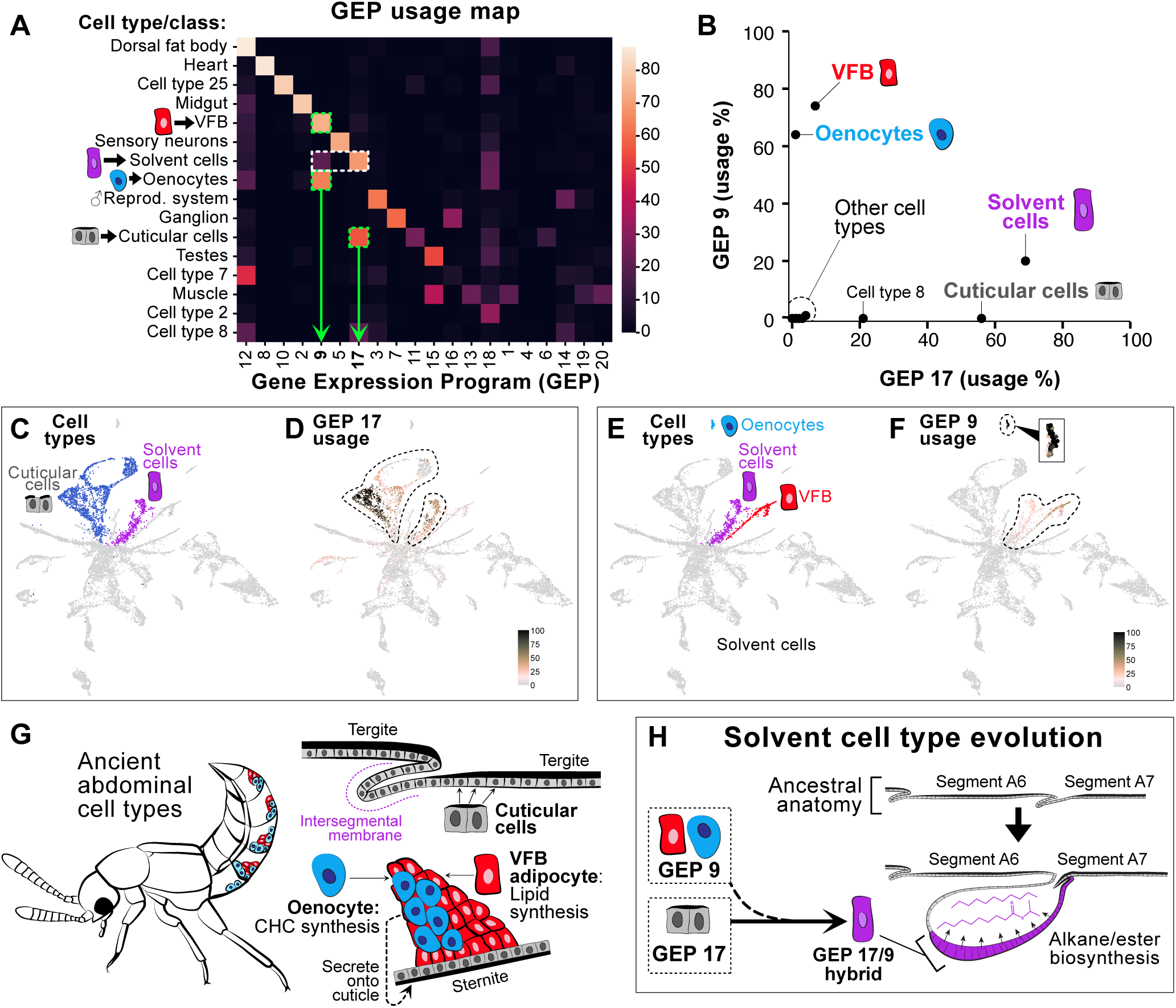
Solvent cell evolution. **A:** Usage map showing contribution of 20 GEPs to each cell type’s transcriptome. For clarity and to facilitate computation, some cell types were grouped together into cell classes. **B:** Percentage contribution of GEPs 9 and 17 to each cell type or class reveals the solvent cells are a hybrid of both GEPs. **C:** UMAP of abdominal cell types (from Fig. 3C), with the cuticle cell class and solvent cells highlighted. **D:** UMAP showing global usage of GEP 17 across all cell types. Dotted lines encircle the cuticle cell class and solvent cells. **E:** UMAP with oenocytes, VFB and solvent cells highlighted. **F:** UMAP of abdominal cell types showing global usage of GEP 9. Dotted lines encircle oenocytes, VFB and solvent cells (inset shows magnification of the oenocyte cluster). **G:** Cartoon of *Dalotia* showing segmentally repeated VFB and oenocyte cell types in ventral abdomen. **H;** Model of solvent cell type evolution: cuticle cells comprising the intersegmental membrane between tergites of segment A6 and A7 expresses GEP 17 as its principal GEP, but acquisition of GEP 9 from ancient oenocyte and fat body cell types transformed them into solvent cells capable of alkane and ester production.

Of particular interest, however, are cell types composed of combinations of GEPs (Fig. 5A). These include the solvent cells, recovered as a composite of GEPs 17 and 9 (Fig. 5A, B). Consistent with solvent cells being part of the cuticular epidermis, GEP 17 is the principal GEP expressed by the cuticular cell class (Fig. 5B-D). This cell class is composed of cell types 0, 9, 17 and 20 (Fig. 3C, Fig S9D), based on expression of multiple cuticle proteins and Laccase 2 that functions in cuticle tanning (Arakane et al., 2005) (Fig. 3B, Fig. 5A, B; Fig. S19C). However, solvent cells are clearly divergent from other cuticular cells in their additional expression of GEP 9, which accounts for ∼20% of the transcriptome (Fig. 5B, C). The composition of GEP 9 reveals that it is constituted by 61 core transcripts (applying a stringent z-score filter of < 0.002; Table S5) that show strong and significant enrichment for biological processes related to lipid metabolism and fatty acid biosynthesis (Fig. S17A-D). GEP 9 is thus a transcriptional module that may endow solvent cells with their capacity for high level fatty acid production and modification.

Remarkably, GEP 9 is the principal GEP of both oenocytes and VFB (Fig. 5A, B, E, F). The novel expression module in solvent cells therefore defines the two cell types from which solvent pathway enzymes were co-opted or duplicated. Unlike solvent cells, VFB and oenocytes show no pronounced GEP usage beyond GEP 9 (aside from GEPs 18 and 12 that most cell types express) (Fig. 5A). GEP 9 thus likely contributes to the functional identity of these two cell types, which are both specialized for fatty acid biosynthesis. It follows that GEP 9 probably imparts this same function in solvent cells. Close relationship between solvent cells, VFB and oenocytes is further supported by the three cell types forming a clade based on mean transcript expression (Fig. S9D), and random forest classifies them into the same cell class (Fig. S9E, F). That solvent cells are a novelty within the rove beetle’s cuticle, while oenocytes and fat body cells are ancient, non-cuticular cell types in all insects, implies an evolutionary scenario (Fig. 5G, H). We suggest that solvent cells arose via transcriptomic hybridization between cell types: they are a cuticular cell type, ancestrally comprising intersegmental membrane, which gained the expression module that evolved in oenocytes and fat body, thereby equipping them for high-level fatty acid biosynthesis (Fig. 5H). As part of this process, the oenocytes and fat body also contributed to distinct branches of the solvent pathway (Fig. 4O).

### The BQ cell type

We next focused on the mechanism of BQ synthesis. Despite widespread use of BQs in chemical defense (Blum, 1981), their mechanistic origins were hitherto unknown. Aromatic compounds in animals are often acquired from dietary aromatic amino acids or symbiotic microbes, but may sometimes be synthesized *de novo* (Brückner et al., 2020; Torres et al., 2020). To determine the synthetic route, we fed *Dalotia* D-glucose-^13^C_6_ and observed negligible ^13^C incorporation into the three BQs (Fig. S18), arguing against complete *de novo* synthesis. Conversely, feeding Tyr-^13^C_6_ or Phe-^13^C_6_ led to strong ^13^C incorporation, with molecular weights of all BQs increasing by exactly 6 (Fig. S18). The benzene rings of *Dalotia*’s BQs thus derive from dietary aromatic amino acids.

Beyond their use in chemical defensive, quinone compounds play key roles in insect metabolism: ubiquinone (coenzyme Q_10_) is a redox-active molecule synthesized in mitochondria where it functions in electron transport (Stefely and Pagliarini, 2017); additionally, quinone intermediates arise during exoskeleton maturation (cuticle tanning) (Noh et al., 2016). In both contexts, aromatic amino acids are the precursors. We asked whether these ancient pathways could give clues to BQ synthesis in the tergal gland. In cuticle tanning, oxidation of Tyr-derived dopa and dopamine creates quinones that are pigment precursors and protein crosslinkers used for cuticle hardening (sclerotization) (Noh et al., 2016). Oxidation is mediated by Laccase 2 (Lac2), a secreted multicopper oxidase (MCO) (Arakane et al., 2005; Asano et al., 2019). A predicted MCO was strongly upregulated in the second cell type within the gland—the bulbous secretory cells with ducts (Fig. S5). This transcript encodes a secreted protein with three cupredoxin domains in the same configuration as other laccases (Fig. 6A) (Dwivedi et al., 2011). Silencing this laccase caused near-total loss of all three BQs, without affecting the solvent compounds (Fig. 6B, Fig. S19A). We name this laccase ‘Decommissioned’ (Dmd) after its loss-of-function phenotype where the toxin is eliminated from the secretion.

**Figure 6.**
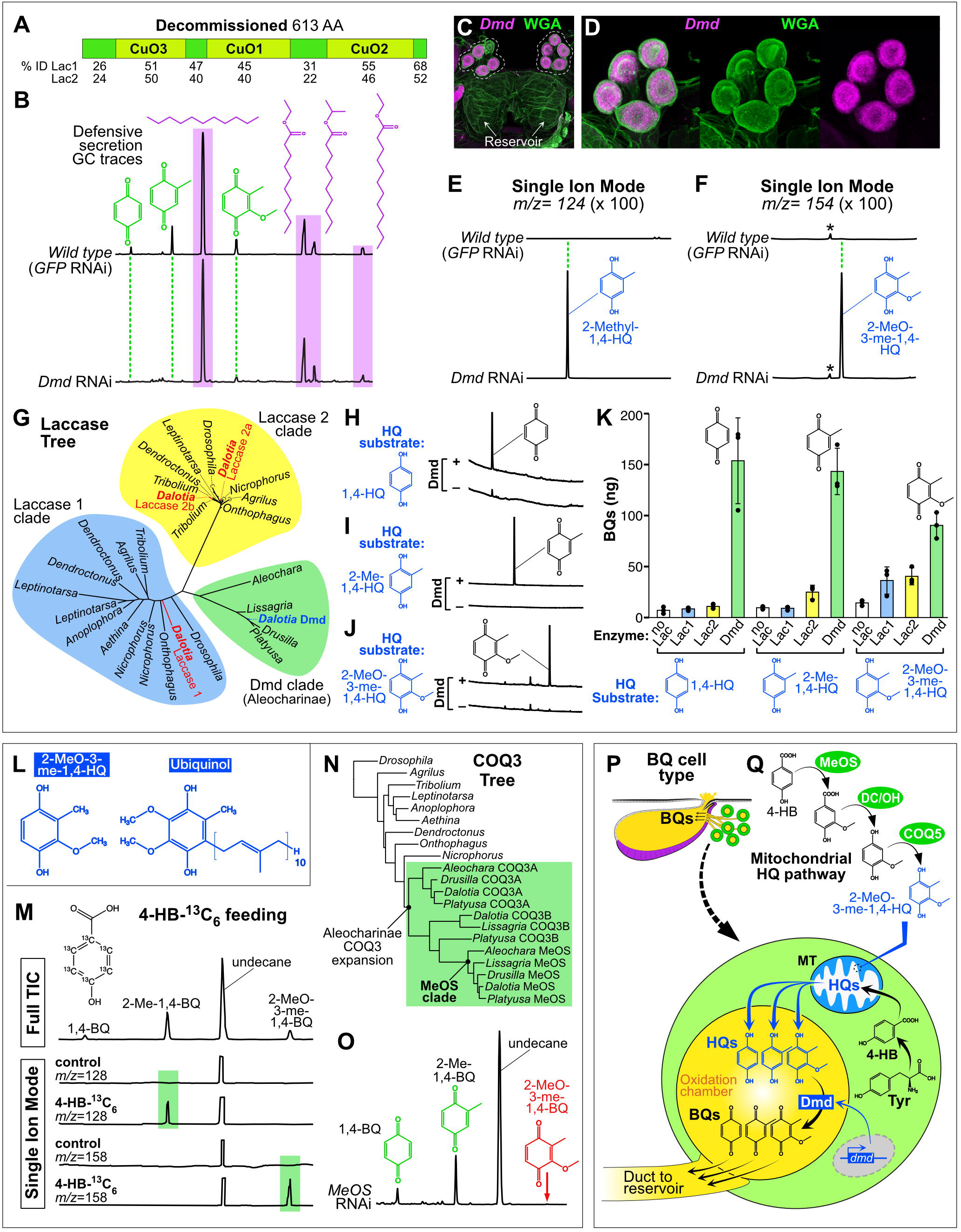
Functional evolution of the BQ cell type. **A:** Domain structure of Decommissioned with percentage sequence homology to domains of Laccase 1 and 2. **B:** GC traces of *Dalotia* tergal gland secretions. Top: wild type secretion. Bottom: Silencing *Dmd* causes specific loss of BQs. **C, D:** HCR (magenta) and Wheat Germ Agglutinin (green) labelling of the tergal gland reveals of *Dmd* expression in the bulbous secretory cell clusters (BQ cells). Panel **E, F:** Single ion GC traces of hydroquinone molecular ions 2-Me-1,4-HQ (*m/z*= 124) (**E**) and 2-MeO-3-Me-1,4-HQ (*m/z*= 154) (**F**) in wild type (top) and *Dmd*-RNAi animals (bottom). **G:** Unrooted tree of insect laccases showing aleocharine-specific Dmd clade, distinct from conserved Lac1 and 2 clades. **H–J:** Activity of purified Dmd *in vitro* as assayed by GC-MS when Dmd protein is provided with different HQ substrates. GC traces reveal efficient oxidation of HQs to BQs in the presence of Dmd (upper traces) but not in controls lacking Dmd (lower traces). HQ substrates are 1,4-HQ (**H**); 2-Me-1,4-HQ (**I**); 2-MeO-3-Me-1,4-HQ (**J**). **K:** Activity of purified Dmd *in vitro* relative to laccase 1 and laccase 2. Enzymes were provided different HQs as substrates and levels of BQs were measured using GC-MS. n=3 replicates per enzyme for the three HQ substrates. **L:** Molecular structures of 2-MeO-3-Me-1,4-HQ and Ubiquinol. **M:** Diagnostic ions for detection of 4-HB-^13^C6 incorporation into 2-Me-1,4-BQ and 2-MeO-3-Me-1,4-BQ. Top trace is the total ion current (TIC) wild type chromatogram; lower traces show extracted ion traces of diagnostic ions for BQs labelled with ^13^C6 (*m/z*= 128 for 2-Me-1,4-BQ and *m/z*=158 for 2-MeO-3-Me-1,4-BQ). The intensity of *m/z*= 128 and *m/z*= 158 ions was magnified 100x in control and 4-HB-^13^C6 fed beetles compared to the TIC chromatogram. **N:** Tree of COQ3 enzymes showing single copies in all outgroup beetle species and *Drosophila* but multiple duplicates in Aleocharinae (green boxed clade) including the unique paralogue Methoxyless (MeOS). **O:** GC trace of gland secretion following MeOS knockdown shows specific loss of 2-MeO-3-Me-1,4-BQ. **P:** Model of BQ cell function: Tyr-derived 4-HB is transported to the mitochondrion (MT) and modified to hydroquinones (HQ) via a ubiquinol-like pathway (**Q**). HQs are secreted into the BQ cell lumen for oxidation by Dmd. BQs travel to the reservoir via the duct. **Q**: Proposed pathway for HQ synthesis with 2-MeO-3-me-1,4-HQ as an example. DC/OH=unidentified decarboxylase/hydroxylase.

We confirmed *Dmd*’s expression in the bulbous secretory cells using HCR (Fig. 6 D, C), establishing them as the source of the BQs. We refer to these cells as the ‘BQ cell type’. Because Dmd is secreted, we hypothesized that it may be a terminal pathway enzyme that oxidizes secreted BQ precursors. Laccases are well known to oxidize hydroxyl groups, and studies in other aleocharine rove beetles have shown that, in addition to BQs, trace levels of corresponding hydroquinones (HQs) can occasionally be detected in the secretion, consistent with HQs being unoxidized precursors (Steidle and Dettner, 1993). In keeping with these findings, we sometimes recover trace levels of 2-methyl-1,4-HQ in *Dalotia*’s secretion; this HQ corresponds to the non-oxidized form of 2-methyl-1,4-BQ, which is the highest abundance BQ. Strikingly, we find that although silencing Dmd diminishes levels of 2-methyl-1,4-BQ, it leads to excess levels of 2-methyl-1,4-HQ (Fig. 6E). Further, a HQ precursor of a different BQ species, 2-methoxy-3-methyl-1,4-HQ, additionally appears (Fig. 6F). Accumulation of HQs following *Dmd* silencing provides strong *in vivo* evidence that this laccase oxidizes -OH groups of secreted HQs, converting them into BQs. We relate this function to the unique BQ cell anatomy, where the cell envelopes a lumen connected directly to the duct (Fig. 6P). We posit that the lumen is an oxidation chamber into which Dmd and HQs are secreted and combine. The duct channels the resulting cytotoxic BQs into the solvent reservoir (Fig. 6P).

We recovered *Dmd* orthologues in genomes of other aleocharines that possess a homologous tergal gland and synthesize BQs (Fig. 6G). Phylogenetic analysis reveals that Dmd does not branch from within either of the conserved insect laccase clades, Lac1 and Lac2 (Fig. 6G). As in other insects, *Dalotia Lac2* is expressed in cuticular cells (Fig. S19E) and silencing it abolishes tanning (Fig. S19B, C), while *Dmd* is not expressed in the cuticle (Fig. S19F) and is not involved in tanning (Fig S19B, D). Dmd thus defines a novel laccase clade specific to aleocharines that functions as a terminal oxidase in BQ synthesis. To corroborate this model, we synthesized Dmd protein and tested its ability to oxidize HQ precursors of all three BQs *in vitro*. Consistent with its inferred function, Dmd strongly catalyzed conversion of all three HQs to BQs (Fig. 6H-J, Fig. S19G). Further, we synthesized *Dalotia* Lac1 and Lac2 and found that although both enzymes exhibited some activity on at least one HQ substrate, neither enzyme was as efficient as Dmd (Fig. 6K). We conclude that Dmd is a catalytically specialized laccase that performs HQ oxidation in BQ cells.

### HQ synthesis via evolution of a ubiquinone-like pathway

What pathway produces the HQ precursors for Dmd? Our data indicate that BQs derive from Tyr (Fig. S18), but how the aromatic ring is hydroxylated to make HQs and decorated with methyl and methoxy groups, is unknown. One eukaryotic pathway exists that integrates these steps: ubiquinone biosynthesis. Here, Tyr is converted to 4-hydroxybenzoic acid (4-HB), which is modified in the mitochondrion to yield the HQ ubiquinol (Wang and Hekimi, 2019). *Dalotia*’s BQs resemble less modified versions of ubiquinol, at most incorporating a methyl and a methoxy group (Fig. 6L). To test whether HQs derive from a similar pathway, we fed *Dalotia* 4-HB-^13^C_6_ and assayed for ^13^C_6_ incorporation into the BQs. Although the absolute level of incorporation was not high, we observed significant ^13^C_6_ enrichment above baseline into the two most abundant BQs (Fig. 6M). The magnitude of ^13^C_6_ incorporation was lower than observed on feeding *Dalotia* Tyr-^13^C_6_ (Fig. S18), presumably because 4-HB is a catabolite, not a dietary precursor, limiting its access from the gut to the correct cellular location. This result nevertheless identifies 4-HB as a likely intermediate in the conversion of Tyr to HQs.

Key steps in 4-HB’s conversion to ubiquinol are its decarboxylation and hydroxylation to create a HQ by an unknown enzyme, and the addition of methyl, methoxy and prenyl groups by sequentially-acting CoQ enzymes (Wang and Hekimi, 2019) (Fig S20I). Two CoQ enzymes are of potential relevance: CoQ5, a methyltransferase that adds the methyl group, and CoQ3, an O-methyltransferase that creates the methoxy groups. We asked whether these enzymes add methyl or methoxy groups to *Dalotia*’s BQs. In most eukaryotes, including insects, CoQ enzymes are encoded by conserved single copy genes (Kawamukai, 2015). Due to their essential role in cellular respiration, studying them *in vivo* is challenging. For example, we observed ∼3-fold higher transcription of *CoQ5* in BQ cells (Fig. S20B), but knockdown with even low dsRNA levels led to complete lethality. Unusually, however, we found that *CoQ3* has duplicated in *Dalotia*, as well in the genomes of all other BQ-producing aleocharines surveyed (in some cases *CoQ3* has duplicated twice; Fig. 6N). Conspicuously, expression of one of *Dalotia*’s CoQ3 paralogues is strongly upregulated in BQ cells (Fig. S20D). Studies in humans, rats, yeast, and *E. coli*, have shown that CoQ3 adds two methoxy groups to ubiquinol using *S*-adenosylmethionine as the methyl donor (Jonassen and Clarke, 2000; Poon et al., 1999). An analogous reaction could conceivably yield the methoxy group of 2-methoxy-3-methyl-1,4-BQ. To test this idea, we fed *Dalotia* CD3-labelled methionine and observed direct incorporation of CD3 into 2-methoxy-3-methyl-1,4-BQ, confirming that the same reaction takes place (Fig. S20A). Remarkably, silencing the BQ cell-expressed CoQ3 duplicate led to complete loss of 2-methoxy-3-methyl-1,4-BQ in the secretion (Fig. 6O, Fig. S20E-G). Levels of *Dalotia*’s two other BQs were unaffected (Fig. 6O, Fig. S20G). We name this CoQ3 paralogue ‘*Methoxyless*’ (*MeOS*) and deduce that it performs an identical modification to a defensive BQ as canonical CoQ3 does to ubiquinone.

Canonical CoQ3 functions on the mitochondrial inner membrane, and we confirmed that MeOS is likewise targeted there (Fig. S20N). Furthermore, another enzyme, CoQ6, which performs essential priming hydroxylation prior to methoxylation of ubiquinol by CoQ3, is also upregulated in BQ cells (Fig. S20C), presumably permitting MeOS to methoxylate HQs (Fig. S20I). However, silencing *CoQ6* led to beetle lethality. Discovery of MeOS therefore provides a serendipitous genetic window into a mitochondrial route where 4-HB is modified by CoQ enzymes to make HQs, closely paralleling ubiquinol synthesis (Fig. S20I). Fully delineating the pathway *in vivo* is currently impossible owing to essential systemic functions of other CoQ enzymes. However, we postulate their co-option as a parsimonious model, and conjecture that *Dalotia*’s three BQs arise via differential processing: MeOS+CoQ5 activities produces 2-methoxy-3-methyl-1,4-HQ; CoQ5 activity alone leads to 2-methyl-1,4-HQ; whereas 1,4-BQ may arise from 4-HB decarboxylation and hydroxylation without further modification (Fig. 6Q, Fig. S20I).

### Coevolution of BQ and solvent cell types

Why did cell type evolution follow the routes that we have uncovered? The tergal gland secretion enhances beetle survival (Fig. 1C), so we reasoned that the BQ and solvent cells together underlie the gland’s adaptive value. To test this hypothesis, we performed a large-scale selection experiment, introducing *Dalotia* into multiplexed arenas with predatory ants and quantifying relative survival of beetles that were either wild type (GFP RNAi; 12 arenas, n=120 beetles), *MFASN*-silenced to inhibit solvent production (12 arenas, n=120 beetles), or *Dmd*-silenced to inhibit BQ synthesis (12 arenas, n=120 beetles). We allowed ants and beetles to interact for 48 h before assaying the survival difference between control and gene-silenced groups. In both *MFASN*- and *Dmd-*silenced treatments, we observed a comparable, significant reduction in survival (64% for GFP RNAi, 50% for *MFASN* RNAi, p <0.001, 45% for *Dmd* RNAi; p <0.001; Fig 7A). This result demonstrates each cell type’s adaptive value at the organismal level.

**Figure 7.**
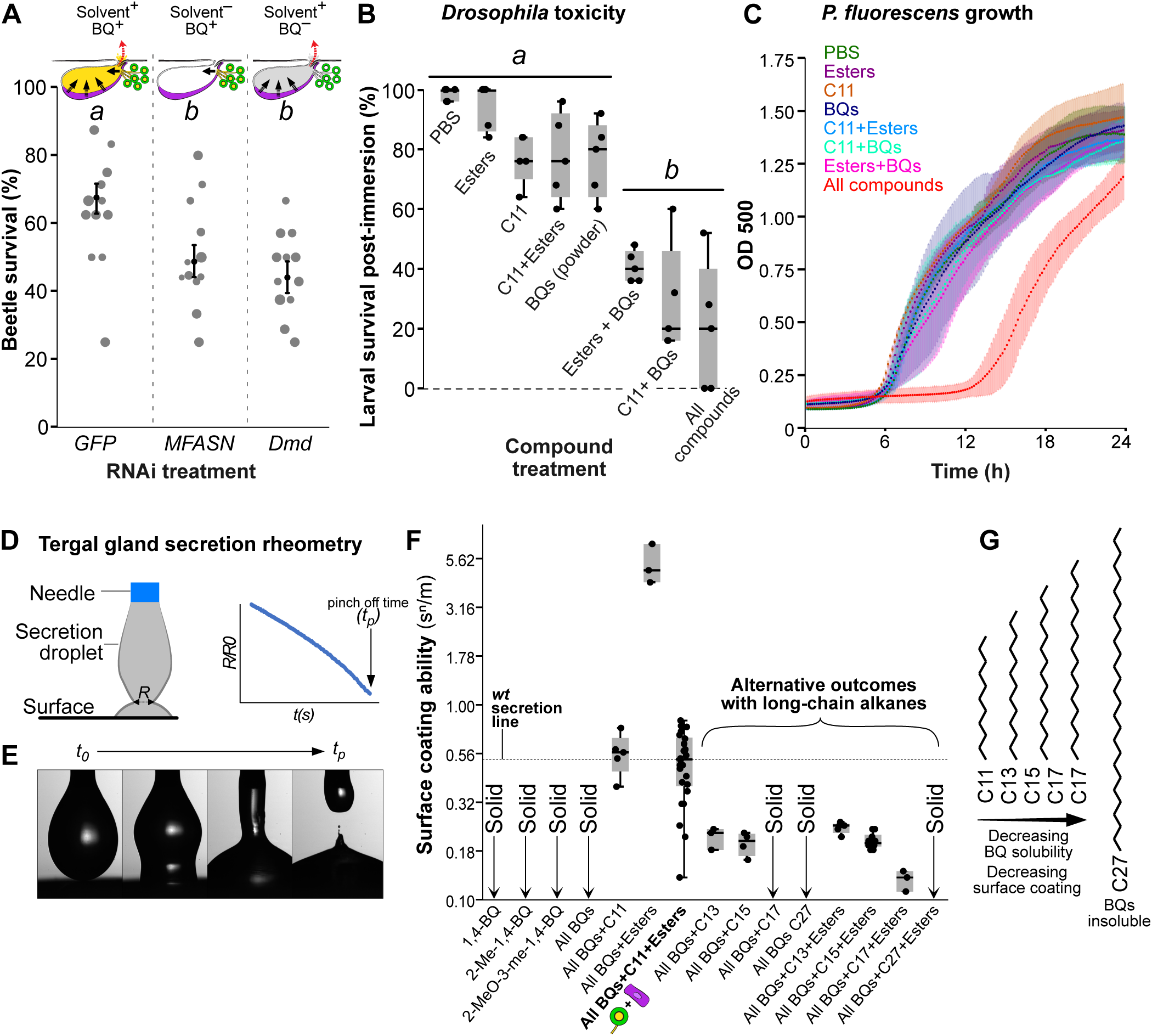
Synergism between solvent and BQ cells. **A:** *Dalotia* survival with predatory ants when BQ or solvent cells are compromised (GLM: Χ^2^ = 53.5, p< 0.001). Dot size corresponds to number of armed beetles per arena that were included in the statistical analysis. Black dots and bars are estimates and standard errors of the binomial model. **B:** Survival of *Drosophila* larvae 1h post-immersion in synthetic tergal gland secretion or its constituent (ANOVA: F_7,32_= 26.7, p< 0.001). Letters indicate significant differences. **C:** Growth curves of *Pseudomonas fluorescens* over 24h in presence of synthetic gland secretion or components (ANOVA_endpoint_: F_7,96_= 6.9, p< 0.001). **D**: Dripping-onto-Substrate (DoS) rheometer: synthetic secretion is dispensed via a nozzle onto an aluminum substrate. Droplet diameter (R) is measured over time until droplet pinch off. **E:** Video sequence of DoS measurement. **F:** Surface coating abilities (K/σ x 100 in s^n^m^-1^) of each mixture. **G:** With increasing chain length and boiling point of the alkane, the solubility of BQs and surface coating ability decrease. Detailed statistical analyses for all experiments are in supplementary Table S1.

Because the BQ and solvent cells secrete into a common reservoir, their products may combine to make a bioactive secretion. To test this, we explored how the compounds interact to shape the secretion’s physicochemical characteristics. Two parameters that capture salient properties of defensive secretions are the surface coating ability (SCA), a measure of wetting potential, and extensional viscosity (EV), a substance’s resistance to deformation when force is applied. The SCA and EV of arthropod secretions are species-specific and highly variable, presumably arising as a balance of factors including safe containment in gland reservoirs, controlled exudation, efficient spread across target tissues, as well as final irritancy. We employed a high-speed video-based rheometer to quantify SCA and EV of droplets of synthetic mixtures of different defensive compounds (Fig. 7D, E) (Dinic et al., 2016; Marshall et al., 2017; Rosello et al., 2019; Sousa et al., 2017). We found that SCAs of the BQ species alone or in combination are zero, whilst their EVs are virtually infinite (Fig. 7F, Fig. S21), because the three compounds remain in solid phase, even when mixed. However, adding undecane dissolves the BQs, creating a moderately viscous liquid with high surface activity (Fig 7F, Fig S21). Undecane thus acts as a solvent, unlocking the BQs. Equivalent SCA and EV are observed whether esters are present or not (Fig. 7F). Undecane is therefore the primary solvent, and the main determinant of the secretion’s SCA and EV.

We examined the biological consequences of this chemical synergism by measuring toxicity against other organisms. Using viability of *Drosophila* larvae as a readout, we made synthetic secretions and measured survival rates one hour after drastic, whole-body immersion. Dipping fly larvae for one second in a solution composed solely of undecane or the esters, or both compound classes combined, did not significantly reduce survival (Fig. 7B). Due to the BQs being solids, we could not immerse fly larvae in these compounds, but found that completely bathing larvae in BQ powder also did not significantly impact survival (Fig. 7B). However, when the BQs were combined with the undecane and esters at their natural ratios, mortality increased dramatically, reaching 80% (with 100% mortality in 2/5 replicates; Fig 7B). Comparable mortality was obtained if undecane was substituted for a higher ester fraction, or vice versa, demonstrating the critical effect of dissolving the BQs (Fig. 7B). These data indicate that the BQ and solvent cells are indeed engaged in biosynthetic synergism; only when their products are combined is a potent secretion with demonstrable adaptive value obtained (Fig. 1C, Fig. 7A).

We examined the antimicrobial effects of the secretion, noting that *Dalotia* self-applies the secretion topically to its own body. To do so, we assayed how synthetic combinations impact growth of the Gram-negative bacterium, *Pseudomonas fluorescens*—a common soil species and potential insect pathogen (Pineda et al., 2010; Scales et al., 2014). When added to *P. fluorescens* culture medium, no single compound class altered growth (Fig. 7C). Similarly, adding pairs of compound classes (undecane+esters, undecane+BQs, or esters+BQs) caused no observable effect (all pairwise tests p≥ 0.18) (Fig. 7C). Remarkably, however, adding all three compound classes together caused dramatic growth rate suppression (all pairwise tests p≤ 0.007) (Fig. 7C). Antimicrobial efficacy arising from combining these compounds is unexpected and striking, and to our knowledge has not been previously reported. The effect is again consistent with the adaptive utility of the tergal gland arising from cooperativity between the BQ and solvent cells.

*Dalotia*’s tergal gland secretion is thus a multi-compound cocktail with emergent properties possessed by no single component. While the BQs form the active defensive agent, the solvents provide a vehicle for the BQs while simultaneously conferring surface coating and tissue penetrating properties. We find two principal ways in which the solvent pathway may be specialized to meet these demands. First, the pathway favors high-level alkane and low-level esters synthesis. We noticed that silencing either *TG-CYP4G* or *TG-FAR* strongly increased levels of the C12 ester (‘ester shift’ in Fig. 2H, I). This is consistent with TG-CYP4G and TG-FAR normally titrating most C12-CoA towards undecane, permitting TG-αEst to make only trace ethyl dodecanoate (Fig. 1C; Fig. 2F). Accordingly, silencing *TG-CYP4G* or *TG-FAR* frees up C12-CoA for TG-αEst, resulting in elevated ethyl dodecanoate (Fig. 2H, I). The alkane branch is thus specialized for C12-CoA, while TG-αEst can use both C10- and C12-CoA. In this way, the solvent pathway produces large amounts of undecane as both the primary solvent and determinant of EV and SCA. This alkane bias strongly impacts the secretion: if undecane is replaced with the esters at their natural ratios, substantially higher EV and exceptionally high SCA are attained (Fig. 7F, Fig S21). Such a sticky, surface active secretion may be challenging to secrete and difficult to contain inside the reservoir. Low abundance esters are nevertheless critical for the secretion’s antimicrobial effect (Fig. 7C) and may also promote penetration into cuticles (Dettner, 1991).

A second aspect is the use of undecane as opposed to a longer chain alkane as the solvent. Undecane stands in contrast to the very-long-chain CHCs produced in oenocytes. During solvent pathway evolution, the importance of selective recruitment of a reductase (FAR) and decarbonylase (CYP4G) but not an elongase (ELO) is underscored by insolubility of BQs in even the shortest of *Dalotia*’s CHCs, heptacosane (C27) (Fig. 7F). However, the lack of ELO recruitment was not the only critical feature. Most insect FASNs produce C14-C18 fatty acids, rather than C12. We find that the corresponding alkanes (C13, C15 and C17), though only moderately longer than undecane, create secretions with much lower SCAs when mixed with the BQs, even in the presence of esters (Fig. 7F). In the case of heptadecane (C17), adding BQs without esters caused the secretion to freeze (Fig. 7F), and BQ crystals were still evident when esters were included. Hence, synthesis of C12-CoA by MFASN is central to creating an alkane that can both solubilize the BQs and create a topical secretion.

## Discussion

We have traced the evolution of new cellular functions comprising an organ novelty—the defensive tergal gland of rove beetles. First, we uncovered small molecule enzyme pathways that execute the biosynthetic functions of two taxon-restricted secretory cell types. Second, we presented evidence that these pathways—as well as an transcriptomic module expressed within one of the cell types—arose via repurposing components that functioned in more ancient cell types within the beetle. Third, we demonstrated the consequences of cell type evolution at the organ level, assigning adaptive value to the coordinated actions of the two cell types, including specialized features of the solvent pathway that render the secretion effective in chemical defense. Our findings underscore how the functional evolution of animal cell types can be a constrained process, employing pre-existing pathway motifs and expression programs and hence potentially convergence-prone. Conversely, our study demonstrates that the route followed by cell type evolution may also be a highly contingent process, conditional on other cell types within the organ and the fitness consequences of their collective output.

### Evolution of new biosynthetic functions

Using scRNAseq, we pinpointed specific cell types from which solvent pathway enzymes had been ancestrally sourced, enabling us to reconstruct its assembly via co-option or duplication and recruitment of oenocyte and fat body enzymes. We also determined that a transcriptome module that defines oenocytes and VFB, GEP 9, has been re-employed in solvent cells. Through these changes, an ancestral region of abdominal cuticle evolved into specialized secretory cells comprising the reservoir. That the capacity to manufacture defensive solvents is derived the beetle’s primary metabolism supports the view of genetic paths of least resistance to new animal chemistries—a notion stemming from pervasive convergence in the compounds that animals produce (Beran et al., 2019; Blum, 1981; Brückner and Parker, 2020). The BQ pathway likewise points to an underlying genomic blueprint; here, the HQ pathway, though not fully delineated at this time, owes at least part of its existence to the ubiquinone pathway from which the CoQ3 paralogue, MeOS, originated via duplication. We predict the connection will turn out to be still more extensive, with ubiquinone and HQ pathways sharing a subset of enzymes (Fig. S20I). Likewise, the role of the unique laccase, Dmd, in HQ oxidation parallels or may derive from the conserved function of Lac2 in cuticle tanning (Asano et al., 2019). We propose that repurposing pathway motifs from ancient metabolic cell types—or entire gene expression programs that facilitate production of certain compounds—may represent accessible routes to new biosynthetic functions in animal cell types. A consequence is that exploration of chemical space is constrained, leading to widespread chemical convergence across species.

### Cell type evolution via hybridization

Together with recruitment of solvent pathway enzymes, we propose that acquisition of GEP 9 was causal in transforming the intersegmental membrane between abdominal segments A6 and A7 into a biosynthetic epithelium. The membrane is naturally invaginated between segments in rove beetles (Fig. 5G); consequently, further growth created a reservoir into which the solvent cells could secrete their products (Fig. 5H). The antiquity of the oenocytes and fat body, which originated at the base of the Insecta, combined with the fact that GEP 9 is the sole program recovered in these cell types, indicates that the polarity of recruitment was from these ancient cell types into the solvent cells rather than vice versa. However, we do not rule out that some newer GEP 9 components may have evolved functions in the solvent cells and found secondary, pleiotropic utility in the oenocytes or fat body.

The solvent cell type sheds mechanistic light on the phenomenon of ‘cell type fusion’, where a novel cell type appears to take on features of two ancestral cell types (Arendt et al., 2016; Schlosser, 2018). Redeployment of a pre-existing transcriptional program in a new cellular context may be a common means for generating cellular- and organ-level novelties that lack direct homologs in other species or serially homologous body regions (typified by many animal glands). This mode of cell type evolution contrasts with duplication and divergence of sister cells within organs, or cells within repeating developmental fields such as body segments (Marioni and Arendt, 2016) or brain nuclei (Kebschull et al., 2020). Transcriptomic hybridization may be mechanistically facile if extensive expression programs conferring cell identity are under the control of one or a few transcription factors (i.e. ‘terminal selectors’; Hobert, 2016). The regulatory basis of GEP 9 expression in solvent cells is unknown, but this abdominal location is a region of overlapping expression of two Hox proteins, Abdominal A (AbdA) and Abdominal B (AbdB), both of which are needed for solvent cell development (Parker et al., 2017). Conceivably, a terminal selector controlling GEP 9 in oenocytes and VFB could have come under the control of AbdA and AbdB. Future work on the control of solvent and BQ cell identity may uncover how batteries of loci encoding the biosynthesis and secretory apparatus evolved coordinated expression in these cell types.

### Cell type evolution shaped by cooperative interactions at the organ level

Demonstrating adaptive change at the molecular level depends on connecting such changes to phenotypic outcomes that differentially impact fitness (Barrett and Hoekstra, 2011). Our data connect the evolution of new cellular functions to a cooperative interaction between cell types that dictates whole organ performance, and directly impacts animal survival. Hence, we propose that coevolution between the BQ and solvent cells has been driven, at least in part, by natural selection for organ-level properties. Our findings underscore how the phenomena of metazoan cell type and organ evolution are intrinsically coupled, with changes at the cellular level being comprehensible—and conferring adaptive value—only when their impact at the collective, multicellular organ level is considered (Kishi and Parker, 2021).

Inferring the steps leading to cooperativity is challenging due to the lack of known extant taxa with intermediate phenotypes. However, the recalcitrance of the solid BQs makes a ‘solvents first’ scenario likely, where alkane or ester compounds, or fatty acid-derived progenitors thereof, initially arose. Such compounds may have conferred modest chemical defense or acted as pheromones or lubricants for the flexible abdomen. They set the stage for subsequent evolution of BQs, the solvents unlocking their potency. Features of the solvent pathway such as the alkane bias, the use of medium chain compounds, and the presence of low-abundance esters, can be interpreted as further modifications to harness the BQs’ toxicity and manipulability (Fig. 7F, Fig. S21). Across aleocharine species, BQs are a constant feature of the secretion but the fatty acid derivatives vary extensively, including alkanes, alkenes, esters and aldehydes of differing chain lengths and ratios (Steidle and Dettner, 1993). These differences are expected to strongly influence the secretion’s physicochemical nature, emphasizing reciprocal molecular coevolution between the solvent and BQ cells.

Cooperative behavior is a feature of biological systems at all scales of organization. The tergal gland, comprising the simplest case of only two cell types, presents a model for explaining the evolution of cooperativity at the organ level. According to this model, the solvent cells created a niche for the evolution of the BQ cells, which enhanced the gland’s adaptive value. The dependence of the BQs on the product of the solvent cells, and the reciprocal dependence of the solvent cells on the BQs to maintain the gland’s higher adaptive value, meant that the two cell types became ‘locked in’ as a unit evolving within constraints set by performance at the organ level. This hypothesized route towards cooperativity contrasts with models for cooperativity within protein complexes. Here, dependencies between subunits can arise via entrenchment of binding interactions that were ancestrally selectively neutral (Finnigan et al., 2012; Hochberg et al., 2020; Lukeš et al., 2011). In the tergal gland, we posit that interdependence has likewise been enforced, but through the addition of a new cell type that is functionally contingent on a pre-existing cell type, which itself became obligately reliant on the second cell type to realize a relative selective advantage. Such a scenario could iterate through further cycles, informing how cooperation between diverse cell types may arise generally within organs.

## Acknowledgements

We thank Yuriko Kishi, Tom Naragon and Julian Wagner for help with *in situ* HCR, transcriptome assembly and bioinformatics, respectively; Fan Gao, Lior Pachter and the Bioinformatics Resource Center at Caltech; Sisi Chen, Jong H. Park, Matt Thomson and the Single Cell Profiling and Engineering Center (SPEC) in the Beckman Institute at Caltech; Melanie Spero for help with bacterial experiments and Chelsey M. VanDrisse for providing reagents. We are grateful to Marianne Bronner, Michael Dickinson, Lior Pachter and members of the Parker lab for feedback on this paper. AB is a Simons Fellow of the Life Sciences Research Foundation (LSRF). This work was supported by a Rita Allen Foundation Scholars Award, an Alfred P. Sloan Research Fellowship, a Shurl and Kay Curci Foundation grant, a Klingenstein-Simons Fellowship Award and a National Science Foundation CAREER award (NSF 2047472) to JP.

## Author contributions

AB and JP designed the study. AB performed experiments with help from JP (microdissections), JMB (*in vitro* protein studies) and MY (microinjections). RWL performed rheological measurements and processed raw data, AB analyzed the data with input from SAK. SAK performed GO-term analysis. JP supervised the project. AB and JP wrote the manuscript with input from SAK, JMB and RWL. All authors discussed and commented on the manuscript.

## Materials and methods

### Beetle husbandry

The Greenhouse Rove Beetle (*Dalotia coriaria*, Kraatz) strain used in this study was originally donated by Applied Bionomics (Canada), and inbred for nine generations. Beetles were kept as previously described (Parker et al., 2017) and fed with oat/poultry-rearing pellet-powder three times per week.

### Microbial suppression

Late third instar larvae were fed with a mixture of sterilized oat/poultry-rearing pellet-powder with amoxicillin, streptomycin and tetracycline antibiotics (2.5% w/w for each) or pure sterile oat/poultry-rearing pellet-powder as a control. Beetles were single-housed in 5 cm petri dishes with a thin layer of Plaster of Paris. Petri dishes were moisturized three times a week and fresh food was provided ad libitum. Ten days after adult eclosion, tergal gland contents were extracted in hexane and analyzed via GC-MS (see below). To assess the effect of antibiotic treatment on absolute numbers of bacteria associated with the beetles, bacterial 16S rRNA copy numbers determined by quantitative PCR (qPCR). Since many insects have bacteria on their cuticles, the beetles were surface washed in 5% (*v/v*) sodium dodecyl sulfate solution before bacterial quantification. For the control and antibiotic treatments, DNA was extracted from eight replicates of individual *Dalotia* using the Zymo Research Quick-DNA Miniprep Plus Kit according to manufacturer’s instructions. For qPCR, we used universal eubacterial 16S rRNA gene primers (Univ16SRT-F: 5’-ACTCCTACGGGAGGCAGCAGT-3’; Univ16SRT-R: 5’-TATTACCGCGGCTGCTGGC-3’) (Clifford et al., 2012). For quality assessment of DNA extracts and standardization of bacterial titers, qPCR with primers targeting host 28S rRNA was conducted simultaneously (D3A _F: 5’-GACCCGTCTTGAAACACGGA-3’; and D3B_R: 5’-TCGGAAGGAACCAGCTACTA-3’) (Litvaitis and Rohde, 1999). qPCR was performed on an Applied Biosystems 7300 Real Time PCR System in final reaction volumes of 25 µl, including the following components: 1 µl of DNA template, 2.5 µl of each primer (10 µM), 12.5 µl of autoclaved distilled H_2_O, and 6.5 µl of Luna® Universal qPCR Master Mix (New England BioLabs). PCR conditions were: 95°C for 5 minutes, 40 cycles at 95°C for 10 seconds, 70°C for 15 seconds, and 72°C for 10 seconds. A melting curve analysis was performed by increasing the temperature from 60 °C to 95 °C within 20 min. Standard curves were established for host 28S and bacterial 16S using PCR product as templates. A Qubit fluorometer (Thermo Fisher) was used to measure DNA concentrations to calibrate standard curves. The ratio between absolute copy numbers of bacterial 16S and host 28S (=bacterial/host copy ratio) was used as a standardized measure of bacterial abundance per beetle sample and the difference between the antibiotic-treatment and control group beetles was assessed with a Mann-Whitney-U-test.

### Biochemical tracer experiments and SIM mass spectrometry

Late third instar larvae were fed sterilized oat/poultry-rearing pellet-powder with amoxicillin, streptomycin and tetracycline antibiotics (2.5% w/w for each) plus 25% (*w/w*) stable isotope-labeled precursors: ^13^C_6_-D-glucose, ^13^C_6_-tyrosine, ^13^C_6_-phenylalanine (all > 99% enrichment, Sigma-Aldrich), D_3_-methione, D_23_-dodecanoid acid and ^13^C_6_-4-hydroxybenzoic acid (98-99% enrichment, Cambridge Isotope Laboratories, Inc.) as well as a control with sterile oat/poultry-rearing pellet-powder. Beetles were single-housed and glands of adults were extracted ten days after eclosion using hexane. Crude hexane extracts were analyzed with a GC-MS as described in detail below. Electron ionization mass spectra of characteristic fragment ions were monitored in single ion mode (SIM) and at 70 eV.

### Artificial disarming and survival biotest against ants

We developed a protocol to artificially disarm *Dalotia*, creating beetles that lack the tergal gland secretion. Adult beetles were collected from laboratory stock populations and placed on a CO_2_ fly pad; after the beetles were initial anesthetized the valve was closed and beetles could recover. Subsequently, we pulsed the beetles with low doses of CO_2_ which initiated abdomen flexing and visible release of chemicals from the tergal gland. We repeated this cycle of anesthesia, recovery, and low pulses of CO_2_ five times. To check success of the protocol, a subgroup of CO_2_-treated beetles, as well as control animals form the stock population beetles were individually extracted in 70 μl hexane for 10 min and their glandular compounds profiled with GC-MS (see below). Control group beetle glands contained 6.3 ± 5.6 μg (mean±SD) of secretion, while CO_2_-treated animals contained 0.5 ± 1 ng (Fig. S1C). CO_2_-treatment did not affect survival of the beetles and after 72 h all 25 beetles in both the treated and control groups were still alive. For the survival biotest, ten *Dalotia* beetles were paired with five *Liometopum occidentale* Emery ants (collected: Chaney Canyon, Altadena, CA; 34°13’04.1″N 118°09’06.4″W). Beetles and ants were placed in a 100×100×50mm plastic box with 10mm Plaster of Paris and two rolled pieces of Kimwipe to create a habitat-free foraging space (Vucic-Pestic et al., 2010). In total, we prepared 14 boxes each of control and disarmed beetles (=140 beetles and 70 ants per treatment). The experiment ran for 48 h and the the percentage of surviving beetles calculated for each box. Difference in survival between control and disarmed groups was analyzed using a Mann-Whitney-U-test.

### Double-stranded RNA preparation and RNAi knockdown in *Dalotia*

Double-stranded RNA was prepared from cDNA from pooled from all life-stages of the beetle. Regions of 450-650 bp for locus were amplified using primers with T7 linkers (5’-TAATACGACTCACTATAGGG-3’). Fragments were cloned into a pCR™4-TOPO™ Vector (TOPO™ TA Cloning™ Kit, ThermoFisher) using primers listed in Table 1.

The same primers were subsequently used to amplify template DNA from the TOPO vector for dsRNA synthesis, using the MEGAscript™ T7 Transcription Kit (ThermoFisher). After synthesis, dsRNA was cleaned using acid phenol/chloroform (50:50) and adjusted to a concentration of ∼6 mg/ml. For injection, dsRNA stock was then diluted 1:1 in DEPC-treated 1x PBS and blue food dye following a previously published protocol (Philip and Tomoyasu, 2012). Following the same protocol, dsRNA against green fluorescent protein (*GFP*) was prepared and injected as a control. Late third instar larvae were collected from laboratory stock populations and microinjection was performed according to Parker et al. (2018). After injection, the larvae were individually placed into 5 cm plastic Petri dishes with thick moistened filter paper. Larvae that died before pupation or did not pupate by the end of ten days were discarded. The filter paper was moistened three times per week. After adults had eclosed, beetles were additionally fed with frozen fruit flies and oat/poultry-rearing pellet-powder on the same schedule for ten days. Mature beetles were used for experiments chemical assays.

### Extraction of gland compounds and gas chromatography/mass spectrometry

To screen chemical phenotypes of dsRNA knockdown and dsGFP control beetles, single specimens were submersed (Dettner, 1984, 1993b; Steidle and Dettner, 1993) in 70 μl hexane (company, GC/MS analytical grade) containing 150 ng/μl n-octadecane (Sigma Aldrich) as an internal standard; after 10 min the solvent was separated from the insect, transferred into a new vial and frozen at −80°C for further analysis. A GCMS-QP2020 gas chromatography/mass-spectrometry system (Shimadzu, Kyōto, Japan) equipped with a ZB-5MS fused silica capillary column (30 m x 0.25 mm ID, df= 0.25 μm) from Phenomenex (Torrance, CA, USA) was used for chemical profiling. Crude hexane sample aliquots (2 μl) were injected by using an AOC-20i autosampler system (Shimadzu) into a split/splitless-injector which operated in splitless-mode at a temperature of 310°C. Helium was used as the carrier-gas with a constant flow rate of 2.13 ml/min. The chromatographic conditions were as follows: The column temperature at the start was 40°C with a 1-minute hold after which the temperature was initially increased 30°C/min to 250°C and further increased 50°C/min to a final temperature of 320°C and held for 5 minutes. Electron impact ionization spectra were recorded at 70 eV ion source voltage, with a scan rate of 0.2 scans/sec from m/z 40 to 450. The ion source of the mass spectrometer and the transfer line were kept at 230°C and 320°C, respectively.

Compounds were identified based on their *m/z* fragmentation patters and by comparison to authentic standards (1,4-benzoquinone, 2-methyl-1,4-benzoquinone, undecane, isopropyl decanoate, ethyl decanoate, ethyl dodecanoate; all SigmaAldrich; 2-methoxy-3-methyl-1,4-benzoquninone was synthesized as outlined below). Additionally, compound identity was confirmed by comparison of retention indices and MS data with published literature (Dettner, 1984; Dettner et al., 1985; Steidle and Dettner, 1993). The authentic standards were used to construct four-point calibrations curves for external standardization and quantification of BQs, esters and alkanes. Semi-quantification of bulk cuticular hydrocarbons (CHCs) was based on the internal C18 standard. We quantified the ion abundance and the absolute amounts of compounds in ng based on the internal or external standard (see above), as well as the relative composition of individual CHCs compared to ion abundance of the other CHCs. For quantifying gland compounds following RNAi silencing, we compared the target compound amounts between dsRNA knockdown and dsGFP control beetles with a Kruskal-Wallis test or Mann-Whitney-U-tests. Compositional data of CHCs was compared by non-metric multidimensional scaling (NMDS) ordination of a Bray-Curtis similarity matrix and analyzed using PERMANOVA (Brückner and Heethoff, 2017). GC-MS data related to RNAi can be found at CaltechData (https://doi.org/10.22002/D1.1917).

### In situ hybridization chain reaction (HCR)

For *in situ* HCR, DNA probe sets, DNA HCR amplifier, HCR hairpins as well as hybridization, wash and amplification buffers were purchased from Molecular Instruments (Beckman Institute at Caltech; https://www.molecularinstruments.com/) for each target transcript (see Table 2). For dissections, adult beetles were immersed in ice-cold DEPC-treated PBS and their abdomens were removed. Ventral segments were removed with dissection scissors. Dorsal abdominal segments were fixed in 4% PFA in DEPC-treated PBST for 25 minutes at room temperature and subsequently washed with DEPC-treated PBST (3×5 min). Fixation was followed by a 5 min proteinase K (10 µg/mL) treatment at room temperature, and samples were then rinsed in DEPC-treated PBST (3x). Samples were postfixed in 4% PBST-PFA for 25 minutes at room temperature and washed again with DEPC-treated PBST (3×5 min). The amplification and detection stages followed published protocols (Choi et al., 2018). Probes were either initiated with B1-Alexa546 or B4-Alexa488 amplifiers. After amplification and before the final wash steps, DAPI or Hoechst 33342 (1:2000) to mark nuclei, Alexa-488- or Alexa-647-Wheat Germ Agglutinin Conjugate (1:200) to label cell membranes and in selected samples BODIPY™ 493/503 (1:200) to stain for neutral lipids were added. Tissue samples were imaged as whole mounts of dorsal abdomens in ProLong Gold Antifade Mountant (ThermoFisher), using a Zeiss LSM 880 with Airyscan fast.

**Table 2.** In situ HCR targets and their amplifier systems.

| Target | Amplifier system | Molecular Instruments unique ID |
| --- | --- | --- |
| *O-CYP4G* | Alexa546-B1 | 3752/D845 |
| *Dmd* | Alexa546-B1 | 3752/D837 |
| *MFASN* | Alexa546-B1 | 3768/D873 |
| *TG-FAR* | Alexa546-B1 | 3752/D847 |
| *TG-CYP4G I* | Alexa546-B1 | 3834/E003 and 3752/D853 |
| *TG-CYP4G II* | Alexa488-B4 | 4096/E272 and 4096/E274 |
| *TG-α-EST* | Alexa546-B1 | 3834/E004 and 3752/D851 |
| *Tcas_MFASN* | Alexa546-B1 | 4096/E270 |

### Fly toxicity biotest

To test the toxicity of *Dalotia* tergal gland compounds, we followed an approach by Dettner (1984). First, artificial test solutions were prepared using the main gland constituents mimicking the natural ratios. An all-compound mixture (∼100 µl) was prepared by mixing 8 mg 1,4-benzoquinone, 28 mg 2-methyl-1,4-benzoquinone, 47 mg undecane, 7 mg isopropyl decanoate, 13 mg ethyl decanoate and 7 mg ethyl dodecanoate (all Sigma Aldrich). Subsequently, mixtures were prepared without certain compounds to create the following treatments: undecane+BQs, esters+BQs, all BQs, undecane+esters, undecane, esters; 1x PBS was used as control. For the survival tests a group of 25 *Drosophila melanogaster* third instar wandering larvae were immersed for 1 sec in 1 ml of artificial secretion or dipped into solid BQ powder and subsequently moved into a fresh culture tube. This protocol was repeated five times for each of the eight different treatments. Numbers of surviving fly larvae were counted after 1 h and the percentage calculated for each tube individually. The normally distributed and homoscedastic data were analyzed with a one-way ANOVA followed by a Tukey’s HDS pairwise posthoc test.

### Antimicrobial assay

To assay the antimicrobial properties of the *Dalotia* tergal gland secretion, we performed bacterial growth assays in a standard 96-well plate design. Artificial test solutions that mimicked natural ratios of compounds, as well and individual and pairs of compound classes were prepared as above (Fly toxicity biotest) and diluted in LB broth to a concentration of ∼6 µg/µl. 1x PBS was used as a control. Cultures of *Pseudomonas fluorescens* wild type (strain WCS365) (Jorth et al., 2019) were grown on LB agar plates overnight at 30°C. Subsequently, ∼5 µl of the bacterial colony was transferred into 500 µl and the OD_500_ was measured on a NanoDrop Spectrometer (Thermo Fisher). Based on the OD reading, the cell suspension was diluted to OD_500_= 0.1. For the growth assay, 15 µl of the cell suspension was inoculated with 134 µl LB and 1 µl of test secretion, and overlayed with 50 µl sterile mineral oil. Growth curves were recorded over 24h every 10 min on a Biotek Synergy 4 running in continuous shaking mode at 30°C. Each treatment and the negative PBS control were replicated 13 times across two plates. The OD_500_ values were plotted over time and the endpoint data after 24h (normally distributed and homoscedastic) was analyzed with a one-way ANOVA followed by a Tukey’s HDS pairwise posthoc test.

### Ant selection experiment

To test for an adaptive role of the BQ and solvent cells in chemical defense, we generated beetles with either BQ-free or solvent-free gland secretions by knocking down *Dmd* or *MFASN*, respectively, by injecting dsRNA (∼3 mg/ml) against these targets. As a control treatment, we injected dsRNA against *GFP* (∼3 mg/ml). Due to some mortality post-injection, we injected ∼400-500 larvae for each dsRNA target. The detailed RNAi protocol, primer sequences and beetle care were the same as those used for pathway characterization (see below). 10-day old adults were used to set up the following survival assays: like previously, ten beetles were paired with five ants (collected: Chaney Canyon, Altadena, CA; 34°13’04.1″N 118°09’06.4″W) in a 100×15 mm plastic Petri dish with a thin layer of Plaster of Paris and two rolled pieces of Kimwipe. In total, we prepared 12 dishes per treatment (=120 beetles and 60 ants per treatment), running the assay for 48 h before counted the number of surviving beetles. Because the amount of secretion per beetle cannot be determined before the experiment, we collected and extracted every beetle, regardless if dead or alive, in 30 µl of a mixture of 1:1 (v/v) hexane:chloroform for 15 min. We then ran a total of 360 beetle samples on the GC-MS using the same chromatographic conditions as outlined below. To determine if RNAi was successful in each individual beetle, we used single-ion detection of characteristic ions of 2-methyl-1,4-benzoquione (*m/z*= 122) and undecane (*m/z*= 156) to determine whether the glands of extracted beetles were benzoquinone-free (*Dmd* RNAi), solvent-free (*MFASN* RNAi) or not influenced (*GFP*). Only beetles with the correct gland composition were included in the final data set; beetles with completely empty glands at the end of the experiment were removed from the dataset because it is possible they possessed empty glands at the start of the experiment. For statistical analysis of survival differences among BQ-free, solvent-free and control beetles, we used a generalized linear model (GLM) with binomial error distribution and logit as link-function in R. We fitted ‘beetle survival’ as a binomial response variable of counted surviving beetles with a non-empty gland and total number of beetles (dead+alive) with a non-empty gland to account for the different sample sizes in each replicate. Finally, we used the R-implemented ‘Simultaneous Tests for General Linear Hypotheses’ with Tukey contrast for pairwise comparisons. The R script for this experiment can be found at CaltechData (https://doi.org/10.22002/D1.1918).

### Bulk RNAseq and transcriptome assembly

For bulk RNA sequencing (3 biological and 3 technical replicates), a total of 105 gland (Segment A7) and control (Segment A6) segments from male beetles were dissected and separately collected for total RNA extraction with Trizol™. Library preparation followed the Illumina TruSeq mRNA stranded kit protocol, including chemical shearing to obtain an average final library size of ∼300 bp. Illumina 100 bp paired-end sequencing was performed on a HiSeq2500 platform in which the twelve *Dalotia* gland libraries were multiplexed on the same lane. For genome-guided assembly of the transcriptome, a bam-file was created from the *Dalotia coriaria* reference genome assembly: (http://131.215.78.39/genomebrowser/cgi-in/hgGateway?genome=Dcor2&hubUrl=http://131.215.78.39/genomebrowser/evolution/parkergroup/Dalotia_coriaria/hub2.txt) using STAR (Dobin et al., 2013), while the approximately 240 million RNAseq reads were in silico normalized and subsequently used together with the bam-file to assemble transcripts using Trinity v2.8.4 (Grabherr et al., 2011). Finally, transcript sequences were clustered and concatenated to remove duplications using CD-HIT (Fu et al., 2012) with a sequence identity threshold of 0.95, yielding an assembly with a total length of 61.1 Mbp, an N50= 3097 bp and a BUSCO score (Simão et al., 2015) of C:97.6%[S:67.5%,D:30.4%],F:0.5%,M:1.6%. Annotations of transcripts were transferred from the respective gene models from the genome and additionally confirmed by blastx (Altschul et al., 1990) of all transcripts against the non-redundant NCBI protein database (Pruitt et al., 2007). Finally, the bulk-transcriptome was used to build a kallisto (Bray et al., 2016) index file for pseudoalignment of single-cell RNAseq reads and cell-type specific, ultra-low input RNAseq (SMART-Seq). The assembled transcriptome can be found at CaltechData: https://doi.org/10.22002/D1.1914.

### Cell-type specific transcriptome sequencing (SMART-seq)

For microdissection of specific gland cell types, male *Dalotia* beetles were injected with Invitrogen™ Wheat Germ Agglutinin, Alexa Fluor™ 488 Conjugate, to visualize the glandular structure *in vivo*, using a microinjector. Under fluorescence microscopy, we dissected the BQ cells in groups of 3-7 cells, the solvent cell tissue (∼1000 cells) and entire tergite 6 (control tissue) in ice-cold DEPC-treated PBS. We used the NEBNext® Single Cell/Low Input RNA Library Prep Kit for Illumina® together with NEBNext® Multiplex Oligos for Illumina® (New England Biolab) for library preparation, including cell dissociation, reverse transcription of poly(A) RNA, amplification full-length cDNA, fragmentation, ligation and final library amplification according to the manufacturer’s protocol. We performed cDNA amplification for 18-20 PCR cycles and final library amplification for 8 PCR cycles. In total, we constructed 31 libraries (ten D1, ten D2 and eleven control). The quality and concentration of the resulting libraries were assessed using the Qubit High Sensitivity dsDNA kit (Thermo Scientific) and Agilent Bioanalyzer High Sensitivity DNA assay. Libraries were sequenced on an Illumina HiSeq2500 platform (single-end with read lengths of 50 nt) with ∼20-25 million reads per library. Illumina sequencing reads were pseudoaligned to the bulk transcriptome and quantified (100 bootstrap samples) with kallisto 0.46.0 (Bray et al., 2016).

### Cell-type specific transcriptome analysis

For cell-type specific differential expression analysis, kallisto-quantified RNAseq data were processed with sleuth 0.30.0 (Pimentel et al., 2017) using Likelihood Ratio tests in R 3.6.1 (R_Core_Team, 2019). Normalized transcripts per million (tpm) from gland and control samples were extracted from the sleuth object, averaged and used to assign either up or down regulation for each transcript using a custom R script (CaltechData: https://doi.org/10.22002/D1.1918). Based on the differential expression data, we extracted transcripts that were down or upregulated in the BQ or solvent gland cells compared to control tissue.

### Preparation of single cell suspension for 10x scRNAseq

One hundred and fifty tergal gland segments (Segment 7) and control segments (Segment 6) of *Dalotia* males were dissected in EBSS (Earle’s Balanced Salt Solution; ThermoFisher) and transferred to into 150 µl ice-cold Schneider’s *Drosophila* medium and fetal bovine serum (SDM+FBS, 90:10 V/V; both ThermoFisher). After centrifugation at 250x g for 2 min, subsequent washing in 300 µl ice-cold EBSS and centrifugation for another 2 min at 250x g, the supernatant was replaced by 150 µl of pre-warmed (37°C) trypsin (Corning™ 0.25% Trypsin, 0.1% EDTA; Fisher Scientific). Cells were dissociated at 37ᵒC for 25 min and the reaction was reinforced by pipette mixing every 5 min. After 25 min, trypsin was inhibited with 100 µl soybean trypsin inhibitor (SigmaAldrich) and incubated for 5 min. Subsequently, 500 µl SMD+FBS was added and the cell suspension was passed through a 40 µm cell strainer (pluriStrainer, ImTec Diagnostics). Finally, cells were pelleted at 300x g for 8 min and resuspended in Dulbecco’s Phosphate-Buffered Saline (DPBS) 0.04% BSA (ThermoFisher) prior to 10x scRNAseq runs. For cell counting the LIVE/DEAD™ Viability/Cytotoxicity Kit (Invitrogen, ThermoFisher) was used. The above protocol was repeated twice for segment 6 and three times for segment 7, yielding five 10x runs.

### Single-cell RNA sequencing (scRNAseq) library preparation and sequencing

After manually determining the cell concentration using a hemocytometer, suspensions were further diluted to desired concentrations (∼1000 cells/µL) if necessary. The appropriate volume of reverse transcription mix was added to target 2,500–10,000 cells for loading into the 10x chip. The Chromium Single Cell 3’ Library & Gel Bead Kit v3, Chromium Single Cell 3’Chip kit v3 (PN-120236), and Chromium i7 Multiplex Kit were used for all downstream RT, cDNA amplification (12 PCR cycles), and library preparation as instructed by the manufacturer (Chromium Single Cell 3’Reagents Kits v3 User Guide). scRNAseq libraries were sequenced on an Illumina HiSeq4000 or NovaSeq6000 (paired-end with read lengths of 150 nt) and reads were aligned to the reference transcriptome using kallisto 0.46.0 (Bray et al., 2016). From a total of 1,669,423,289 reads, 86.3% mapped to the *Dalotia* transcriptome using kallisto | bustools (Melsted et al., 2019, 2021), giving an approximate sequencing depth of 100,000 reads per cell. The total number of transcripts detected across all cells was 26,259 (58.3%) out of 45,044 RNAs in the transcriptome. The mean number of genes per cell ranged between 1,227 for Segment 7 and 1,219 for Segment 6 (CaltechData: https://doi.org/10.22002/D1.1915 and https://doi.org/10.22002/D1.1918).

### Analysis of scRNAseq data

Kallisto alignment files were quantified with BUStools (v.0.39.3) (Melsted et al., 2021) and loaded into R using BUSpaRse (Moses and Pachter, 2021). We used knee plot methods as implemented in DropletUtils (Lun et al., 2019) to filter empty droplets before downstream analysis in Seurat v3.1 (Stuart et al., 2019). Briefly, we created Seurat objects for each library, filtering cells with low or overly large transcript counts, and merged all objects into one for quality assessment, and then split by library for normalization and scaling. From three 10x runs for Segment 7 and two 10x runs for Segment 6, we obtained 10,364 high-quality cells after stringent filtering (4,663 and 5,701 cells from Segments 6 and 7, respectively). Individual transcript counts in each cell were normalized by the total number of counts for that cell, multiplied by a scale factor (10,000), and natural log-transformed (NormalizeData function). Expression of each transcript was scaled to achieve a mean of zero and variance of 1 across cells (ScaleData function). Transcripts with high overall expression and variance across cells were identified (FindVariableFeatures function; highest standardized variance was selected by selection.method = ‘vst’). Subsequently, all five objects (libraries) were integrated, and shared sources of variation were identified by Seurat CCA alignment (Butler et al., 2018; Stuart et al., 2019). We selected the 3000 most variable transcripts across datasets (SelectIntegrationFeatures function) and used this set of transcripts to identify correspondences among the datasets, termed anchors (FindIntegrationAnchors function). Anchors were used to integrate the five datasets (IntegrateData function). The integrated Seurat object was again scaled and used as input for dimensionality reduction via Principal Component Analysis (PCA). Subsequently, we constructed a Shared Nearest Neighbor (SNN) graph for the dataset (FindNeighbors function) using the first 20 principal components of the PCA as well as an SNN pruning cutoff of 0.067 before performing cell clustering with the Louvain algorithm (FindClusters function; resolution of 0.25-1.0). We eventually used a resolution of 0.55, as this resulted in the separation of cell clusters that we had previously identified as different cell types by HCR (see CaltechData: https://doi.org/10.22002/D1.1918). Finally, we ran the Uniform Manifold Approximation and Projection (UMAP) dimensional reduction technique on the 20 principal components using default options, creating the abdominal cell type atlas (Becht et al., 2019; McInnes et al., 2018).

### Cell type annotation and statistical validation of cell classes

To annotate cell types, differentially expressed transcripts in cell clusters were identified in Seurat v3.1 (FindAllMarkers function) using both the Wilcoxon Rank Sum test as well as ROC analysis. Marker transcripts were computed using default settings for each cluster individually (FindMarkers function). A dendrogram was built in Past 4.02 (Hammer et al., 2001) using neighbor joining clustering (NJ) (see Musser et al., 2019 for a similar approach) on a Bray-Curtis similarity matrix (n=10,000 bootstraps) based on the normalized and scaled cluster average expression profiles of the 3000 most variable transcripts. We used the list of overall differentially expressed genes, the individual cluster markers, the results of the NJ clustering and the transcript annotation, to connect cell types to known cell type classes using published marker genes from fly expression data using http://flybase.org/, https://www.uniprot.org/ and https://bgee.org.

To test the robustness of NJ clustering, we employed random forest analysis to assign cells to their pre-defined cell type classes. Random forest is a machine learning classifier that assigns samples (here cells) to predefined groups (here cell type classes) in multiple iterations and estimates the correct classification (Breiman, 2001). For the analysis, ntree = 1,000 bootstrap replicates were drawn with mtry = 55 variables randomly selected at each node. A confusion matrix showing the number of correctly assigned cell communities to either cell type as well as proportional class errors and total out-of-basket (OOB) estimate of error rates were computed.

### Consensus non-Negative Matrix Factorization

We used the filtered cells-by-transcripts count matrix (10,634 cells by 26,259 transcripts) to infer gene expression programs (GEPs) of single cells using consensus Non-negative Matrix Factorization (cNMF) as implemented in Python (Kotliar et al., 2019). The method takes the count matrix (N cells x T transcripts) as input and produces both a gene expression program (GEP) x T matrix and a N x GEP matrix that specify the usage of each program for each cell in the data. The cNMF implementation runs multiple NMFs for each number of components (K) over a range of selected K, and combines the results of each replicate for different K to obtain more robust consensus estimates for the stability (silhouette score) and Frobenius error of each NMF model (Kotliar et al., 2019). We selected the most over-dispersed transcripts (T= 2000) using v-scoring (Klein et al., 2015) as the input for cNMF, which is essential for performing cNMF on biologically meaningful transcriptional variation (Kotliar et al., 2019). Expression of each over-dispersed transcript T was then scaled to unit variance and subjected to cNMF over a broad range of K (K= 16-36). The cNMF implementation was run with default parameters with the maximum number of iterations set to 250. Each iteration was run on the same normalized dataset over the range of Ks but with different randomly selected seeds. Subsequently, the factorizations for all iterations of each K were combined and the stability and error were computed. To choose the number of components K, we followed several approaches suggested by Kotliar et al. (2019); i) we plotted the stability and error as a function of K (Alexandrov et al., 2013), ii) performed a scree plot analysis of variance explained per principal component for the same data set to confirm the appropriate choice of K (Eckart–Young–Mirsky theorem; Eckart and Young, 1936) and iii) re-ran the same cNMF steps with a variable number of over-dispersed T to see whether stability and error converge on a similar K (CaltechData: https://doi.org/10.22002/D1.1915 and https://doi.org/10.22002/D1.1918; Fig. S16).

Ultimately, we selected K=20 and proceeded with the downstream cNMF workflow using default options, with density threshold r set to 0.05 in order to filter out replicates with low matching to a respective GEP (Fig. S16). We used the K=20 cNMF dataset to calculate the normalized GEP usage score for each cell, and then loaded the cell type or cell class assignment from our previous Seurat clustering/NJ+random forest analysis into Python to calculate GEP usage (in %) for each cell type/class. The resulting GEP usage matrix for each cell type was visualized as a heatmap. We also exported the normalized cells-to-GEPs (N x GEP) and GEPs-to-transcript (GEP x T) matrices (CaltechData: https://doi.org/10.22002/D1.1915), and loaded the N x GEP matrix into R as a matrix, integrating the GEP usage data as a new assay into the pre-existing Seurat object (CreateAssayObject function). This enabled us to plot % GEP usage across single cells, permitting visualization of GEP usage across the UMAP cell atlas.

Finally, we extracted the expression data for the GEP 9 to examine its composition. We assigned GO terms to each GEP 9 transcript and performed over-representation analysis (Yu et al., 2012) of GO terms to visualize the number of genes for each GO-term after GO-slim. GO terms for the respective gene models were assigned based on the gene id with highest homology from the SwissProt database (download February 2019) or NCBI nr database (downloaded February 2019). A custom database of GO terms was created with makeOrgPackage function in the R package AnnotationForge v1.26.0 (Carlson and Pages, 2019). Over-representation analysis of GO terms was tested using the enrichGO function in the R package clusterProfiler v3.12.0 (Yu et al., 2012) with a hypergeometric distribution and a Fisher’s Exact test. P-values were adjusted for multiple comparisons using false discovery rate correction (Benjamini and Hochberg, 1995). Enriched GO-terms were subsequently visualized using the REViGO tool available at http://revigo.irb.hr and the MonaGO tool available at https://monago.erc.monash.edu/.s17

### Phylogenetic analyses

To infer enzyme relationships, we first ran TransDecoder (Haas et al., 2013) using default settings to predict the longest ORFs in the bulk transcriptome and translate them into protein sequences. We then ran cd-hit (Fu et al., 2012) with default options and a sequence identity threshold of 0.9 on this protein set to cluster them and remove short-length sequence artifacts. The protein sequences of eight previously published beetle genomes and *Drosophila melanogaster* were then downloaded (Table 3) and we again ran cd-hit with default options and an identity threshold of 0.98 to remove isoforms. The ten protein sets were clustered into orthogroups with Orthofinder v2 (Emms and Kelly, 2015, 2019), using initial Diamond sequence alignment (Buchfink et al., 2015) and MCL clustering (Enright et al., 2002) to assign orthogroups. This was followed by multiple sequence alignment using MAFFT (Katoh and Standley, 2013) to align orthogroups and FastTree 2 (Price et al., 2010) to infer Maximum Likelihood protein trees for each orthogroup. When then used the orthogroup alignments containing our target sequences and concatenated them into one master fasta-file for each enzyme family and realigned the protein sequences using MAFFT. All alignment files and tree files have been deposited at CaltechData (https://doi.org/10.22002/D1.1916). Master alignments were each analyzed using IQ-TREE v1.6.12 (Nguyen et al., 2015) with 1,000 bootstrap replicates. Where appropriate, enzyme trees were rooted using *D. melanogaster*. Trees were visualized with FigTree v1.4.4 (Rambaut, 2012). For the analysis of laccase and CoQ3 enzymes, we queried sequences against a set of rove beetle genomes or transcriptomes and included homologous sequences in the phylogenetic analysis.

**Table 3.** Protein data sets used for OrthoFinder.

| Species | URL | Date |
| --- | --- | --- |
| *Drosophila melanogaster* | https://ftp.ncbi.nlm.nih.gov/genomes/all/GCF/000/001/215/GCF_000001215.4_Release_6_plus_ISO1_MT/GCF_000001215.4_Release_6_plus_ISO1_MT_protein.faa.gz | 01/21/2020 |
| *Agrilus planipennis* | https://ftp.ncbi.nlm.nih.gov/genomes/all/GCF/000/699/045/GCF_000699045.2_Apla_2.0/GCF_000699045.2_Apla_2.0_protein.faa.gz | 05/21/2020 |
| *Anoplophora glabripennis* | https://ftp.ncbi.nlm.nih.gov/genomes/all/GCF/000/390/285/GCF_000390285.2_Agla_2.0/GCF_000390285.2_Agla_2.0_protein.faa.gz | 05/21/2020 |
| *Leptinotarsa decemlineata* | https://ftp.ncbi.nlm.nih.gov/genomes/all/GCF/000/500/325/GCF_000500325.1_Ldec_2.0/GCF_000500325.1_Ldec_2.0_protein.faa.gz | 05/21/2020 |
| *Dendroctonus ponderosae* | https://ftp.ncbi.nlm.nih.gov/genomes/all/GCF/000/355/655/GCF_000355655.1_DendPond_male_1.0/GCF_000355655.1_DendPond_male_1.0_protein.faa.gz | 05/21/2020 |
| *Aethina tumida* | https://ftp.ncbi.nlm.nih.gov/genomes/all/GCF/001/937/115/GCF_001937115.1_Atum_1.0/GCF_001937115.1_Atum_1.0_protein.faa.gz | 05/21/2020 |
| *Tribolium castaneum* | https://ftp.ncbi.nlm.nih.gov/genomes/all/GCF/000/002/335/GCF_000002335.3_Tcas5.2/GCF_000002335.3_Tcas5.2_protein.faa.gz | 01/21/2020 |
| *Onthophagus taurus* | https://ftp.ncbi.nlm.nih.gov/genomes/all/GCF/000/648/695/GCF_000648695.1_Otau_2.0/GCF_000648695.1_Otau_2.0_protein.faa.gz | 05/21/2020 |
| *Nicrophorus vespilloides* | https://ftp.ncbi.nlm.nih.gov/genomes/all/GCF/001/412/225/GCF_001412225.1_Nicve_v1.0/GCF_001412225.1_Nicve_v1.0_protein.faa.gz | 01/21/2020 |

To characterize the cytochrome P450s of *Dalotia* belonging to the 4G subfamily, we downloaded a curated dataset of >350 sequences by Feyereisen (2020) and added the two *Dalotia* paralogues, performing a phylogenetic analysis with the same parameters as described in Feyereisen (2020). The tree was rooted with a *Panonychus citri* CYP450 sequence.

### Protein Expression and *in vitro* enzymatic assays for laccase

Synthetic cDNAs encoding the three laccases (Dmd, Lac1 and Lac2) were codon optimized for *E. coli* (CaltechData: https://doi.org/10.22002/D1.1917). Sequences were cloned into pTEV16 (VanDrisse and Escalante-Semerena, 2016) with a cleavable N-terminal 6xHis tag and TEV protease site. All proteins were expressed in BL21 competent *E. coli* cells (provided by C. VanDrisse, Caltech). Cells were grown in TB and protein expression was induced by 0.25 mM IPTG at OD600 ≈ 0.7 and harvested after 12 h at 20°C. Cells were lysed by freeze-thawing six times in lysis buffer (50 mM MES, 150 mM NaCl, 25 mM imidazole, DNase, and Thermo Scientific Halt Protease Inhibitor Cocktail). Proteins were passed over a column with Ni-NTA resin at room temperature, washed with 50 mM MES, 300 mM NaCl, 25 mM imidazole (pH= 7.5) and then eluted with 50 mM MES, 150 mM NaCl, 300 mM imidazole (pH= 7.5). Proteins were dialyzed with Thermo Scientific SnakeSkin for 4 h at room temperature into a final buffer of 10 mM MES and 100 mM CuSO_4_ and then concentrated to 5 mg/ml for experiments and storage at −80ᵒC.

Enzymatic activity of each protein was tested against a standard substrate, ABTS. The reaction mixture was prepared as 5 mM MES, 0.3 M CuSO_4_, and 2 mM of ABTS, with 2 mM of laccase. The UV spectrum was traced for 10 minutes. Subsequently, reactions were quenched with 5 M urea. To verify the RNAi results of the gland laccase Dmd, we set up the following reaction: 2mM of each HQ (1,4-hydroquinone, 2-methyl-1,4-hydroquinone and 2-methoxy-3-methy-1,4-hydroquinone) was separately added to vials (n=20 for each compound) containing the same reaction buffer previously used. To half of the samples we added 0.5 mM Dmd, while the other half served as control. Reactions were UV recorded for 1 minute and directly quenched with 0.05 M EDTA before being flash frozen and stored at −80ᵒC until further analysis. For the comparative analysis of Dmd, Laccase 1 and Laccase 2 we used the same buffers and substrates as previously. A total of three replicates and the same number of controls for each substrate-enzyme combination (i.e., 36 samples in total) were assayed.

To prepare samples for chemical profiling with either GC-FID or GC/MS for the Dmd vs control and comparative laccase assay, respectively, we used the following liquid-liquid extraction protocol. Frozen reaction mixtures were thawed and transferred into a clean glass vial with a sealed Teflon lid. For extraction, the aqueous phase was washed twice with 100 µl freshly Na_2_SO_4_-dried hexane. Organic fractions were combined, dried over Na_2_SO_4_ and carefully concentrated under a gentle stream of N_2_ to a volume of ≈50 µl. For chemical analyses sample aliquots (1 µl) were injected by an AOC-20i autosampler system from Shimadzu, into split/splitless-injector which operated in splitless-mode at a temperature of 310°C. Either a GCMS-QP2020 gas chromatography/mass-spectrometry system or a GC-2010PLUS with a flame-ionization detector (all Shimadzu, Kyōto, Japan) both equipped with a ZB-5MS fused silica capillary column (30 m x 0.25 mm ID, df= 0.25 μm) from Phenomenex (Torrance, CA, USA) were used. Helium was used as the carrier-gas with a constant flow rate of 2.13 ml/min. The chromatographic conditions were as follows: column temperature at the start was 40°C with a 1-minute hold after which the temperature was initially increased 50°C/min to 250°C and further increased 60°C/min to a final temperature of 300°C and held for 2 minutes. For the MS, electron impact ionization spectra were recorded at 70 eV ion source voltage, with a scan rate of 0.2 scans/sec from m/z 40 to 450. The ion source of the mass spectrometer and the transfer line were kept at 230°C and 320°C, respectively. For the FID, detector temperature was kept at 350°C. For quantification of the endpoint data [in ng] we used synthetic 1,4-BQ, 2-methyl-1,4-BQ (Sigma-Aldrich) and freshly synthesized 2-methoxy-3-methyl-1,4-BQ (see below).

### Rheological measurements

To measure the surface-coating ability and extensional viscosity of different artificial mixtures of synthetic secretions, we used dripping-onto-substrate (DoS) rheology. In contrast to conventional rheology which requires sample sizes on the order of milliliters, a DoS measurement can be accomplished with microliters. The instrument comprises a controlled dispensing system, a nozzle and a high-speed camera (Dinic et al., 2016; Marshall et al., 2017; Rosello et al., 2019; Sousa et al., 2017). Mixtures were prepared according to Table 4 using 1,4-BQ, 2-methyl-1,4-BQ, undecane, isopropyl decanoate, ethyl decanoate and ethyl dodecanoate (all Sigma Aldrich). 2-Methoxy-3-methyl-1,4-BQ was synthesized as follows: 2 mmol 2-Methoxy-3-methyl-1,4-hydroquinone (Sigma Aldrich) and 2 mmol NaIO_4_ were mixed in a 50 ml round-bottom flask and 10 ml CH_2_Cl_2_ and 5 ml of water were then added. The biphasic mixture was stirred at room temperature for 10 min and the layers were subsequently separated and then the aqueous layer was washed with 10 ml CH_2_Cl_2_. All organic fractions were combined, dried over Na_2_SO_4_ and concentrated *in vacuo*.

The all-compound mixture mimicking the beetles’ secretion was prepared by mixing 8 mg 1,4-benzoquinone, 18 mg 2-methyl-1,4-benzoquinone, 10 mg 2-methoxy-3-methyl-1,4-benzoquinone, 47 mg undecane, 7 mg isopropyl decanoate, 13 mg ethyl decanoate and 7 mg ethyl dodecanoate. For some experimental mixtures, undecane was replaced by tridecane, pentadecane, heptadecane or heptacosane (all Sigma Aldrich). The ratios of compounds in all mixtures always represented the average ratios of the respective compounds in *Dalotia*’s gland secretion (Fig. 1D). Mixtures were dispensed at a rate of 0.02 ml/min through the nozzle with an inner diameter of 0.413 mm and outer diameter of 0.718 mm onto an aluminum substrate a distance of 2.5 mm below the nozzle. The fluid transitions from the nozzle to substrate via an unstable liquid bridge. The behavior of this liquid bridge was captured by the camera at 20,000 to 40,000 frames per second and analyzed to produce rheological data on the fluid (CaltechData: https://doi.org/10.22002/D1.1905 and https://doi.org/10.22002/D1.1918). Each measurement was repeated at least 3 times and the videos were processed using Python 3. Briefly, the videos were converted to a series of binary images and the minimum diameter at each frame was recorded. The code and parameters for each video are available in the supplement (CaltechData: https://doi.org/10.22002/D1.1918). This yielded a dataset of diameter and time which was fit to an equation describing power-law fluids:

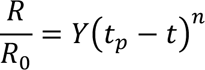

Here, *R_0_* is 0.359 mm, *n* is the flow behavior index, and *t_p_* is the pinch-off time, the moment the liquid bridge separates from the substrate. This equation was solved to estimate the elongational-flow consistency index Y (1/s^n^) and the flow behavior index n. Y describes what happens if a flow in imposed on fluid, while n describes how Y changes as the flow rate changes. Y decreases as the ratio of surface tension to viscosity decreases. When n= 1, mixtures behave as Newtonian fluids (no change with flow rate) and when n≠ 1 they behave as non-Newtonian fluid or power law fluids. For n < 1, fluid flows more easily (shear thinning) as the flow rate increases. The closer n is to 1, the weaker is the dependence on flow rate. For n > 1, the fluid more strongly resists flowing (shear thickening). Fluids with both low Y and low n possess higher extensional viscosity (EV). Furthermore, previous work has shown Y = ϕ(n)σ/K (Doshi et al., 2003). This is the product of the shape factor, ϕ(n), and surface tension divided by the power-law flow consistency index (Blair et al., 1939). The data fit this power-law function from the last frame with R/R_0_ = 1 to t = t_p_, with 0 to 5 datapoints (a maximum of ≈125 µs) excluded near t_p_ because during this period the fluid deviates from the power-law behavior. We solved the previous equation to get K/σ which is the ratio of the flow consistency index and surface tension. We interpret this ratio as a metric of how a fluid flows over a surface at a given surface tension. K/σ can therefore be used as a proxy to describe the ability of a mixture to coat a surface, like the cuticle of other insects. We refer to this metric as the surface coating ability (SCA).

### Data and code availability

#### Data availability

Raw sequence reads related to this manuscript have been deposited on NCBI under the BioProject ‘RNAseq (10x and SMARTseq) of the tergal gland of *Dalotia coriaria*’ (Accession: PRJNA707010; ID: 707010). All other data was uploaded to CaltechData: https://doi.org/10.22002/D1.1915 (processed scRNAseq 10x data), https://doi.org/10.22002/D1.1900 (processed SMARTseq data), https://doi.org/10.22002/D1.1905 (raw rheology video data), https://doi.org/10.22002/D1.1914 (transcriptome data), https://doi.org/10.22002/D1.1917 (RNAi experiments, survival assays, in vitro enzyme data), and https://doi.org/10.22002/D1.1916 (alignment and tree fasta files).

#### Data analysis and visualization software

Analysis and visualization of transcriptomic data were performed using Python 3.0 (Van Rossum, 2000) and R v.3.6.0 (Team and others, 2013) assisted by JupyterLab (Kluyver et al., 2016). The following R packages were used: AnnotationForge (Carlson and Pages, 2019), AnnotationHub (Morgan et al., 2019), biomart (Durinck et al., 2009) boot (Canty and Ripley, 2017), BUSpaRse (Moses and Pachter, 2021) car (Fox et al., 2012), clusterProfiler (Yu et al., 2012), cowplot (Wilke, 2017), data.table (Dowle et al., 2019), dbplyr (Wickham and Ruiz, 2019), DropletUtils (Lun et al., 2019) effects (Fox, 2003), ggplot2 (Wickham, 2011), lattice (Sarkar, 2008), lme4 (Bates et al., 2007), MASS (Ripley et al., 2013) Matrix (Bates and Maechler, 2010), nlme (Pinheiro et al., 2007), randomForest (Liaw and Wiener, 2002), Seurat v3 (Stuart et al., 2019) sleuth (Pimentel et al., 2017), tidyverse (Wickham et al., 2019), vcd (Meyer et al., 2020).

#### Code availability

Detailed code for scRNAseq analyses with Seurat and cNMF; video analyses of rheology data; custom R scripts for SMARTseq analyses via sleuth, GOterm assignments and survival data analysis can be found on CaltechData (https://doi.org/10.22002/D1.1918). All other statistical comparisons using ANOVAs, Kruskal-Wallis tests, U-tests and simple ordinations were done in Past 3.04 (Hammer et al., 2001).

## Supplementary Figure Legends

**Figure S1.**
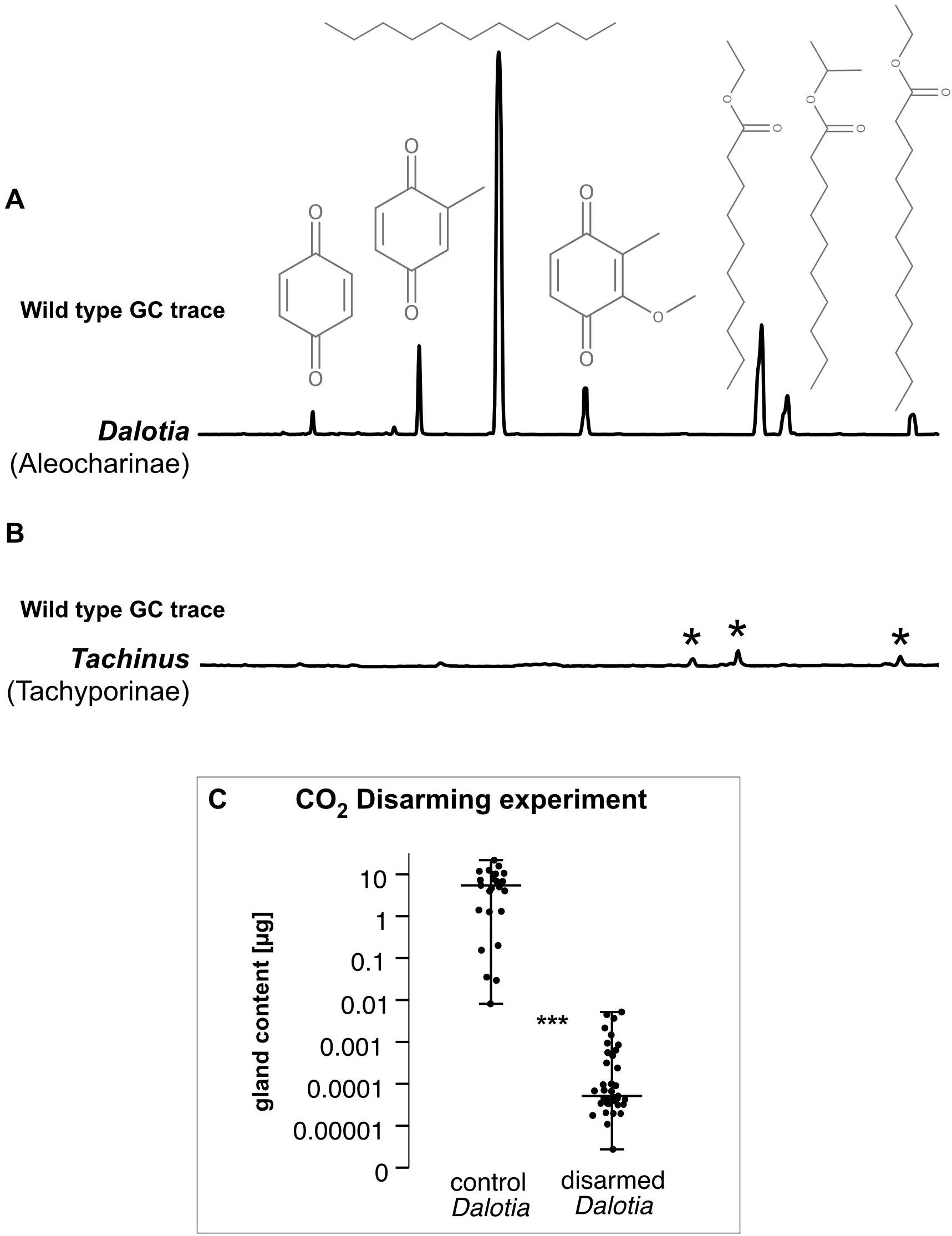
Comparison of the *Dalotia* tergal gland gas chromatographic (GC) trace to a gland-less outgroup rove beetle and CO_2_ disarming. A: Representative GC profile of the *Dalotia coriaria* gland content extracted in hexane. In order of retention: 1,4-benzoquinone (1,4-BQ), 2-methyl-1,4-BQ, undecane, 2-methoxy-3-methyl-1,4-BQ, ethyl decanoate, isopropyl decanoate, ethyl dodecanoate. B: Representative GC profile of *Tachinus* (Tachyporinae) extracted and measured using the same protocol. No gland compounds were detected. Asterisks denote in order of retention: C16:1-fatty acid (FA), C16-FA, C18-FA. C: Repeated treatment with small doses of CO_2_ leads to depletion of the tergal gland content in *Dalotia* effectively disarming the beetles. Significance was assessed using a Mann-Whitney-U test (***= p< 0.001).

**Figure S2.**
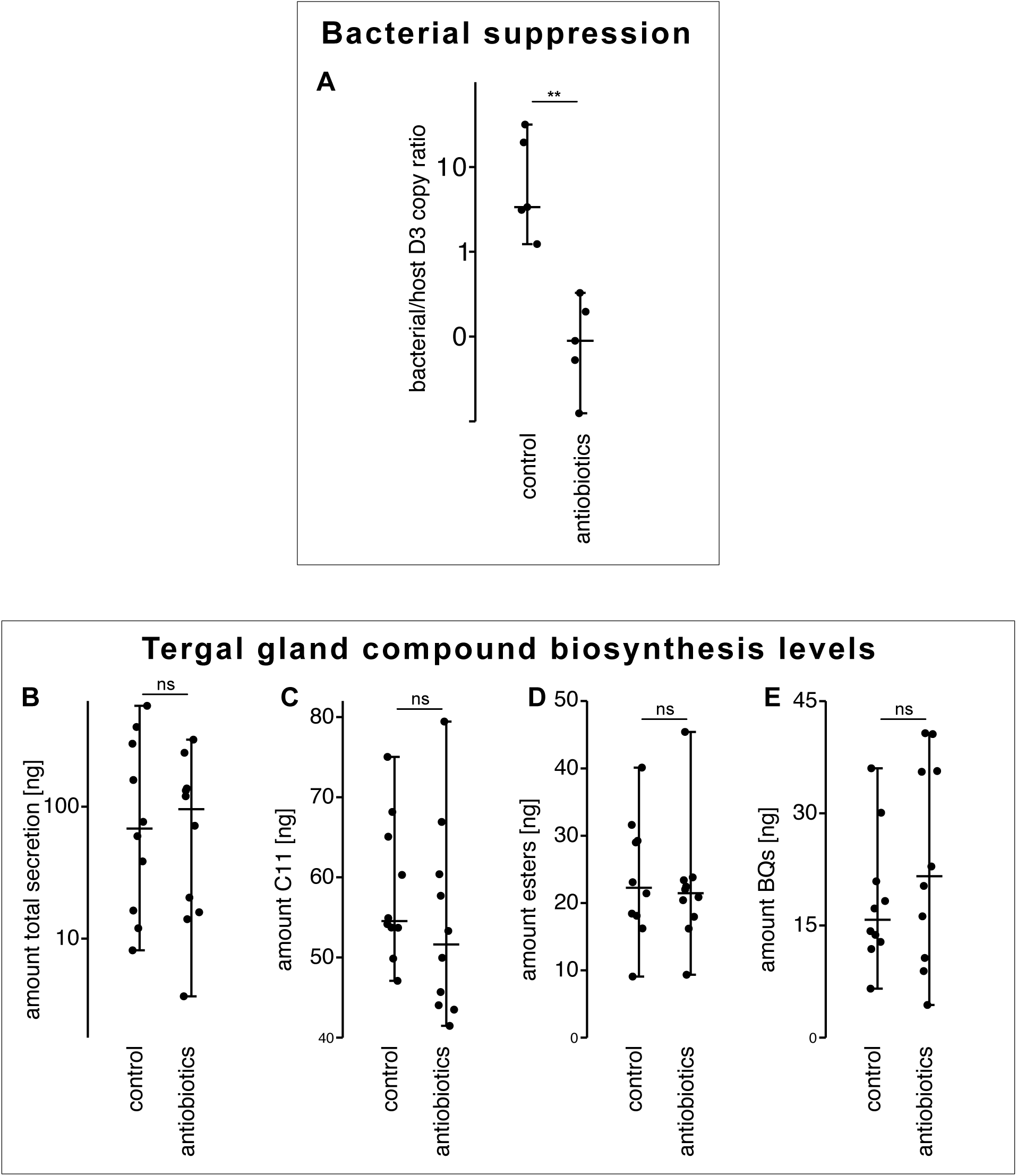
Biosynthesis of tergal gland compounds does not rely of symbiotic bacteria. **A:** Antibiotic treatment eliminates bacterial load in *Dalotia* as determined by qPCR quantification of the ratio of bacterial 16s rRNA to host 28s rRNA levels in antibiotic-treated and control beetles. **B-E:** Quantities of defensive compounds are unaffected by bacterial suppression: total secretion (**B**); undecane (**C**); the ester fraction (**D**); and the benzoquinones (**E**). Significance was assessed using Mann-Whitney-U tests (**= p≤ 0.01; ns= p≥ 0.05).

**Figure S3.**
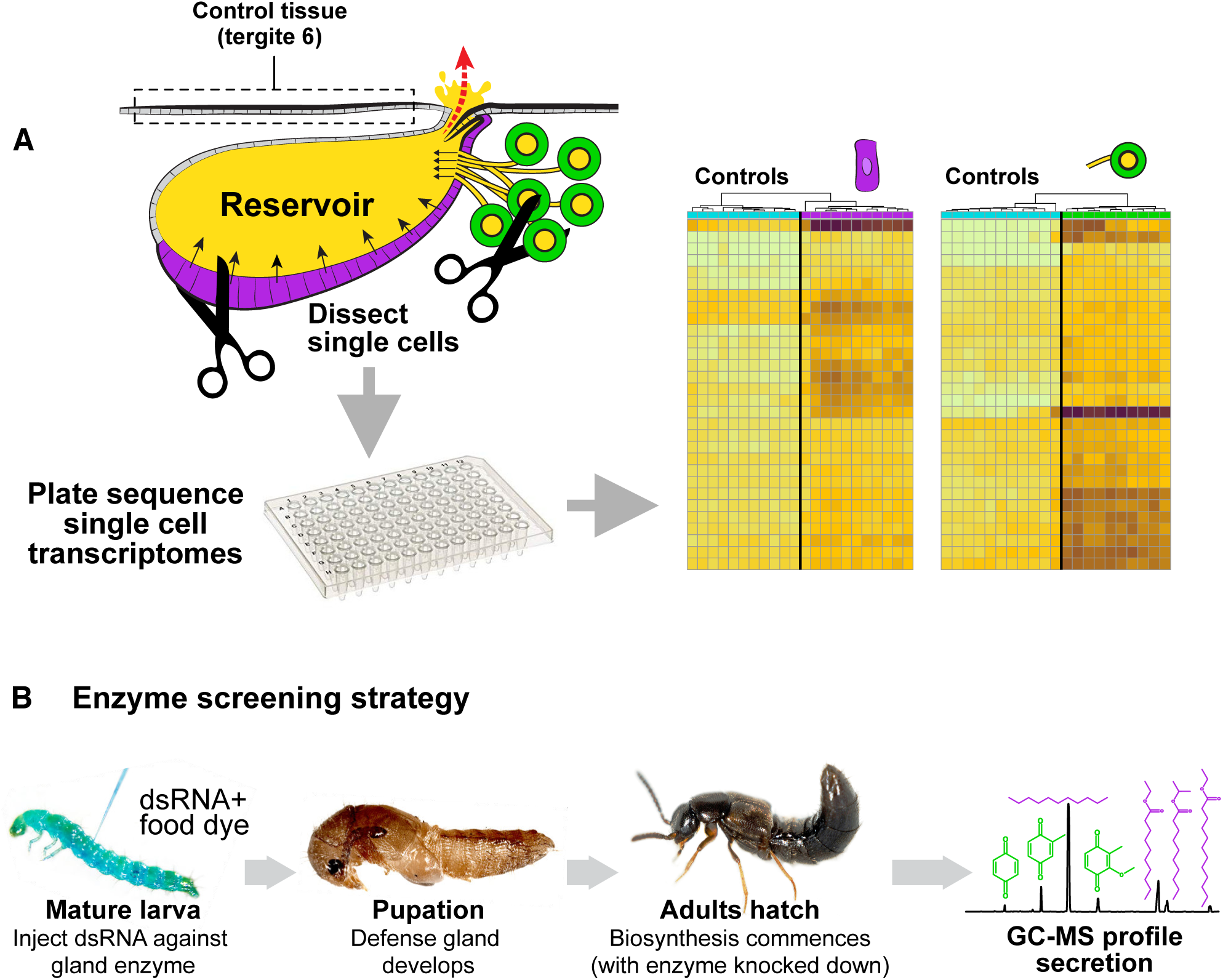
Preparation of cell-type specific transcriptomes and functional screening. **A:** To prepare cell-type specific transcriptomes of the two tergal gland cell types, cells were micro-dissected by hand and segment 6 tergite tissue was used as a control. Cells were lysed, and libraries prepared and sequenced to perform differential expression analysis. **B:** Schematic workflow of the enzyme screening strategy. By injecting mature larvae with double stranded RNA (dsRNA) targeting specific mRNAs, transcription is silenced in gland cells with temporal control, during pupal and early adult life when defensive compound biosynthesis occurs. The chemical composition of the tergal gland secretion can then be quantified in the mature adult via gas chromatography-mass spectrometry (GC-MS).

**Figure S4.**
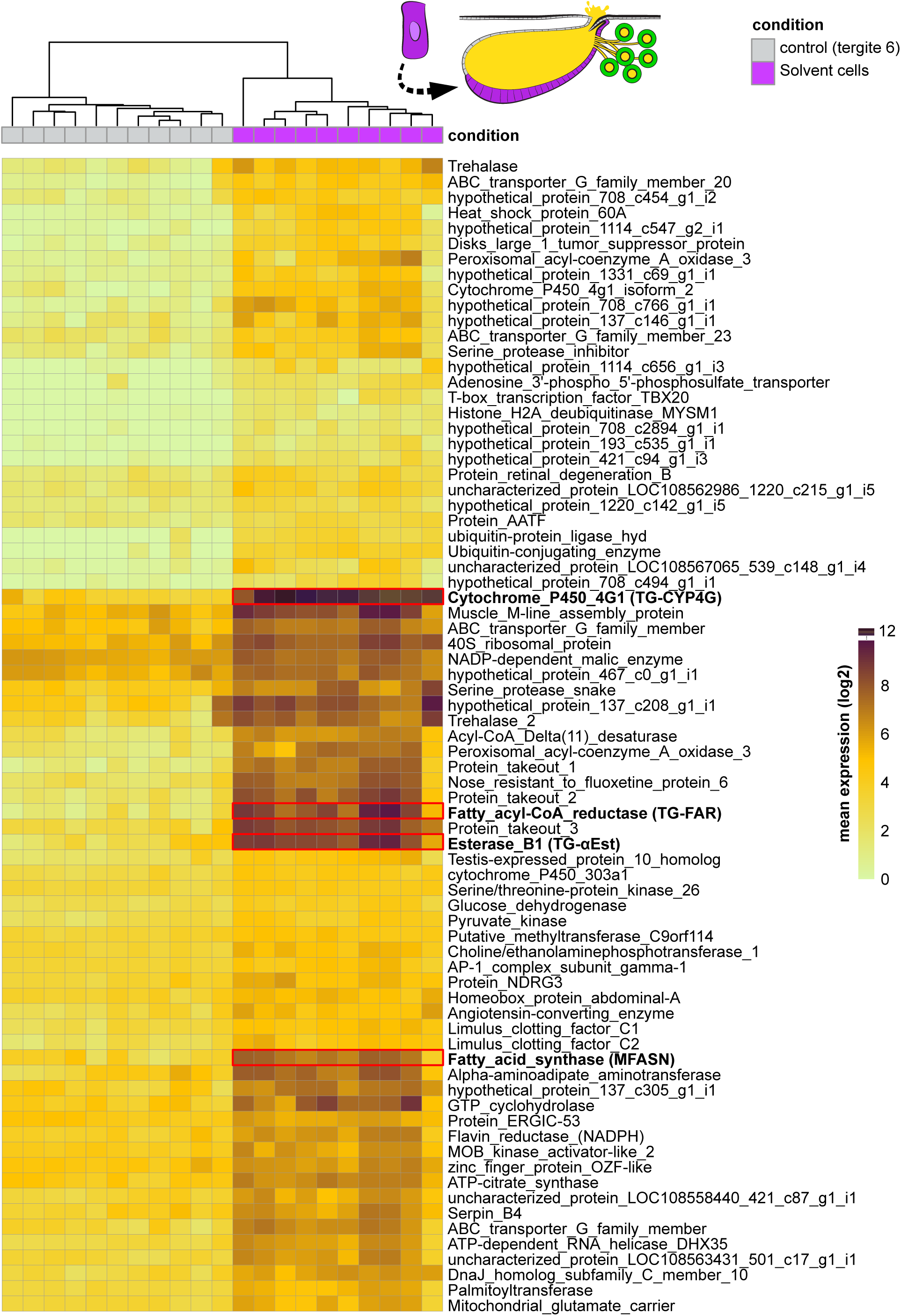
Differential expression analysis of solvent cells vs control cells. Heatmap showing the 75 most upregulated transcripts in the solvent cells (as assessed by q-value using sleuth; see https://doi.org/10.22002/D1.1918). Expression in all individual sequencing libraries is shown. Darker squares denote a stronger mean expression value. A full list of transcripts can be found at CaltechData (https://doi.org/10.22002/D1.1900).

**Figure S5.**
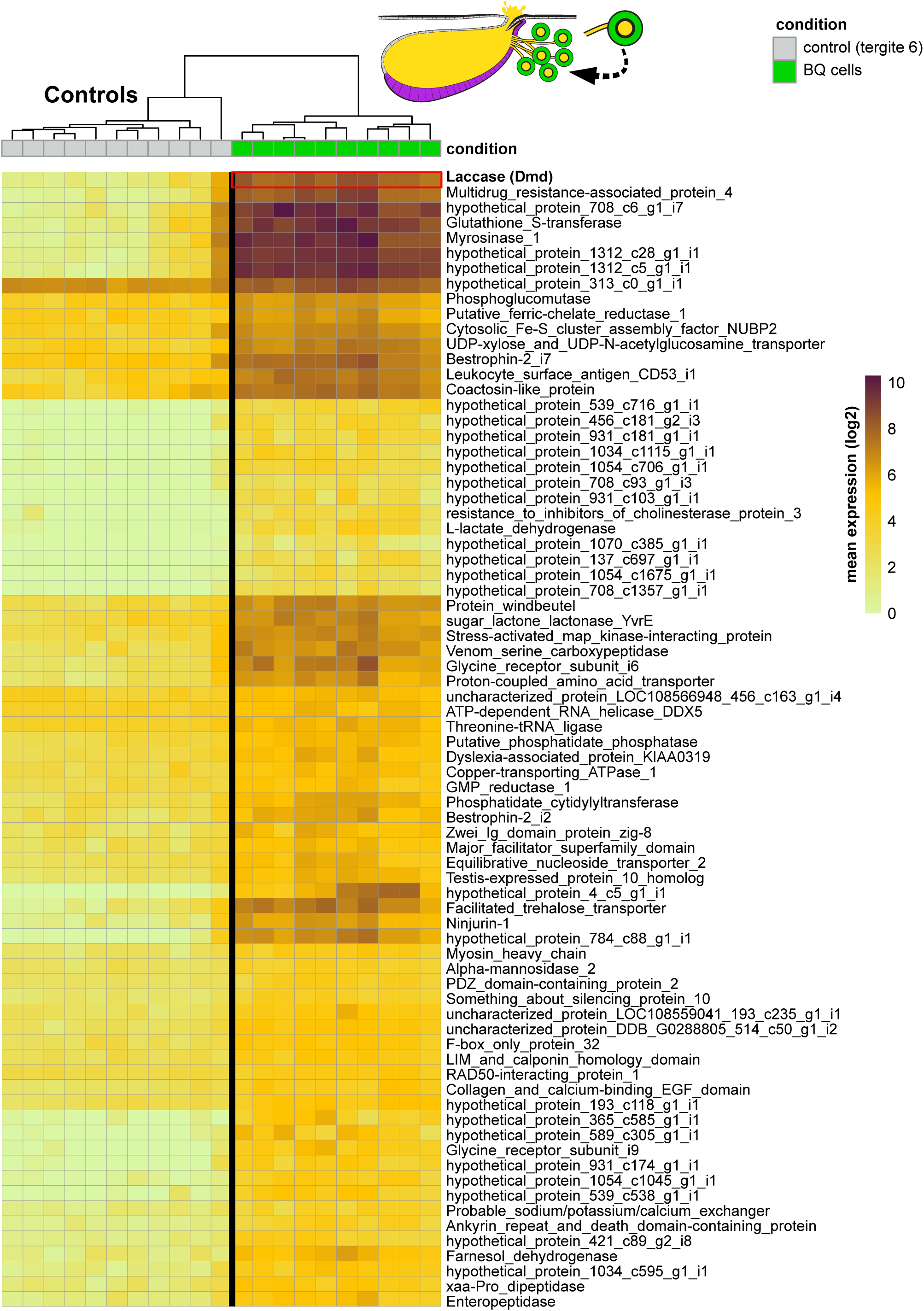
Differential expression analysis of BQ cells vs control cells. Heatmap showing the 75 most upregulated transcripts in the benzoquinone cells (as assessed by q-value using sleuth; see https://doi.org/10.22002/D1.1918). Expression in all individual sequencing libraries is shown. Darker squares denote a stronger mean expression value. A full list of transcripts can be found at CaltechData (https://doi.org/10.22002/D1.1900).

**Figure S6.**
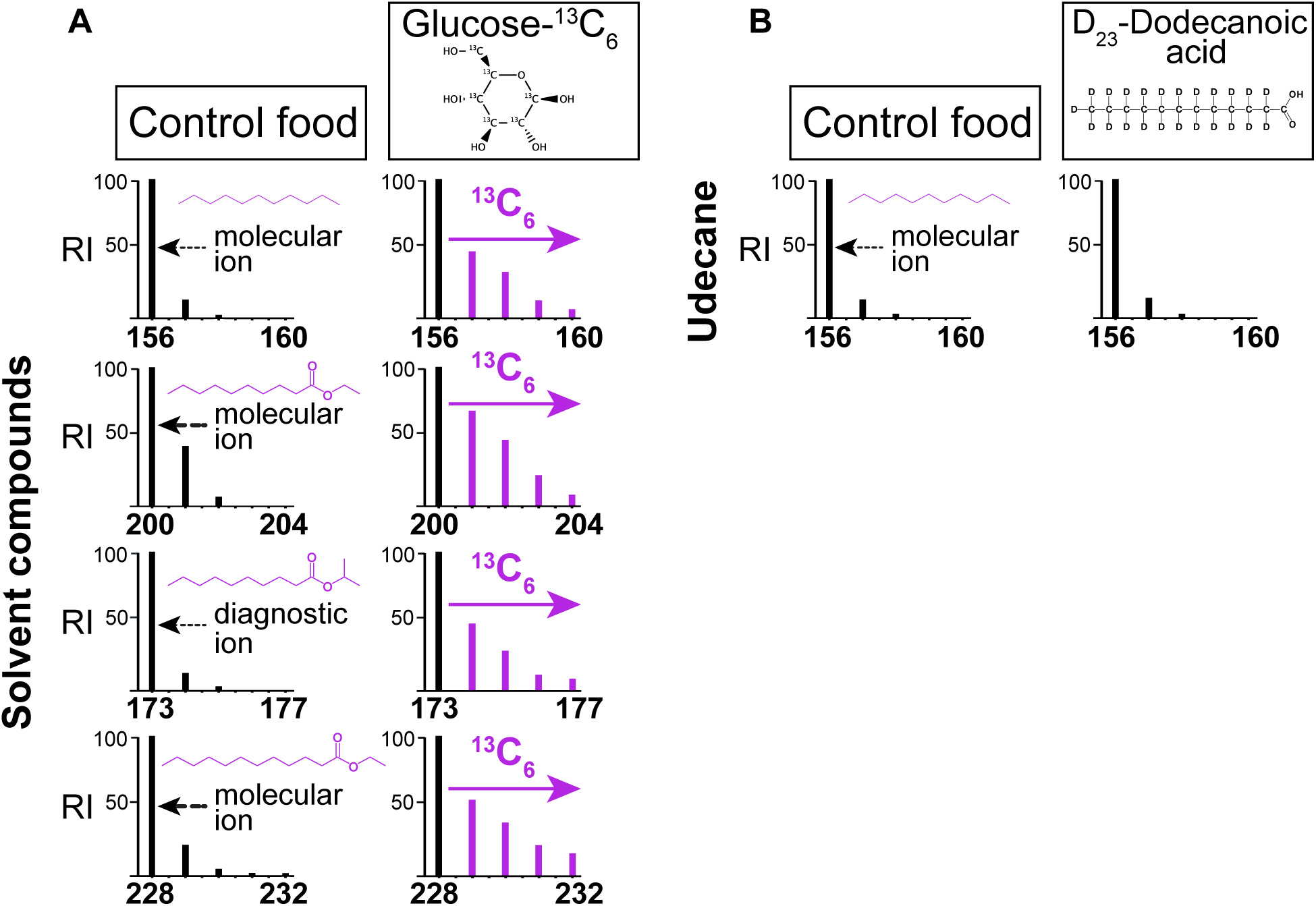
*De novo* undecane and ester biosynthesis from dietary glucose but not dietary fatty acids. **A:** Representative mass spectra of molecular ion region of undecane, ethyl decanoate, isopropyl decanoate and ethyl dodecanoate from beetles fed with unlabeled *Dalotia* diet (control) or infused with D-^13^C_6_-glucose. Spectra recorded in single-ion mode; enriched ions were detected in all four spectra in beetles fed D-^13^C_6_-glucose. **B:** Representative mass spectra of molecular ion region of undecane from beetles fed with unlabeled *Dalotia* diet (control) or food infused with D_23_-dodecanoic acid. Spectra recorded in single-ion mode; no enriched ions were detected.

**Figure S7.**
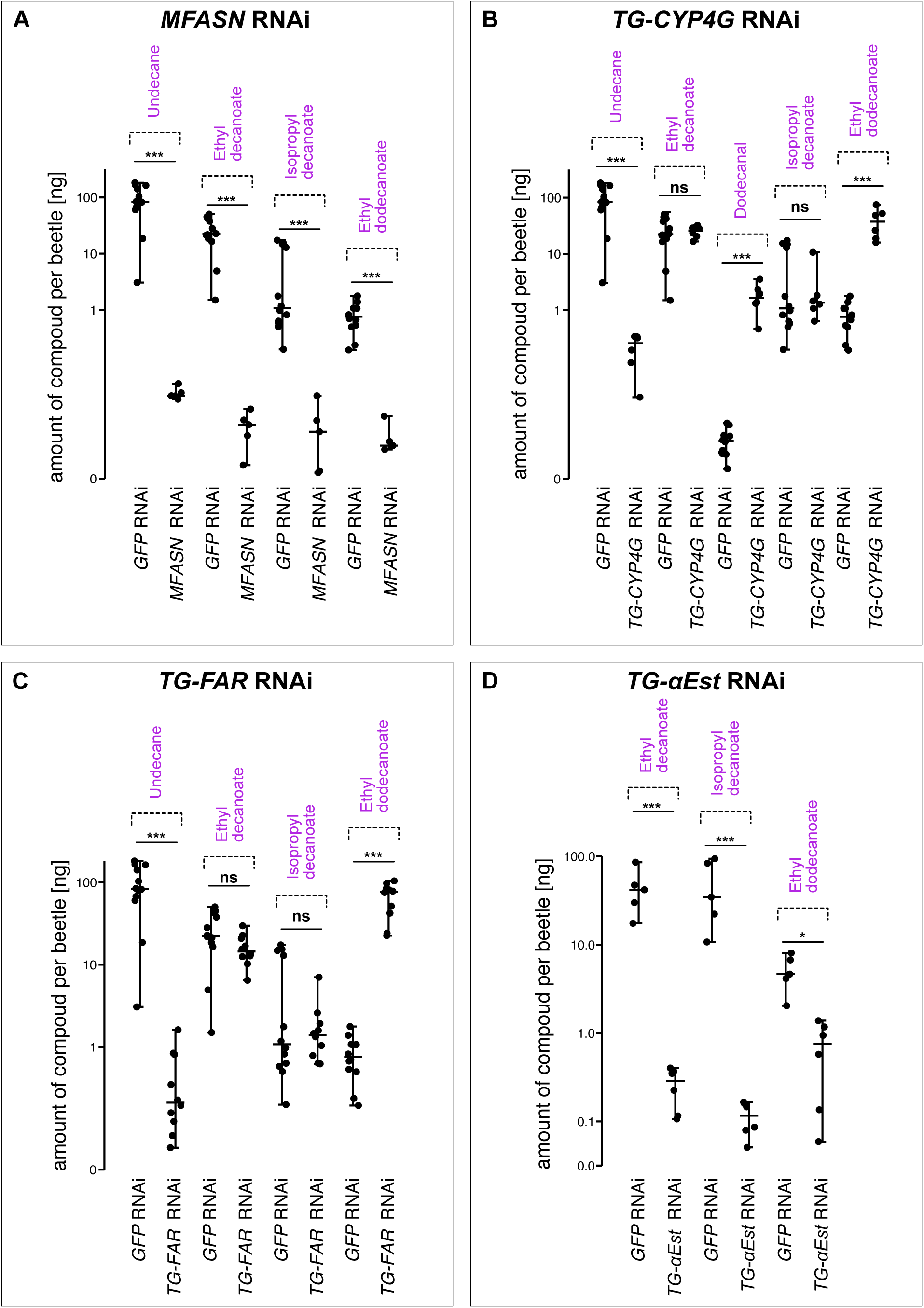
RNAi silencing of solvent pathway enzymes. **A:** Amounts of undecane, ethyl decanoate, isopropyl decanoate and ethyl dodecanoate extracted from beetles injected with dsRNA against *GFP* (control) and *MFASN*. Silencing expression of *MFASN* resulted in near-complete loss of solvents (undecane and esters). **B:** Amounts of undecane, ethyl decanoate, dodecanal, isopropyl decanoate and ethyl dodecanoate extracted from beetles injected with dsRNA against *GFP* (control) and *TG-CYP4G*. Silencing *TG-CYP4G* caused loss of undecane, but a large increase in both dodecanal and ethyl dodecanoate without influencing levels of ethyl decanoate and isopropyl decanoate. **C:** Amounts of undecane, ethyl decanoate, isopropyl decanoate and ethyl dodecanoate extracted from beetles injected dsRNA against *GFP* (control) and *TG-FAR*. Silencing TG-FAR resulted in loss of undecane but a large increase in ethyl dodecanoate without influencing levels of ethyl decanoate and isopropyl decanoate. **D:** Amounts of ethyl decanoate, isopropyl decanoate and ethyl dodecanoate extracted from beetles injected with dsRNA against *GFP* (control) and *TG-αEst*. Silencing TG-αEst resulted in near complete loss of all esters. In all panels, significance was assessed using Mann-Whitney-U tests (***= p≤ 0.001; *= p≤ 0.05; ns= p≥ 0.05).

**Figure S8.**
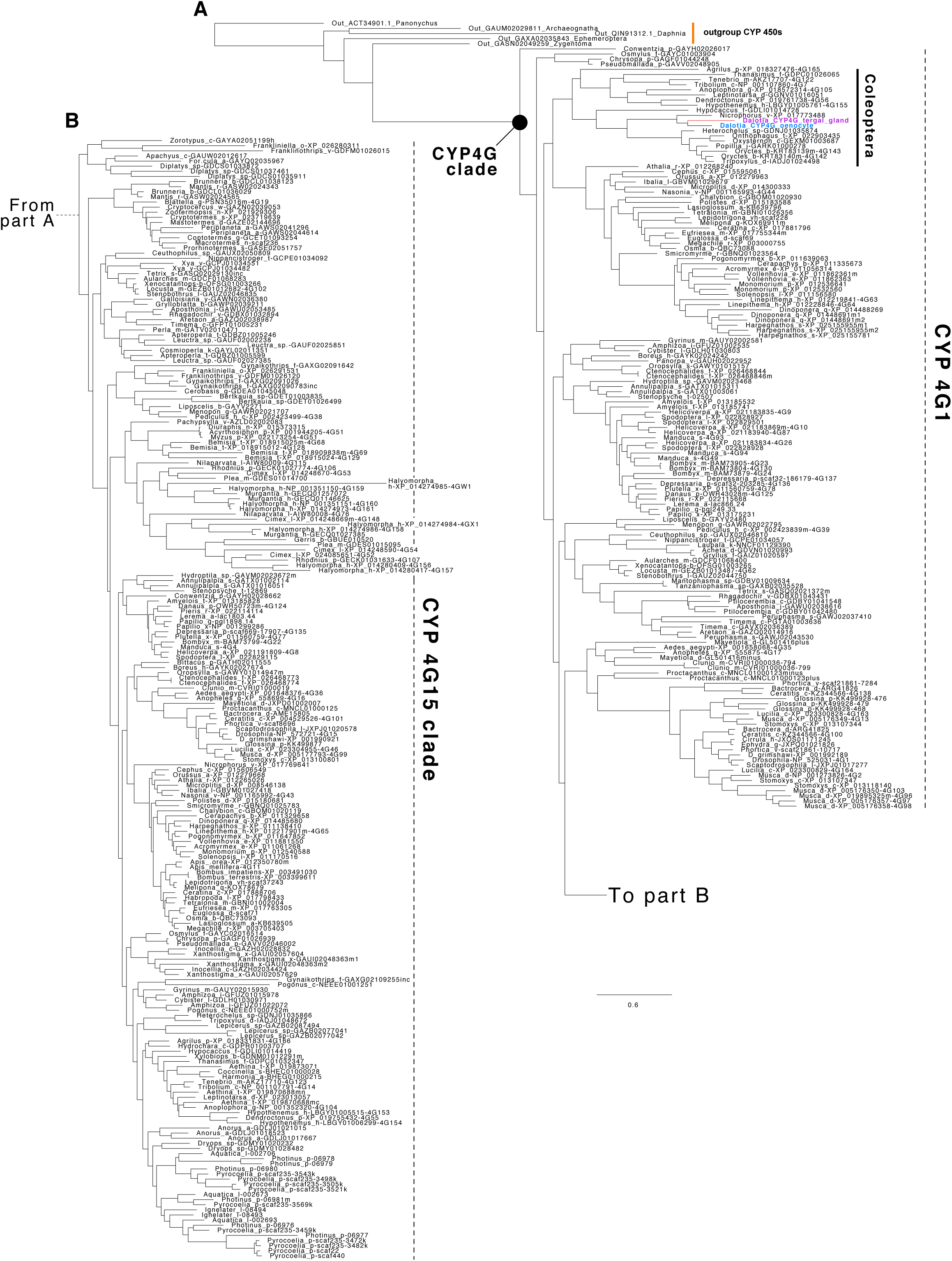
Maximum-likelihood phylogeny of the CYP4G gene family. Part A includes outgroups and the CYP4G1-clade, while part B depicts the CYP4G15 clade. The topology confirms that both *Dalotia CYP4G* copies belong to the CYP4G1 clade, known for their role in cuticular hydrocarbon synthesis in insects.

**Figure S9.**
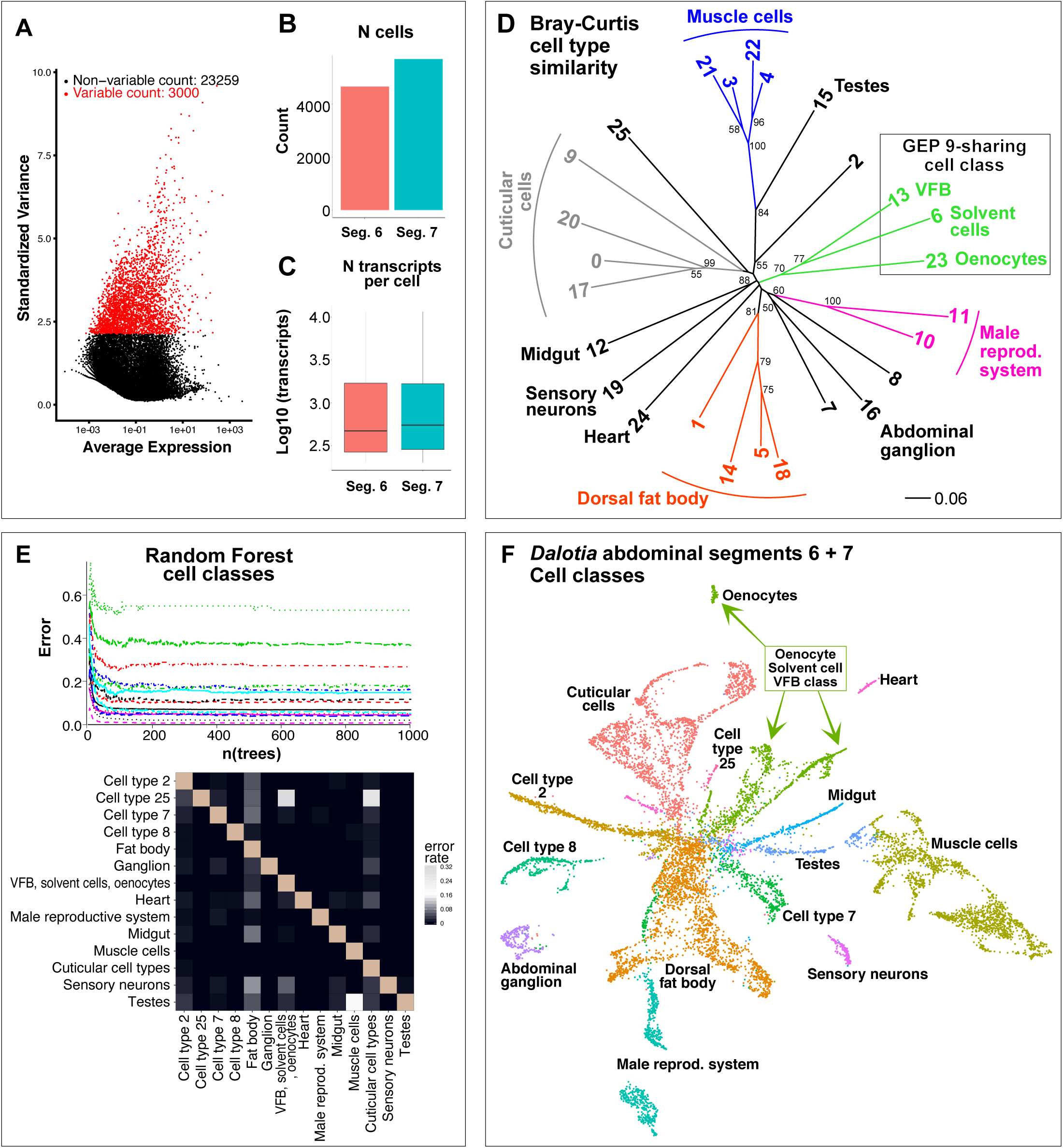
scRNAseq data processing and cell. **A:** Variable feature plot showing the 3,000 transcripts that exhibit high cell-to-cell variation across the dataset (i.e., highly expressed in some cells and not others). These transcripts were used for all downstream analyses and clustering. **B, C:** The total number of cells (**B**) and number of transcripts per cell (**C**) from Segment 6 (control) and 7 (gland) of all concatenated 10x runs after filtering. **D:** Neighbor joining clustering (NJ) on a Bray-Curtis similarity matrix (n= 10,000 bootstraps) based on the normalized and scaled cluster average expression profiles of the 3000 most variable transcripts (**A**). Clades form the basis of further grouping the cell types into 14 higher order cell classes. **E:** Robustness of the 14 cell class assignments tested using a Random Forest classifier. The graph (top) shows the robustness (class error rate) of the cell class assignments running a Random Forest classifier over 1000 generations. The heat map shows the pairwise-class error rates of the assigned groups after 1000 generations. Only cell classes with a relatively low cell number were observed to have a higher error rate. **F:** UMAP plot of *Dalotia* abdominal Segment 6 and 7 cell atlas showing 14 cell classes.

**Figure S10.**
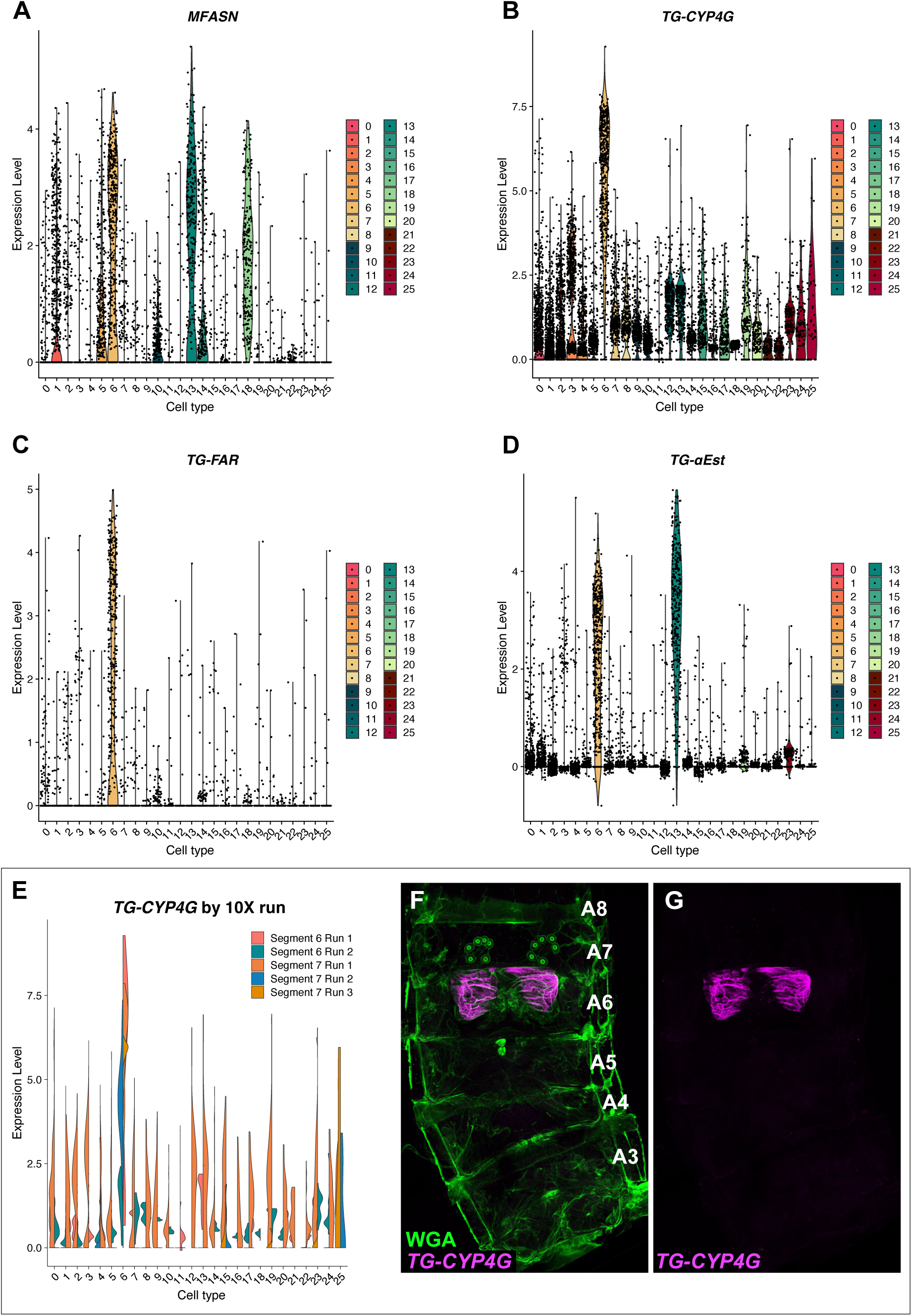
Solvent pathway enzyme expression across cell types. **A-D:** Violin plots showing expression of *MFASN* (**A**), *TG-CYP4G* (**B**), *TG-FAR* (**C**) and *TG-αEST* (**D**) in each cell across all cell types in the abdominal segment cell type atlas. **E:** Expression of *TG-CYP4G* by cell type in each of the five segment-specific 10x runs. *TG-CYP4G* shows a generally high expression across all cell types, but almost all expression in non-solvent cells comes from Segment 6 Run 1 and Segment 7 Run 1. This global expression, with very low levels in almost all cell types, is likely the result of the ‘ambient mRNA’-effect, which is typical of droplet-based methods. Ambient mRNAs are most likely derived from high-abundance transcripts that are leaked from dying cells and bind to beads during the microfluidic step in the 10x workflow. Only in solvent cells are the UMI count levels were high, indicating that the low levels mRNA detected in non-solvent cells likely represent ambient mRNAs. **F, G:** To independently evaluate the cell type specificity of *TG-CYP4G*, we used HCR and imaged the whole beetle abdomen, confirming that its transcription is highly specific to the solvent cells.

**Figure S11.**
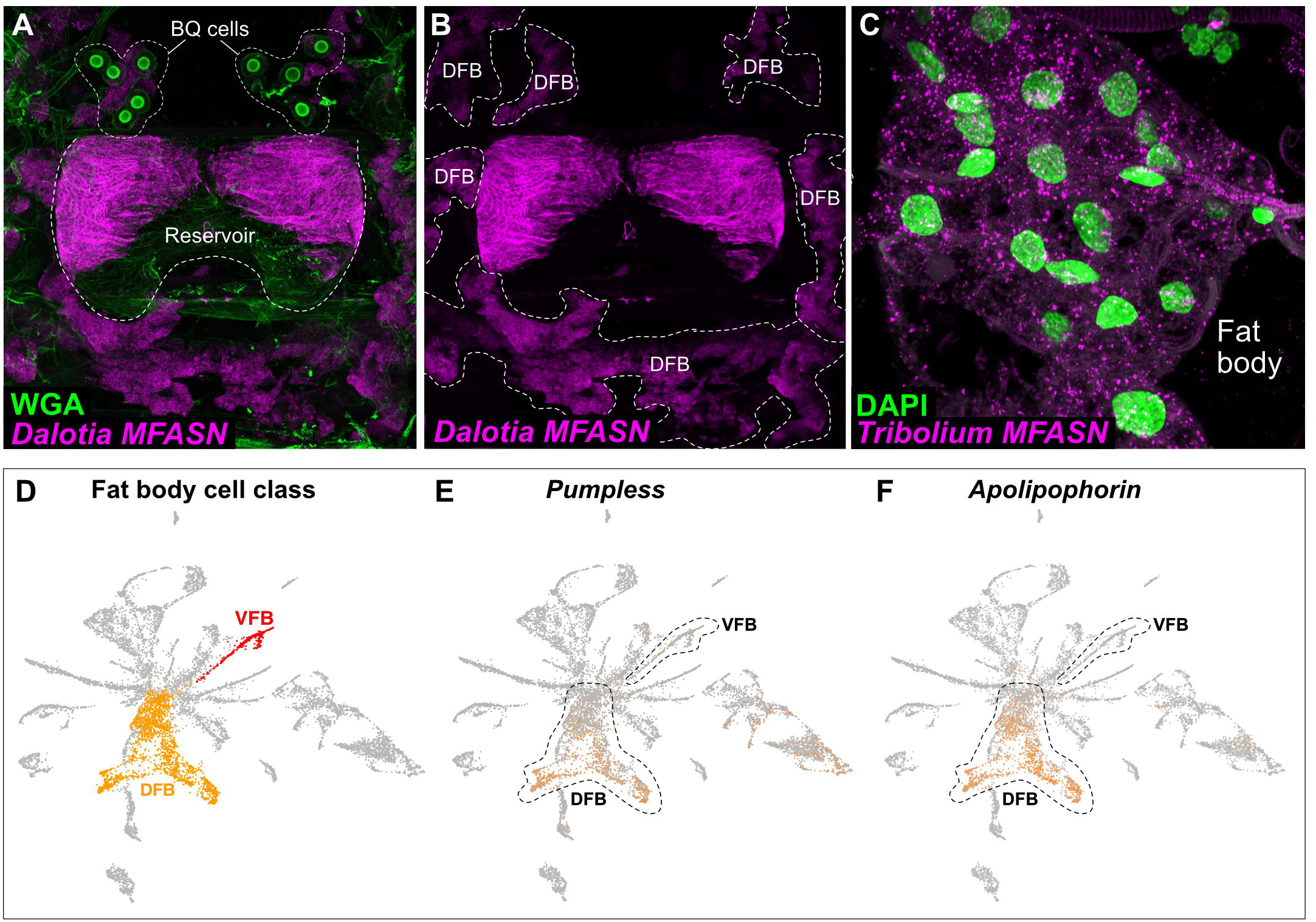
Expression of *MFASN* in *Dalotia* and *Tribolium* and fat body marker genes in VFB and DFB. **A-C:** HCR in *Dalotia* reveals *MFASN* in the solvent cells (**A**) and dorsal fat body (DFB; **B**). **C:** HCR of the *MFASN* orthologue in *T. castaneum* reveals strong expression in fat body **D:** UMAP plot of *Dalotia* abdominal segment 6 and 7 cell atlas highlighting the dorsal (DFB; orange) and ventral fat body (VFB; red) cells. **E, F:** Expression of the *Pumpless* (**E**) and *Apolipophorin* (**F**) fat body marker genes is strong expression in DFB but not VFB.

**Figure S12.**
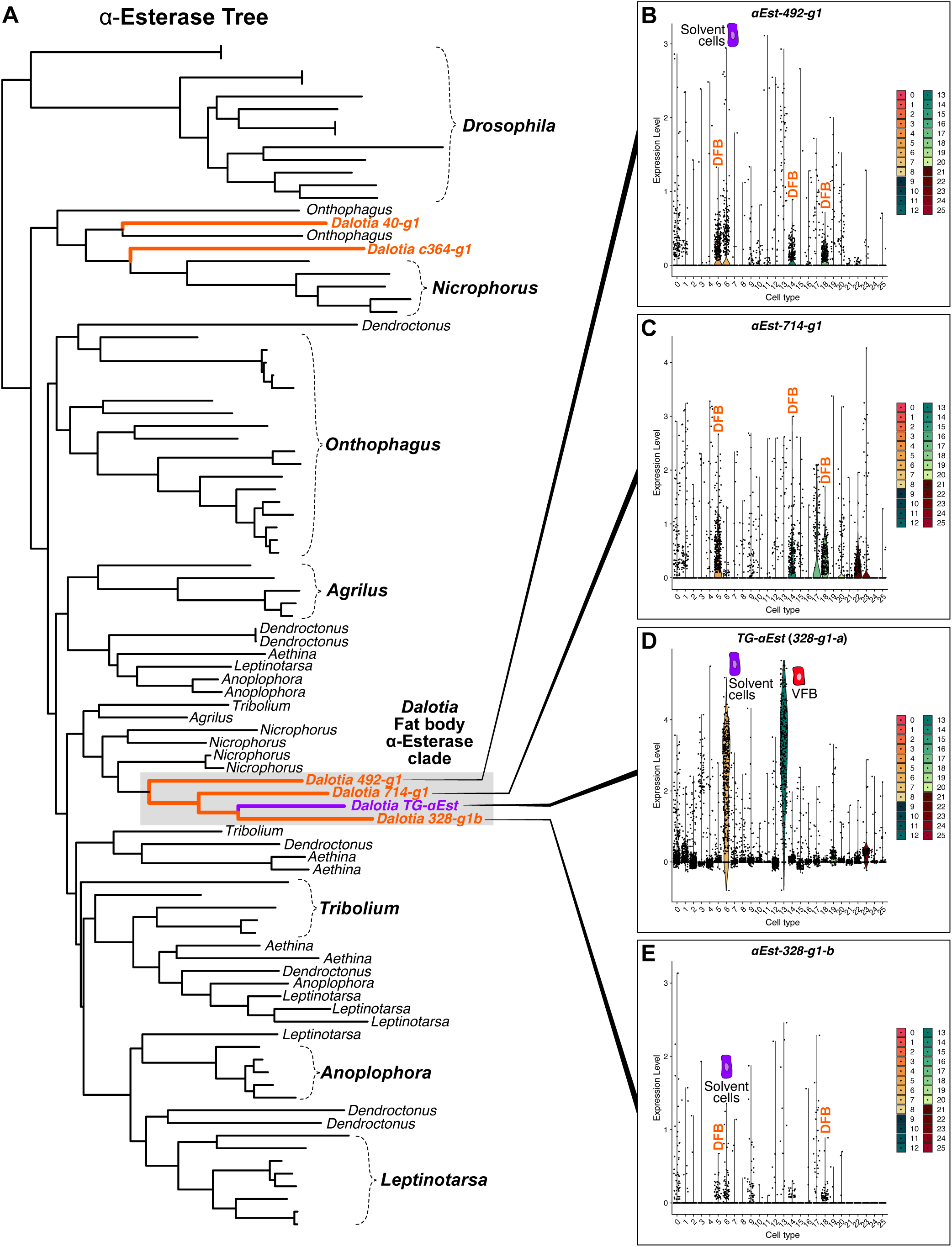
Phylogenetic tree and cell type expression of α-esterases. **A:** Maximum-likelihood tree of the α-esterases, including the six transcripts from *Dalotia*. The topology shows that *TG-αEst* belongs to a *Dalotia*-specific clade. **B-E:** Violin dot plots showing the expression of the *Dalotia* α-esterase clade transcripts across *Dalotia* abdominal cell types. While *TG-αEst* (**D**) shows high expression in the solvent cells and VFB, the three other α-esterases (**B**, **C** and **E**) show low to medium expression primarily in DFB.

**Figure S13.**
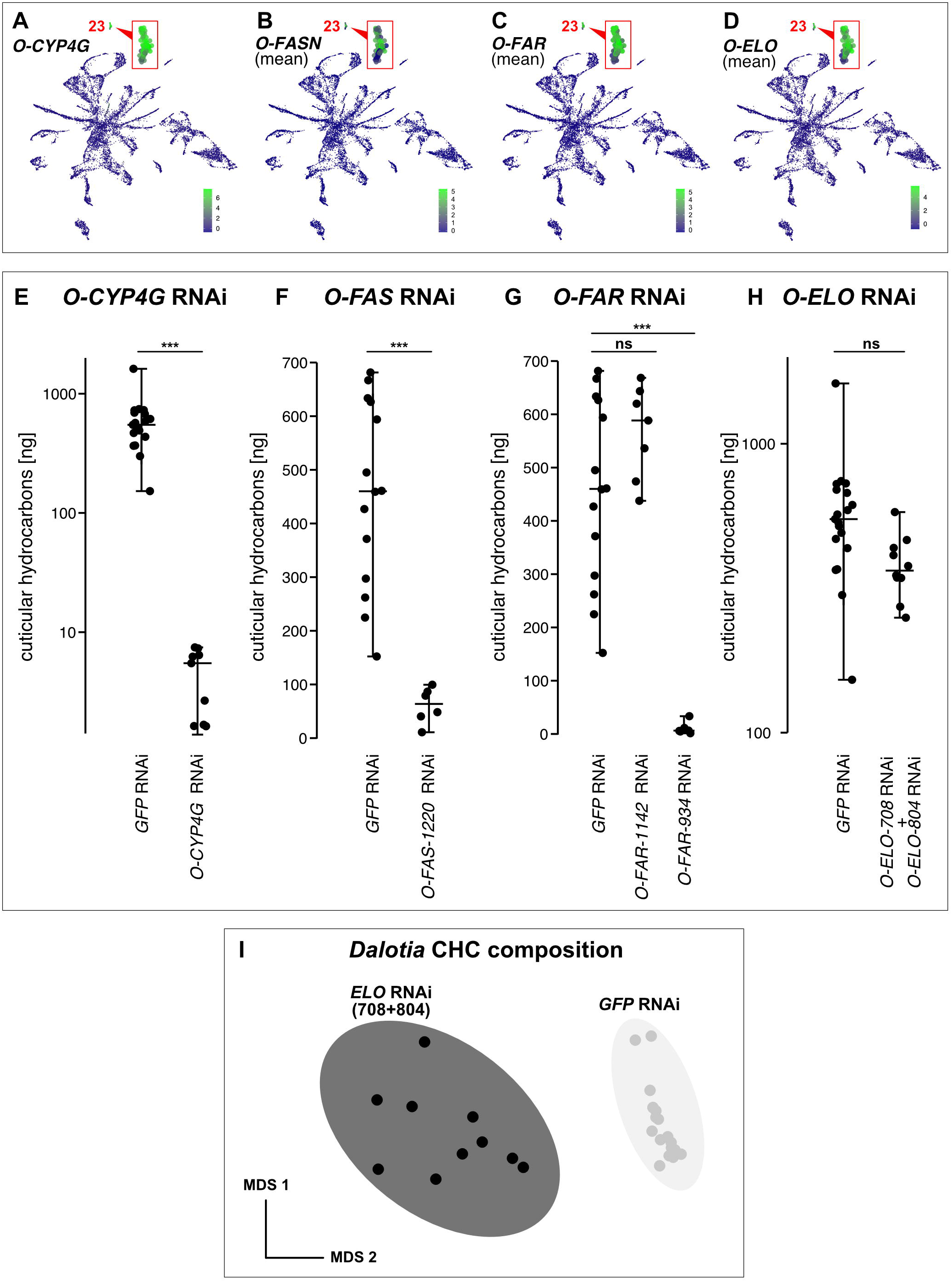
RNAi silencing of CHC pathway enzymes in oenocytes. **A-D:** UMAPs showing expression of *O-CYP4G* (**A**), *O-FASN*s (**B**), *O-FAR*s (**C**) and *O-ELO*s (**D**) in oenocytes (insets show the oenocyte cell cluster magnified). Note that for *FASN*s, *FAR*s and *ELO*s, multiple copies are expressed, and the mean expression of all copies is shown. **E-G:** Levels of cuticular hydrocarbons (CHCs) extracted from beetles injected with dsRNA against *GFP* (control) and *O-CYP4G* (**E**), *O-FAS-1220* (**F**), *O-FAR-1142* and *O-FAR-934* (**G**) Silencing *O-CYP4G* or *O-FAS-1220* results in near-complete loss of CHCs on *Dalotia*’s body surface. Silencing *O-FAR-1142* did not affect CHC levels, while *O-FAR-934* RNAi resulted in CHC loss. **H:** Silencing expression of *O-ELO-708* and *O-ELO-804* did not affect CHC levels (**H**) but shifted the composition of CHCs found on *Dalotia*’s cuticle surface towards shorter-chained CHCs (**I**). Significance in all panels was assessed using Mann-Whitney-U tests or permutative t-tests (***= p≤ 0.001; ns= p≥ 0.05).

**Figure S14:**
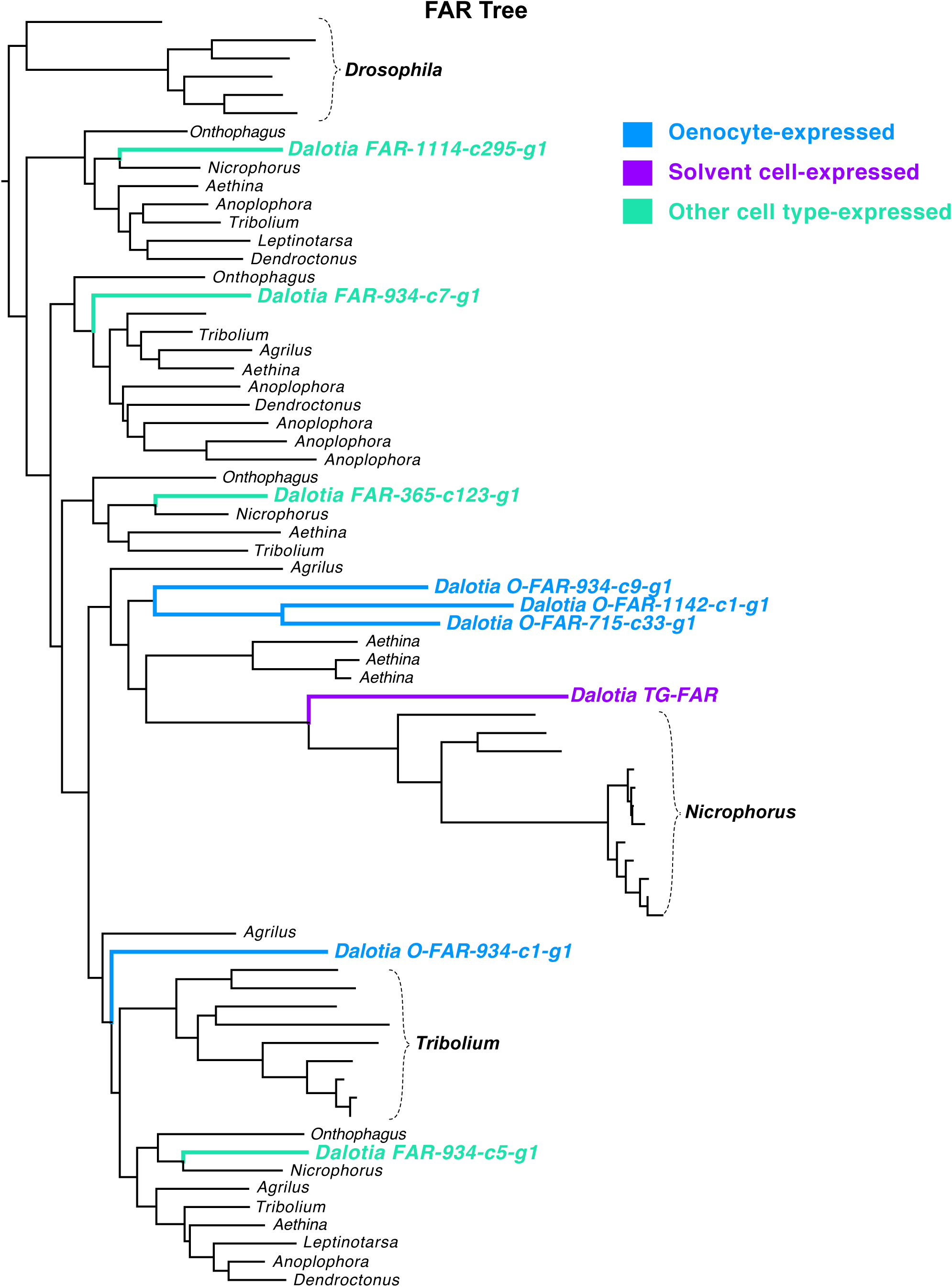
Maximum-likelihood tree of fatty acyl-CoA reductase (FAR) transcripts in *Dalotia* and other insects. The tree shown was rooted with the *Drosophila* FAR encoded by *waterproof*. *Dalotia* sequences in the tree are color-coded by their tissue specific expression.

**Figure S15.**
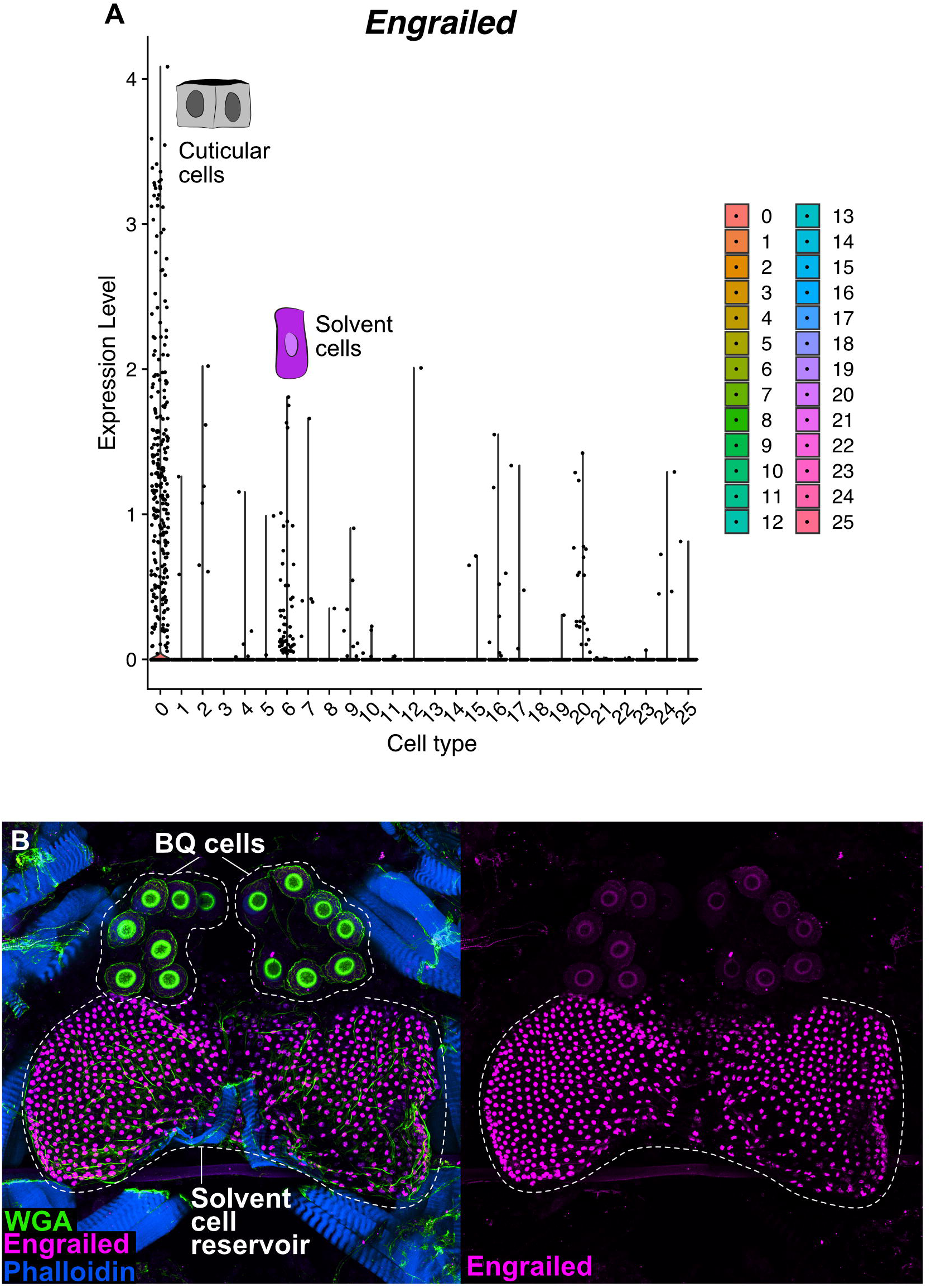
Solvent cells express *engrailed*. **A:** Violin dot plot showing tissue-specific expression of the *en* in the *Dalotia* abdominal cell atlas. Expression in *en* in adults is largely restricted to cuticular cells and the solvent cells. **B:** Immunohistochemistry shows strong labeling of solvent cells with En antibody.

**Figure S16.**
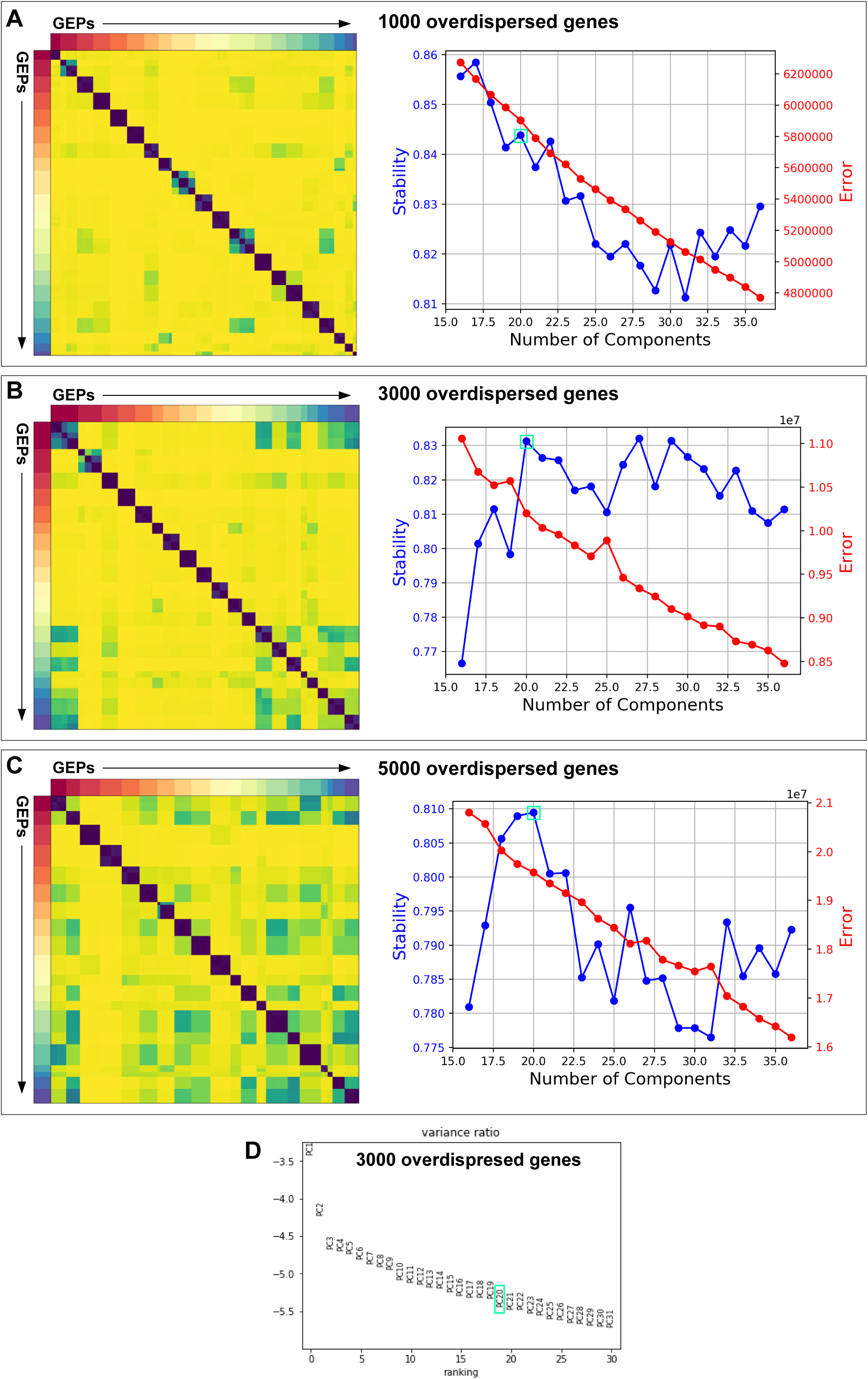
Diagnostic plots for K selection in cNMF. **A:** Clustergram showing the clustered NMF components for K = 20, combined across 250 replicates, after filtering (left). Number of cNMF components (K) against stability (blue) i.e., the silhouette score of the clustering, and Frobenius error (red) for a data set using 1000 overdispersed genes. **B:** Clustergram showing the clustered NMF components for K = 20, combined across 250 replicates, after filtering (left). Number of cNMF components (K) against stability (blue) i.e., the silhouette score of the clustering, and Frobenius error (red) for a data set using 3000 overdispersed genes. **C:** Clustergram showing the clustered NMF components for K = 20, combined across 250 replicates, after filtering (left). Number of cNMF components (K) against stability (blue) i.e., the silhouette score of the clustering, and Frobenius error (red) for a data set using 5000 overdispersed genes. **D:** Plot of the Log10 proportion of variance explained per principal component for a data set using 3000 overdispersed genes. The K selection used in the manuscript is indicated by a turquoise box.

**Figure S17.**
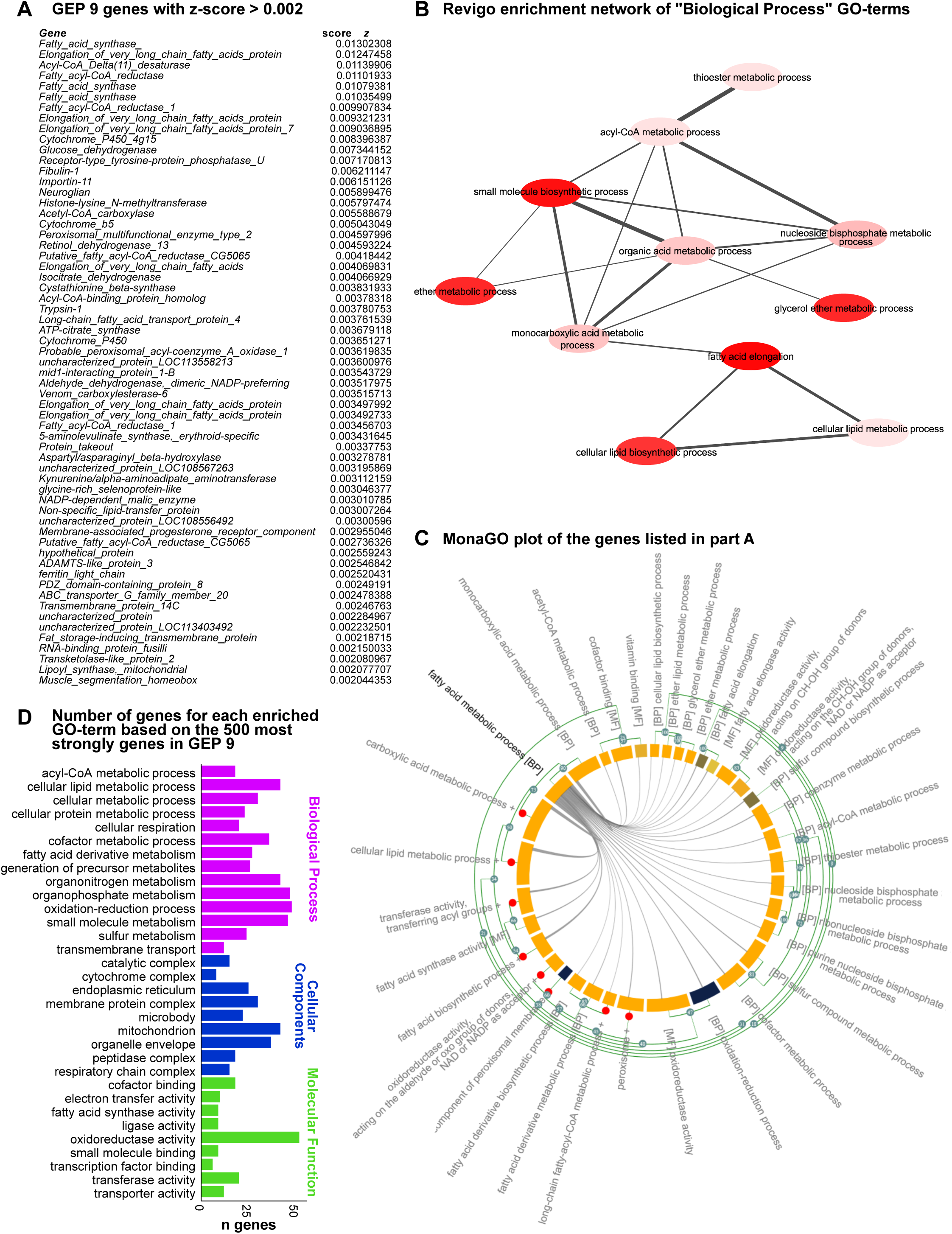
GEP 9 is a fatty acid metabolism transcriptional module. **A:** List of all genes with a z-score > 0.002, denoting a strong association with GEP9. A majority of genes is either related to fatty acid synthesis or modification of fatty acid derivatives. **B:** Gene ontology (GO) term enrichment network of the ‘Biological Process’ category based on the list of genes in **A**. Disc color corresponds to the significance (as assessed by Fisher’s exact test) of the respective GO-terms, while the thickness of the grey lines represents the semantic similarities between the GO-terms. Network created using REViGO. **C:** MonaGO plot of the genes listed in **A**. The inner cord plot shows which genes of the GO-term ‘fatty acid metabolism’ are shared with other GO-terms. The inner ring displays the significance (as assessed by Fisher’s exact test) of enrichment of each GO-term, while the outer cluster dendrogram is based on the pairwise similarity (Jaccard distance) of each GO-term. Collapsed clusters are depicted as red dots. BP= biological process, MF= molecular function. Plot created with MonaGO. **D:** Number of genes for each enriched GO-term (as assessed by Fisher’s exact test) for all three GO-categories based on the top 500 genes (by z-score) of GEP 9.

**Figure S18.**
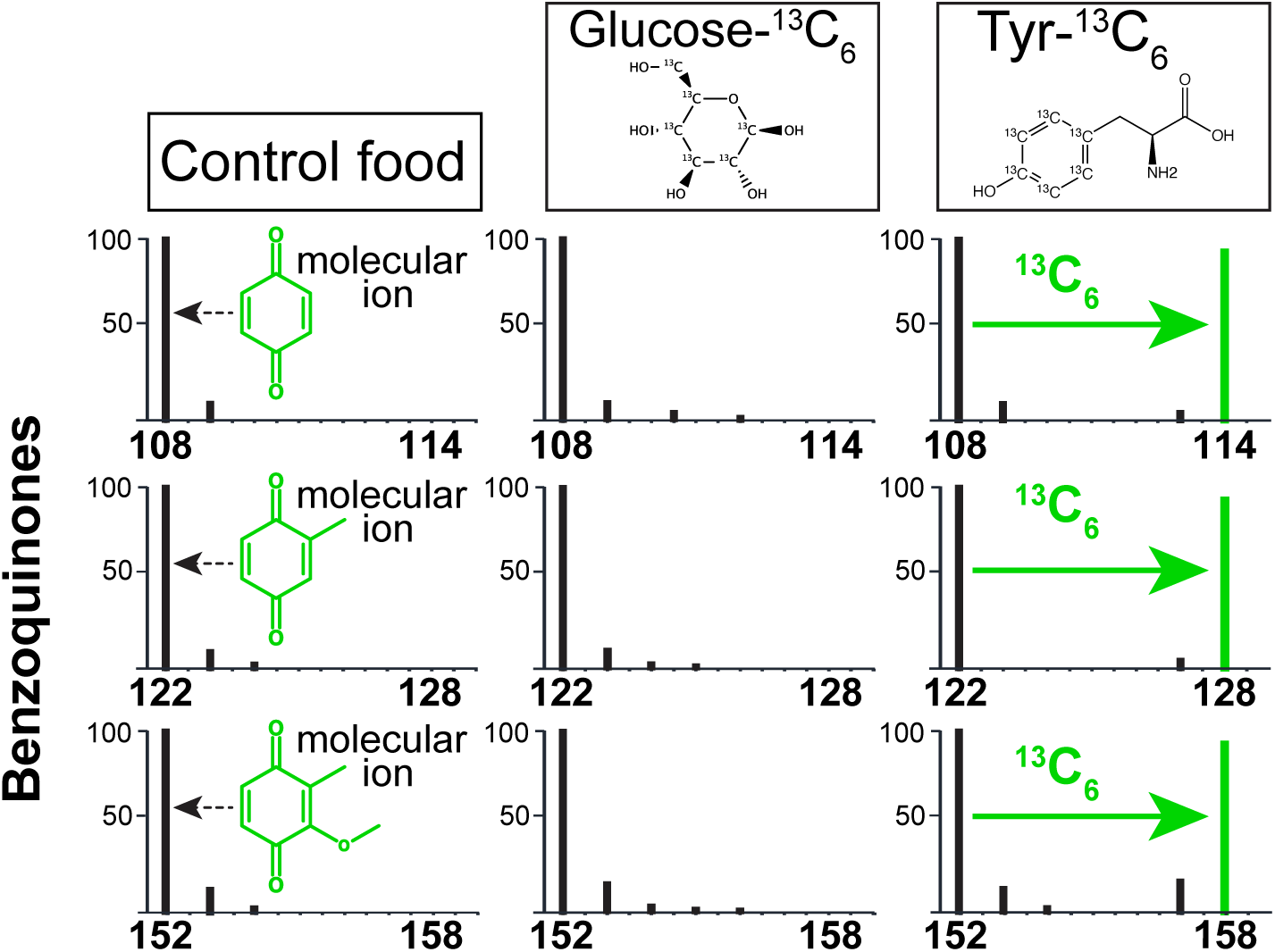
BQs are synthesized from dietary Tyr but not from glucose. Representative mass spectra of molecular ion region of 1,4-BQ, 2-methl-1,4-BQ and 2-methoxy-3-methyl-1,4,-BQ from beetles fed with unlabeled *Dalotia* diet (control), food infused with ^13^C_6_-glucose or ^13^C_6_-Tyr. Spectra were recorded in single-ion mode. No enrichment was detected in any of the three BQs for the ^13^C_6_-glucose fed beetles. In contrast, a strong [M+6]^+^ enrichment was detected for all BQs in ^13^C_6_-Tyr fed beetles.

**Figure S19.**
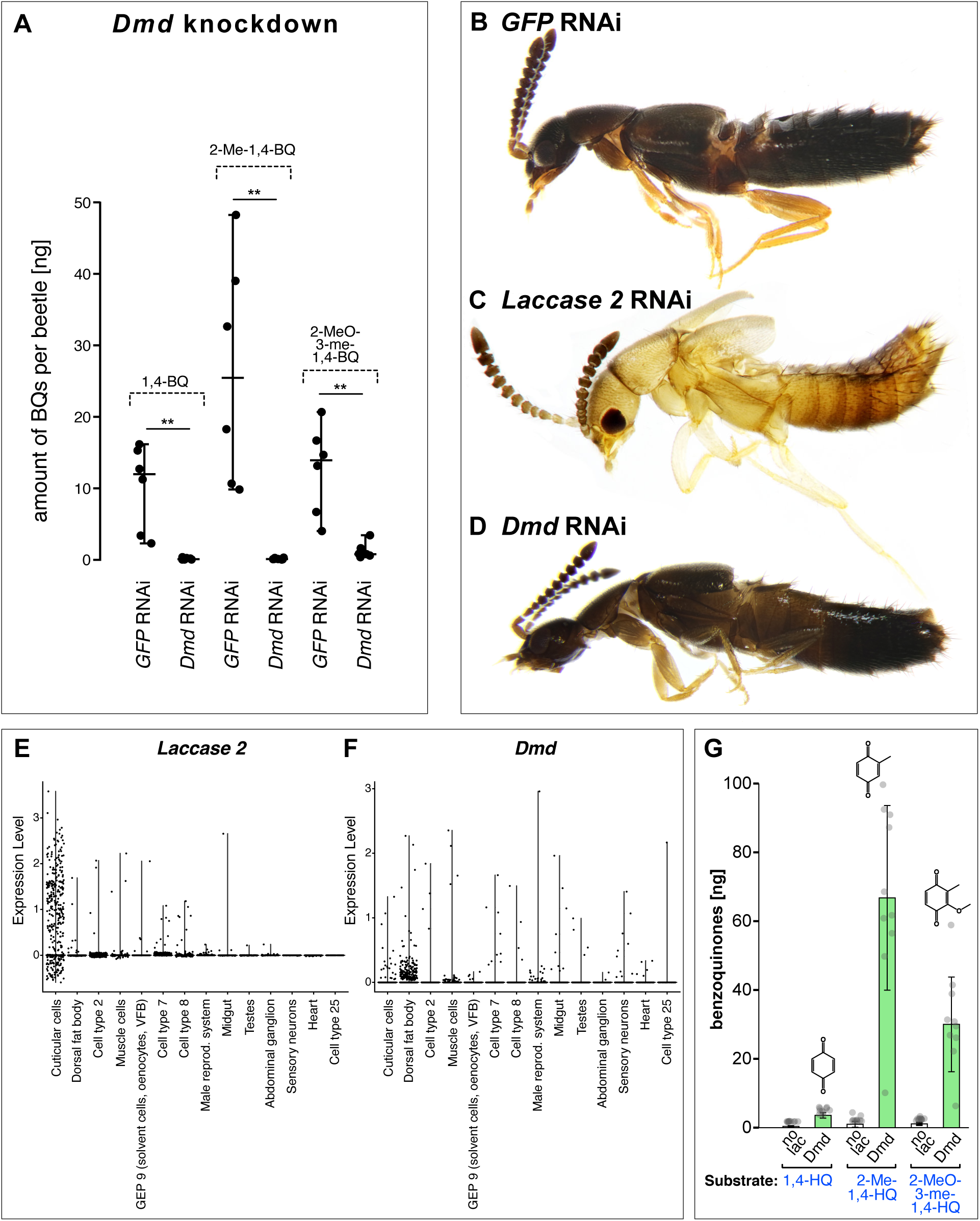
Functional characterization of Dmd. **A:** Levels of 1,4-BQ, 2-methyl-1,4-BQ and 2-methoxy-3-methyl-1,4,-BQ extracted from beetles injected with dsRNA against *GFP* (control) and *Dmd*. Silencing *Dmd* resulted in near-complete loss of BQs in *Dalotia*’s secretion. **B-D:** Adult beetle morphology after late larval injection with dsRNA against *GFP* (**B**; control), *Lac2* (**C**) and *Dmd* (**D**). **E**: Expression of *Laccase-2* across all cell classes in the 10x data reveals strong expression cuticle cells. **F**: Expression of *Dmd* across all cell classes in the 10x data shows no tissue specific expression of *Dmd* as BQ-cells are not captured by 10x. G: Reaction of the three HQ substrates 1,4-HQ, 2-Me-1,4-HQ and 2-MeO-3-Me-1,4-HQ with purified Dmd *in vitro* as compared to a non-enzyme control. BQ levels were measured using GC-FID.

**Figure S20.**
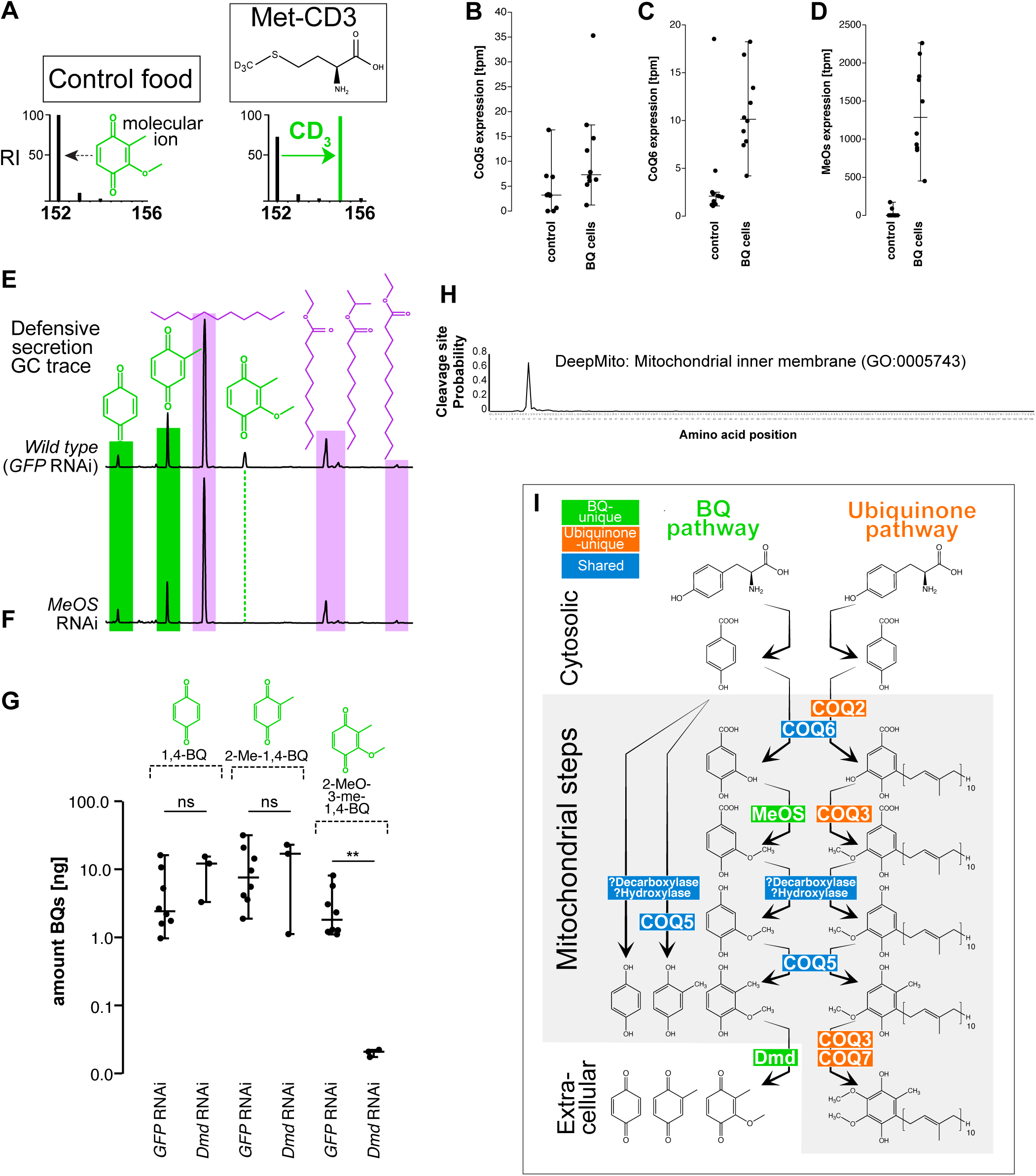
Evidence for a mitochondrial HQ pathway. **A:** Representative mass spectra of molecular ion region of 2-methoxy-3-methyl-1,4-BQ from beetles fed with unlabeled *Dalotia* diet (control) versus food infused with CD_3_-methionine. Spectra were recorded in single-ion mode. A strong [M+3]^+^ enrichment indicates that the methoxy group is derived from dietary Met. **B-D:** Expression of *CoQ5* (**B**), *CoQ6* (C) and *MeOS* (**D**) in BQ cells and control tissue based on SMARTseq. **E, F:** Representative GC traces of gland compounds from wild type (*GFP* RNAi; **E**) and *MeOS* RNAi-injected (**F**) beetles. Silencing of MeOS leads to loss of 2-methoxy-3-methyl-1,4-BQ only, while both other BQs are still present. **G:** Amounts of 1,4-BQ, 2-methyl-1,4-BQ and 2-methoxy-3-methyl-1,4,-BQ extracted from *GFP* RNAi (control) and MeOS *RNAi* beetles. Significance was assessed using Mann-Whitney-U tests (**= p≤ 0.01). **H:** MeOS contains a mitochondrial inner membrane (GO:0005743) signal peptide sequence. **I:** Proposed biosynthesis pathway of *Dalotia*’s BQs versus the mitochondrial ubiquinone pathway. Unique BQ pathway enzymes are in green; unique ubiquinone pathway enzymes are in orange; enzymes shared by the two pathways are in blue.

**Figure S21.**
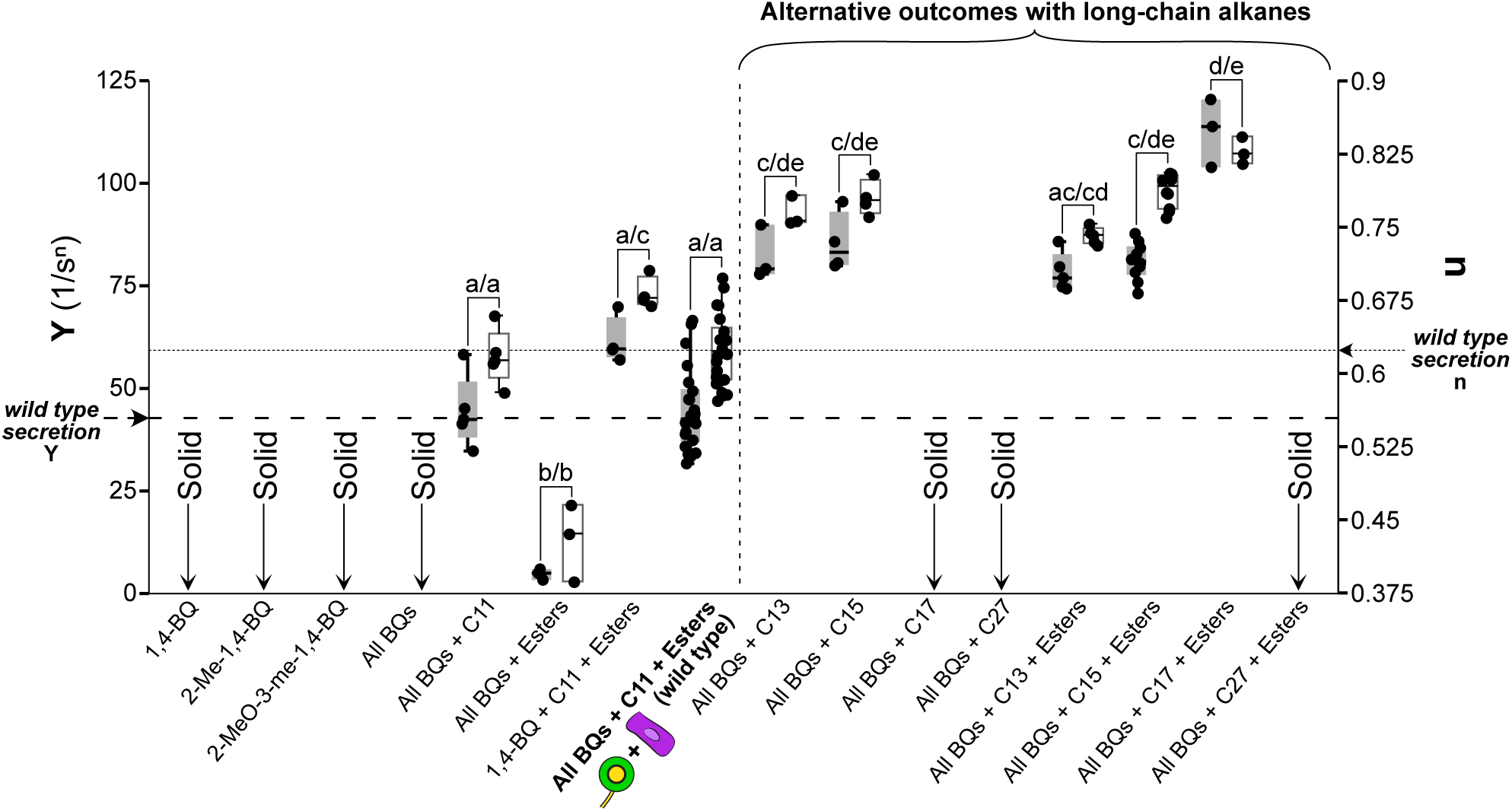
Extensional viscosities (EV) of synthetic secretions. Boxes in grey show data for the elongational-flow consistency index Y (left axis), while the white boxes correspond to the flow behavior index n (right axis). The medians for Y and n of the wild-type mixtures are shown as horizontal lines. Lower values of both Y and n together correspond to higher EV. Different letters indicate significant difference in Y or n across test mixtures as assessed by a Tukey HSD posthoc test following a one-way ANOVA.

**Figure S22.**
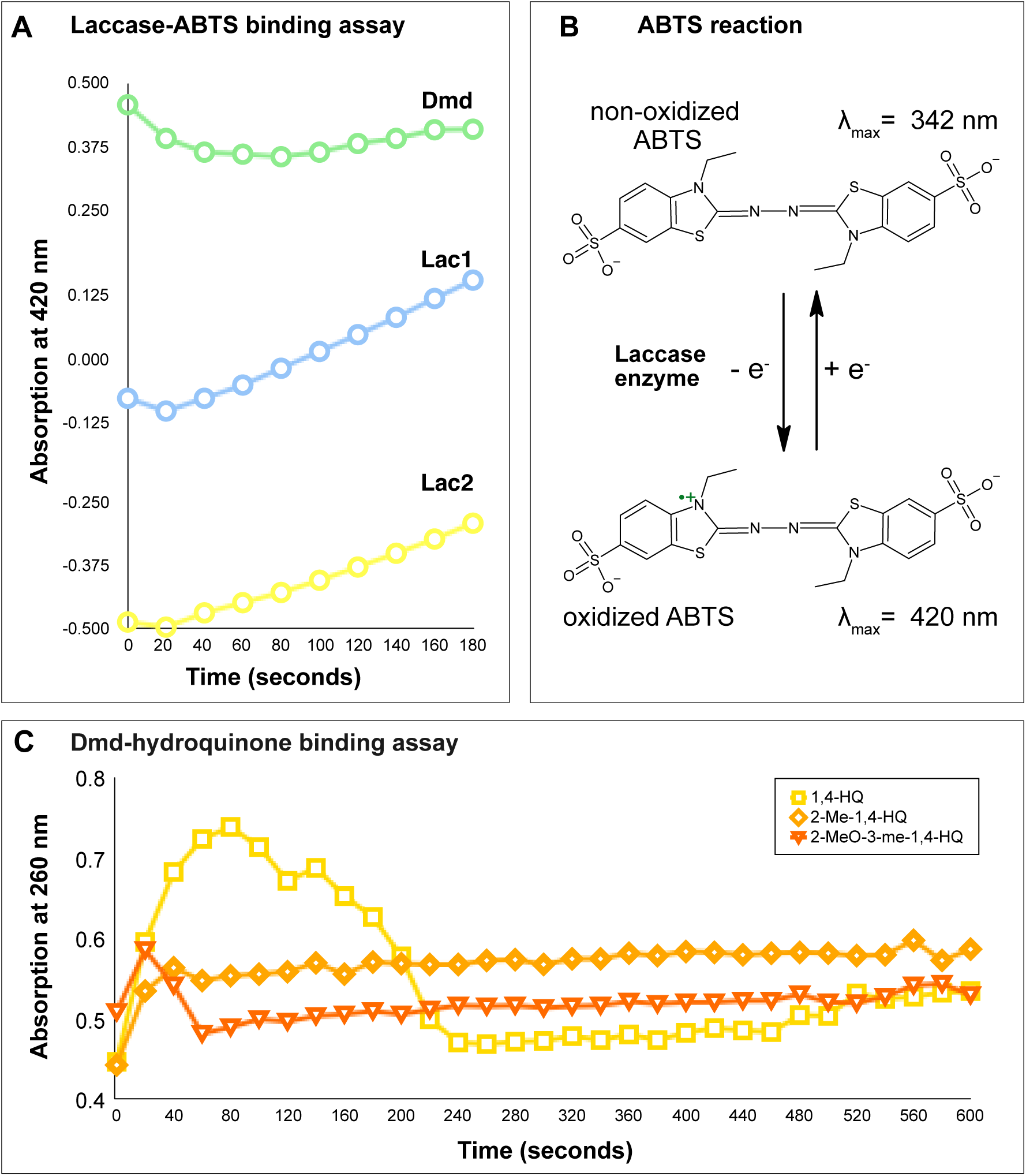
*Dalotia* laccase *in vitro* binding assays. **A:** Absorption measurements at λ= 420 nm for Dmd, Lac1 and Lac2 over 180 s using the Laccase-ABTS binding assay confirms that all three *in vitro*-expressed laccase enzymes are vital and bind to a control substrate. **B:** Scheme showing reaction of ABTS with laccase enzymes used to measure absorption in part **A**. One tertiary N of ABTS (λ_max_= 342 nm) is oxidized by laccase, forming the oxidized N•+ radical with a λ_max_ at 420 nm while reducing Cu^+2^ to Cu^1+^. **C:** Monitoring absorption at λ= 260 nm for the *in vitro* reaction of Dmd with 1,4-HQ, 2-methl-1,4-HQ and 2-methoxy-3-methyl-1,4-HQ over 600 s confirms that Dmd catalyzes oxidation of all three substrates to the respective BQs.

## Supplementary tables

**Table S1. Statistical analyses of all experimental data presented in the main and supplementary figures.** For each analysis the comparison is described, sample sizes are given, and statistical methods are named; p-values or other statistical values are listed for each test. All tests were performed with Past 3.04.

