## Supplemental Table S1 for "Evolutionary assembly of cooperating cell types in an animal chemical defense system"

| **Figure 1** | **Comparison** | **N number** | **methods** | **P value** | **statistical values** |
| --- | --- | --- | --- | --- | --- |
| 1C  (not shown) | Percentage of surviving *Dalotia* with in an arena with aggressive ants after 48h with either armed (control) or disarmed tergal glands  Amount (µg) of total gland content armed (control) vs disarmed (CO_2_ treatment) beetles | 14 vs 14 groups of 10 beetles  25 control vs 39 disarmed beetles | Mann-Whitney U test  Mann-Whitney U test | < 0.0001  < 0.0001 | z= 4.5, N= 28  z= 6.7, N= 64 |
| **Figure 2** | **Comparison** | **N number** | **methods** | **P value** | **statistical values** |
| 2B | Amount (ng) of undecane in GFP vs MFASN RNAi beetles  Amount (ng) of ethyl decanoate in GFP vs MFASN RNAi beetles  Amount (ng) of isopropyl decanoate in GFP vs MFASN RNAi beetles  Amount (ng) of ethyl dodecanote in GFP vs MFASN RNAi beetles | 12 (GFP) vs 5 (RNAi beetles) | Mann-Whitney U tests | 0.00187  0.00187  0.00187  0.00187 | z= 3.1, N= 17  z= 3.1, N= 17  z= 3.1, N= 17  z= 3.1, N= 17 |
| 2C | Amount (ng) of undecane in GFP vs TG-CYP4G RNAi beetles  Amount (ng) of ethyl decanoate in GFP vs TG-CYP4G RNAi beetles  Amount (ng) of dodecanal in GFP vs TG-CYP4G RNAi beetles  Amount (ng) of isopropyl decanoate in GFP vs TG-CYP4G RNAi beetles  Amount (ng) of ethyl dodecanote in GFP vs TG-CYP4G RNAi beetles | 12 (GFP) vs 6 (RNAi beetles) | Mann-Whitney U tests | 0.00088  0.88827  0.00088  0.74306  0.00088 | z= 3.3, N= 18  z= 0.1, N= 18  z= 3.3, N= 18  z= 0.3, N= 18  z= 3.3, N= 18 |
| 2D | Amount (ng) of undecane in GFP vs TG-FAR RNAi beetles  Amount (ng) of ethyl decanoate in GFP vs TG-FAR RNAi beetles  Amount (ng) of isopropyl decanoate in GFP vs TG-FAR RNAi beetles  Amount (ng) of ethyl dodecanote in GFP vs TG-FAR RNAi beetles | 12 (GFP) vs 10 (RNAi beetles) | Mann-Whitney U tests | < 0.0001  0.08058  0.92121  < 0.0001 | z= 3.9, N= 22  z= 1.7, N= 22  z= 0.1, N= 22  z= 3.9, N= 22 |
| 2E | Amount (ng) of ethyl decanoate in GFP vs TG-aEST RNAi beetles  Amount (ng) of isopropyl decanoate in GFP vs TG-aEST RNAi beetles  Amount (ng) of ethyl dodecanote in GFP vs TG-aEST RNAi beetles | 5 (GFP) vs 6 (RNAi beetles) | Mann-Whitney U tests | 0.00811  0.00811  0.00812 | z= 2.6, N= 11  z= 2.6, N= 11  z= 2.6, N= 11 |
| **Figure 4** | **Comparison** | **N number** | **methods** | **P value** | **statistical values** |
| 4G | Amount (ng) of total CHCs in GFP vs O-FASN RNAi beetles | 14 (GFP) vs 6 (RNAi beetles) | Mann-Whitney U test | 0.00062 | z= 3.4, N= 20 |
| 4H | Amount (ng) of total CHCs in GFP vs O-FAR-1142 RNAi beetles  Amount (ng) of total CHCs in GFP vs O-FAR-934 RNAi beetles (not shown in figure) | 14 (GFP) vs 7 (RNAi beetles for each group) | Mann-Whitney U test  permutation t-test | 0.00029  0.12397 | z= 3.6, N= 21  t_1,19_= 1.6, N=21, n_permute_= 9999 |
| 4I | Amount (ng) of total CHCs in GFP vs O-CYP4G RNAi beetles | 19 (GFP) vs 9 (RNAi beetles) | Mann-Whitney U test | < 0.0001 | z= 4.2, N= 28 |
| 4J | Amount (ng) of total CHCs in GFP vs O-ELOVL RNAi beetles  Compositional changes of CHC in GFP vs O-ELOVL RNAi beetles | 19 (GFP) vs 10 (RNAi beetles) | permutation t-test  PERMANOVA (on a Bray-Curtis similarity matrix) | 0.0631  0.0001 | t_1,27_= 1.8, N=29, n_permute_= 9999  _pseudo_F= 59.2, N= 29, n_permute_= 9999 |
| **Figure 6** | **Comparison** | **N number** | **methods** | **P value** | **statistical values** |
| 6F | Amount (ng) of 1,4-benzoquinone (1,4-BQ) in GFP vs Dmd RNAi beetles  Amount (ng) of 2-methyl-1,4-BQ in GFP vs Dmd RNAi beetles  Amount (ng) of 2-methoxy-3-methyl-1,4-BQ in GFP vs Dmd RNAi beetles | 6 (GFP) vs 7 (RNAi beetles) | Mann-Whitney U tests | 0.00340  0.00340  0.00340 | z= 2.9, N= 13  z= 2.9, N= 13  z= 2.9, N= 13 |
| 6G | Comparison of 2-methyl-1,4-HQ amounts (ng) in GFP vs Dmd RNAi beetles monitored by single-ion mode (SIM) | 5 (GFP) vs 6 (RNAi beetles) | Mann-Whitney U test | 0.03030 | z= 2.1, N= 11 |
| 6H | Laccase comparison assay - amount (ng) of 1,4-BQ produced after inoculation of 1,4-HQ with Lac1, Lac2, Dmd and no-enzyme control  Pairwise comparisons of Dmd vs Lac1/2 and control  Pairwise comparisons among Lac1/2 and control  Laccase comparison assay - amount (ng) of 2-methyl-1,4-BQ produced after inoculation of 2-methyl-1,4-HQ with Lac1, Lac2, Dmd and no-enzyme control  Pairwise comparisons of Dmd vs Lac1/2 and control  Pairwise comparisons among Lac1/2 and control  Laccase comparison assay - amount (ng) of 2-methoxy-3-methyl-1,4-BQ produced after inoculation of 2-methoxy-3-methyl-1,4-HQ with Lac1, Lac2, Dmd and no-enzyme control  Pairwise comparisons of Dmd vs Lac1/2 and control  Pairwise comparisons among Lac1/2 and control | 4 groups with 3 replicates each  3 vs 3 (each)  3 vs 3 (each)  4 groups with 3 replicates each  3 vs 3 (each)  3 vs 3 (each)  4 groups with 3 replicates each  3 vs 3 (each)  3 vs 3 (each) | Welch's ANOVA  Tukey HSD tests  Tukey HSD tests  Welch's ANOVA  Tukey HSD tests  Tukey HSD tests  Welch's ANOVA  Tukey HSD tests  Tukey HSD tests | 0.02426  All 3 tests p≤ 0.00016  All 3 tests p≥ 0.9957  0.00339  All 3 tests p≤ 0.0001  All 3 tests p≥ 0.4175  0.00442  All 3 tests p≤ 0.00148  All 3 tests p≥ 0.0553 | F_3,8_= 9.8, N=12  F_3,8_= 29.6, N=12  F_3,8_= 31.3, N=12 |
| 6I | *In vitro* Dmd expression assay - amount (ng) of 1,4-BQ produced after inoculation of 1,4-HQ with Dmd and no-enzyme control  *In vitro* Dmd expression assay - amount (ng) of 2-methyl-1,4-BQ produced after inoculation of 2-methyl-1,4-HQ with Dmd and no-enzyme control  *In vitro* Dmd expression assay - amount (ng) of 2-methoxy-3-methyl-1,4-BQ produced after inoculation of 2-methoxy-3-methyl-1,4-HQ with Dmd and no-enzyme control | 10 Dmd vs 10 control (no enzyme) | Mann-Whitney U tests | 0.00018  0.00018  0.00018 | z= 3.7, N= 20  z= 3.7, N= 20  z= 3.7, N= 20 |
| 6L | Amount (ng) of 1,4-benzoquinone (1,4-BQ) in GFP vs MeOS RNAi beetles  Amount (ng) of 2-methyl-1,4-BQ in GFP vs MeOS RNAi beetles  Amount (ng) of 2-methoxy-3-methyl-1,4-BQ in GFP vs MeOS RNAi beetles | 8 (GFP) vs 4 (RNAi beetles) | Mann-Whitney U tests | 0.07453  0.55221  0.00847 | z= 1.8, N= 12  z= 0.6, N= 12  z= 2.6, N= 12 |
| **Figure 7** | **Comparison** | **N number** | **methods** | **P value** | **statistical values** |
| 7A | Comparison of the survival (%) of beetles previously inject with dsRNA targeting Dmd (BQ-less phenotype), MFASN (solvent-less phenptype) and GFP (control)  Postdoc comparison:  Dmd vs GFP  MFASN vs GFP  MFASN vs Dmd | 3 groups with 14 replicates each  14 groups vs 14 groups  14 groups vs 14 groups  14 groups vs 14 groups | Generalized linear model (GLM) with a binomial distribution  Simultaneous Tests for General Linear Hypotheses using Tukey Contrasts | < 0.0001  <0.001  <0.001  0.339 | Χ^2^_2,39_= 53.5, N= 42  z= -6.9, N= 28  z= -5.5, N= 28  z= -1.4, N= 28 |
| 7B | Comparison of toxicity of *Dalotia* gland chemicals - single compounds and mixtures thereof – against *Drosophila melanogaster* larvae. Number of alive larvae was counted one hour after the exposure to the chemicals  Pairwise comparisons between PBS, esters, C11, BQs and esters+C11 mixture  Pairwise comparisons between esters+BQs mistures, C11+BQs mixture and all-compound mixtures (mimics Dalotia composition)  Pairwise comparisons between PBS, esters, C11, BQs and esters+C11 mixture vs esters+BQs mistures, C11+BQs mixture and all-compound mixtures (mimics Dalotia composition) | 8 groups with 5 replicates of 25 larvae each  5 vs 5 for each test  5 vs 5 for each test  5 vs 5 for each test | One-way ANOVA  Tukey HSD tests  Tukey HSD tests  Tukey HSD tests | < 0.0001  All 10 tests p≥ 0.19  All 3 tests p≥ 0.22  All 15 tests p≤ 0.003 | F_7,32_= 26.7, N= 40 |
| 7C | Bacterial growth as OD_500_ of *P. fluorescens* after 24h inoculation with *Dalotia* gland chemicals  Pairwise comparisons between PBS, single compound classes and two-compound mixtures  Pairwise comparisons between the all-compound mixtures (mimics *Dalotia* composition) and PBS, single compound classes and two-compound mixtures | 8 groups with 13 replicates each  13 vs 13 for each test  13 vs 13 for each test | One-way ANOVA  Tukey HSD tests  Tukey HSD tests | < 0.0001  All 21 tests p≥ 0.18  All 7 tests p≤ 0.007 | F_7,96_= 6.9, N= 103 |
| 7D | Comparison of the surface coating ability (K/σ) of Dalotia gland compounds (compound class, mixtures thereof) and alternative evolutionary scenarios of gland chemistry. Surface coating ability was determined with drip of surface rhenology.  Pairwise comparison across all 8 different test groups (details see raw data files) | 8 groups with a variable number of replicates (see raw data files)  28 pairwise comparisons | Kruskal-Wallis test  Dunn’s pairwise posthoc test (Benjamini-Hochberg correction was used to adjust the level of significance) | < 0.0001  For 11 significant tests p≤ 0.012  For 17 non-significant tests p≥ 0.04 | Χ^2^_7,51_= 42.3, N= 58 |
